# Spleen-derived Small Extracellular Vesicles Protect against Myocardial Infarction via Mediating Spleen-heart Crosstalk

**DOI:** 10.1101/2025.06.16.660039

**Authors:** Keyu Liu, Yue Luo, Weifeng Liao, Riken Chen, Tugang Chen, Haitao Huang, Jinrong Xu, Fang Fang, Qinghua Chen, Dongling Lin, Ting Gu, Burton B Yang, Wenliang Chen, Liangqing Zhang

**Affiliations:** Department of Anesthesiology, the Second Affiliated Hospital of Guangdong Medical University, Guangdong Medical University, Zhanjiang, Guangdong, China; Translational Medicine Center, the Second Affiliated Hospital of Guangdong Medical University, Guangdong Medical University, Zhanjiang, Guangdong, China; Medical Interdisciplinary Science Research Center of Western Guangdong, the Second Affiliated Hospital of Guangdong Medical University, Guangdong Medical University, Zhanjiang, Guangdong, China; Respiratory and Critical Care Medicine, the Second Affiliated Hospital of Guangdong Medical University, Zhanjiang, Guangdong, China; Department of Cardiovascular Medicine, the Second Affiliated Hospital of Guangdong Medical University, Guangdong Medical University, Zhanjiang, Guangdong, China; Office of Academic Research, the Second Affiliated Hospital of Guangdong Medical University, Guangdong Medical University, Zhanjiang, Guangdong, China; Department of Laboratory Medicine and Pathobiology, University of Toronto, 2075 Bayview Ave, Toronto, Ontario, M4N 3M5 Canada

**Keywords:** Acute myocardial infarction, Spleen, Small extracellular vesicles, MPC1, Cardiosplenic axis

## Abstract

**Background and Aims:** Acute myocardial infarction (AMI) triggers systemic responses that influence cardiac injury and repair, but protective mediators within the cardiosplenic axis remain incompletely understood. This study aimed to investigate whether spleen-derived small extracellular vesicles (sEVs) exert cardioprotection after AMI, identify critical cargo, and evaluate their clinical relevance.

**Methods:** The effects of splenectomy, spleen-derived sEVs and pharmacological inhibition of sEVs biogenesis on cardiac injury were evaluated in mice with AMI. Parabiosis, donor splenectomy, and fluorescent labelling traced the origin and myocardial recruitment of circulating sEVs. 4D-proteomics profiled sEVs cargo with a focus on pyruvate carrier-1 (MPC1).

**Results:** Splenectomy worsened survival, ventricular function, infarct size, and fibrosis in mice with AMI. AMI upregulated splenic extracellular-vesicle pathways, with sEVs release peaking at day 3 (M3D-sEVs). Labelled splenic sEVs preferentially accumulated in ischemic myocardium, confirmed by parabiosis experiments. M3D-sEVs improved survival and cardiac function and reduced infarct size, fibrosis, apoptosis, inflammation, and hypertrophy. In addition, anti-apoptotic effects were reproduced in vitro. Inhibition of sEVs biogenesis decreased circulating sEVs, aggravated injury, and was rescued by M3D-sEVs. M3D-sEVs were enriched in MPC1, and MPC1 neutralization or pharmacological blockade abrogated sEVs-mediated restoration of respiration, ATP generation, and reduction of reactive oxygen species. Plasma sEVs-associated MPC1 was highest in patients with AMI, intermediate in coronary heart disease, and lowest in controls.

**Conclusions:** The spleen responds adaptively to AMI by releasing MPC1l⍰enriched sEVs that travel to injured myocardium, preserve mitochondrial energetics, and reduce damage, supporting cargol⍰specific sEVs augmentation and sEVsl⍰MPC1 as a potential therapeutic target and biomarker in ischemic heart disease.

**Structured graphical abstract:** Acute myocardial infarction activates a spleen-heart axis in which the spleen releases mitochondrial pyruvate carrier 1 (MPC1) enriched small extracellular vesicles that home to the infarcted myocardium, preserve mitochondrial oxidative phosphorylation, reduce ROS, and limit infarct size, apoptosis, fibrosis, and hypertrophy. Circulating sEVs-associated MPC1 might serve as a potential biomarker for estimating risk of ischemic heart disease.

**Translational perspective:**

- Our findings reveal a novel heart-spleen communication axis mediated by splenic small extracellular vesicles (sEVs) following AMI. These vesicular messengers predominantly deliver mitochondrial pyruvate carrier 1 (MPC1), enhancing cardiac mitochondrial energy metabolism in the injured myocardium, ultimately improving post-AMI functional recovery in experimental models.
- A higher plasma sEVs-associated MPC1 level was observed in AMI patients when compared with non-AMI patients. These findings support the clinical relevance of the spleen-heart axis, introduce sEVs-associated MPC1 as a potential circulating biomarker for early myocardial injury and therapeutic target.

## Introduction

Acute myocardial infarction (AMI) results from sudden interruption of coronary blood flow, leading to ischemia and subsequent injury to the myocardium. Despite major advancements in early reperfusion therapy, AMI remains a leading cause of morbidity and mortality worldwide^1^. A deeper understanding of the molecular mechanisms involved in AMI pathogenesis and an identification of novel therapeutic targets are still urgently needed.

The concept of spleen-heart crosstalk, termed the cardiosplenic axis, was first described in 1949, when splenic nerves activation by electric stimulation improves ventricle function in dogs^2^. Since then, the spleen has been implicated in cardioprotection, potentially through its regulation of inflammatory processes after myocardial injury by mobilizing immune cell populations such as monocytes and by releasing signaling factors that influence downstream effects within the heart^3,4^. For example, stimulation of splenic activity via the vagus nerve during remote ischemic conditioning was shown to reduce cardiac infarct size in a rat model of AMI^5^, suggesting that spleen-derived factors may play a critical role in modulating ischemic cardiac injury. However, the precise molecular mechanisms underlying heart-spleen communication remain to be fully elucidated.

Small extracellular vesicles (sEVs), including exosomes, have been proposed as critical mediators of intercellular and inter-organ signaling^6^, particularly in the context of cardiovascular injury^7^. sEVs are nanosized vesicles (30-150nm) that carry a diverse range of bioactive molecular cargo, including proteins, lipids, and nucleic acids. These cargos are shuttled systemically to recipient cells to regulate their function. The roles of sEVs in mediating cellular and organ-level communications make them promising candidates for use as disease biomarkers, therapeutic agents, or drug delivery vehicles^8^. Recent studies have shown that plasma sEVs derived from AMI patients promote angiogenesis, regulate ferroptosis and autophagy, thereby alleviating hypoxic myocardial injuries^9–11^. It has also been reported that the majority of plasma sEVs in AMI patients originate from the spleen^12^, indicating that spleen-derived sEVs may play critical roles in the pathophysiology of AMI.

Ischemic injury triggers a cascade of pathological responses, including inflammation, metabolic dysfunction, and cell death, which ultimately contribute to adverse cardiac remodeling and progression to heart failure^13^. Among these processes, mitochondrial dysfunction plays a central role in AMI, as the heart’s high energy demands are heavily dependent on mitochondrial oxidative metabolism^14^. Ischemia-induced impairment of mitochondrial function compromises ATP production, generates excessive reactive oxygen species (ROS), and promotes cardiomyocyte apoptosis, further exacerbating myocardial injury and impairing functional recovery^15^. Mitochondrial Pyruvate Carrier 1 (MPC1), a key subunit of the mitochondrial pyruvate carrier (MPC) complex, has gained attention as a critical regulator of mitochondrial bioenergetics. The MPC complex, composed of MPC1 and MPC2, facilitates transport of pyruvate from the cytosol into the mitochondrial matrix, enabling its subsequent entry into the tricarboxylic acid (TCA) cycle and oxidative phosphorylation^16^. Pyruvate metabolism is essential for efficient energy production, particularly in cardiomyocytes, which has a limited capacity to adapt to metabolic stress^17^. Dysregulation of MPC1 expression or function in ischemic conditions disrupts this process, leading to cytosolic pyruvate accumulation, diminished TCA cycle activity, and a shift toward less efficient anaerobic glycolysis^18^. Restoration of MPC1 expression during ischemic injury has been shown to enhance mitochondrial function, reduce oxidative stress, and facilitate cardiomyocyte survival, making MPC1 a promising therapeutic target for metabolic management of AMI^19^.

In this study, we investigated the crosstalk between spleen and heart during AMI, focusing on the potential role of spleen-derived sEVs in delivering MPC1 to the ischemic heart and preserving mitochondrial function in cardiomyocytes. Elucidating these mechanisms may offer valuable insights into systemic regulation of mitochondrial bioenergetics and identify novel therapeutic strategies to enhance myocardial repair and recovery following ischemic injury.

## Methods

The comprehensive descriptions of experimental procedures, protocols, and reagents used in this study are provided in the Supplementary Material.

## Results

### Splenectomy aggravates ischemic injury and adverse remodeling in AMI mice

We first assessed splenic immune activity after myocardial ischemia. On day 3 postl⍰AMI, both heart and spleen weights were increased versus sham (Figure S1A-S1C). The white pulp, which is critical for the initiation of adaptive immune responses in spleen^20,21^, was significantly enlarged in AMI mice, as demonstrated by H&E staining; the marginal zone was also expanded relative to sham mice (Figure S1D-S1F). Spleens from AMI mice showed increased mRNA expression of inflammatory cytokines (TNFl⍰α, ILl⍰10, ILl⍰6, and IFNl⍰γ), peaking at day 3 (Figure S1G-S1J), indicating an obvious splenic response to injury after AMI.

Previous studies have reported variable effects of splenectomy on cardiac injury^22–24^. To clarify the impact of spleen removal, we performed splenectomy immediately following ligation of the left anterior descending (LAD) coronary artery. Compared with AMI mice with intact spleens, splenectomized AMI mice exhibited significantly poorer survival rates, reduced left ventricular ejection fraction and fractional shortening (Figure 1A-1D), larger cardiac infarct size (Figure 1E-1F), and higher serum creatine kinase-MB (CK-MB) levels (Figure S2A). Masson’s trichrome staining showed that splenectomy also increased cardiac collagen deposition (Figure 1G-1H), and elevated expression of fibrosis-related proteins, such as α smooth muscle actin (α-SMA), collagen I (Col), and fibronectin (FN) (Figure 1I-1J), indicating that splenectomy increased cardiac fibrosis post AMI. Furthermore, splenectomized mice also exhibited an increased TUNEL positive rate (Figure 1K-1L), with higher Bax and cleaved caspase-3 (C-Cas3) and lower Bcl2 expression in ischemic hearts at day 3 (Figure 1M-1N). Standardized short-axis, mid-ventricular transverse sections stained with wheat germ agglutinin (WGA), cardiac troponin I and nuclear counterstain (DAPI), confirmed aggravated myocyte hypertrophy in splenectomized AMI mice at day 28 (Figure 1O-1P). H&E staining revealed that splenectomy worsened myocardial structural disarray in AMI mice (Figure S2B). These data support a protective role of the spleen in limiting ischemic injury and adverse remodeling.

**Figure 1.**
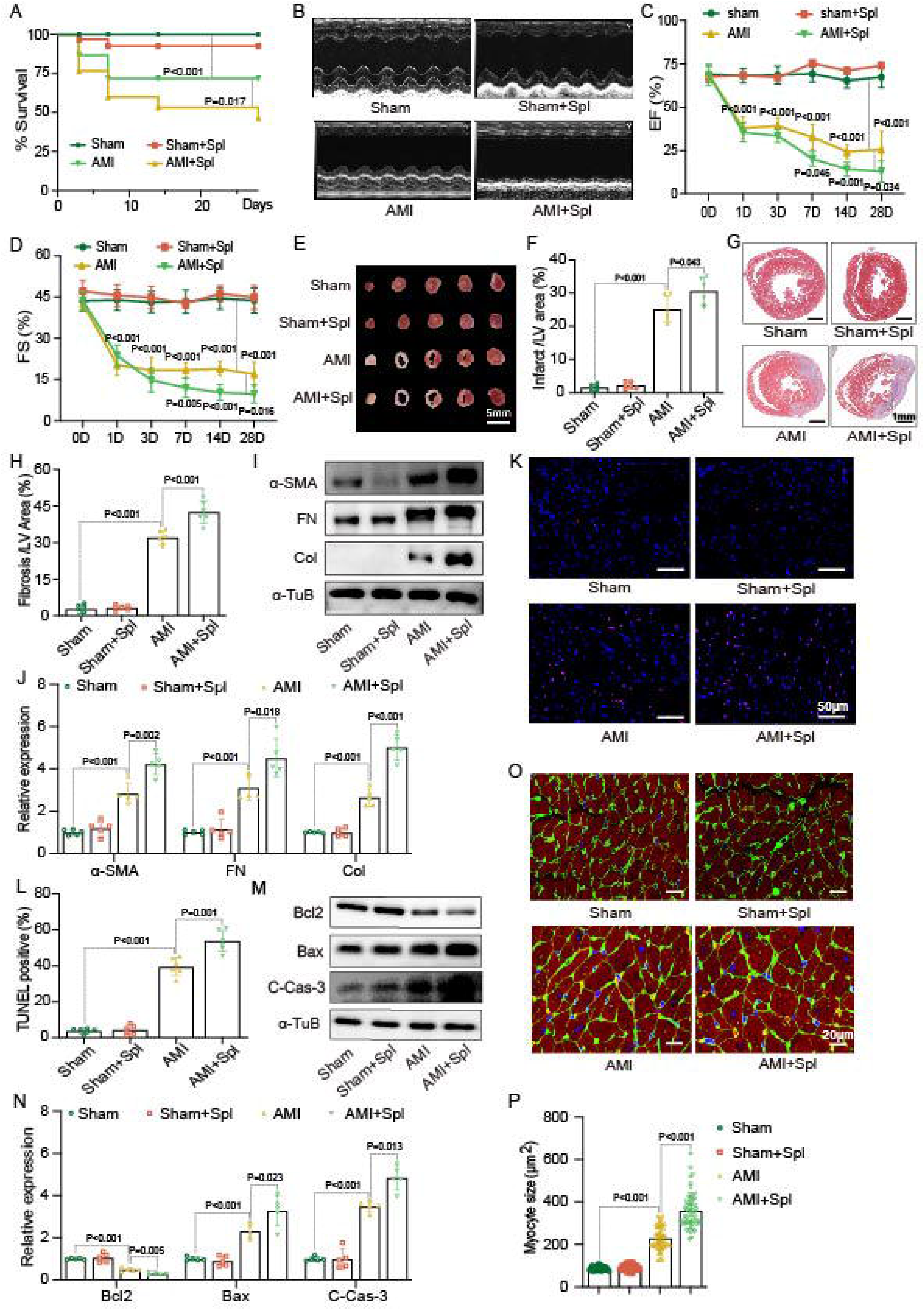
Splenectomy aggravates cardiac injury in AMI mice. **A**, Twenty-eight-day Kaplan-Meier survival curves for sham operation (Sham) or AMI mice, with or without splenectomy (n=30 mice/group). **B**, Representative echocardiograms from sham, splenectomized sham (Sham+Spl), AMI, and splenectomized AMI (AMI+Spl) mice 7 days post AMI (n=6/group). **C** and **D**, Summary of left ventricle EF (EF%) and FS (FS%) in Sham, Sham+Spl, AMI, and AMI+Spl mice at indicated time-points (n=6/group). Detailed echo parameters at day 7 post AMI were shown in Table S6. **E** and **F**, Representative TTC images of cross-sectional cardiac slices and quantitation of infarct size in hearts from Sham, Sham+Spl, AMI, and AMI+Spl mice 1 day post AMI (n=6/group). **G** and **H,** Representative images of Masson’s trichrome staining and quantitation of the fibrosis area in hearts from Sham, Sham+Spl, AMI, and AMI+Spl mice 14 days after AMI (n=6/group). **I**-**J**, Representative western blots quantitation of collagen I, fibronectin, and α-SMA expression in mice hearts 14 days post MI (n=6/group). **K**-**L,** Representative TUNEL images and quantitation of TUNEL-positive rates in myocardial tissues from Sham, Sham+Spl, AMI, and AMI+Spl mice 3 days post AMI (n=6/group). **M**-**N,** Representative western blots and quantitation of C-Cas-3, Bax, and Bcl2 in mice hearts 3 days post AMI (n=6/group). **O**-**P**, Representative WGA staining images of cardiac sections and quantitation of cardiomyocyte size in mice hearts 28 days post AMI (n=6/group). Statistical analyses were performed using mixed-effects ANOVA followed by the Sidak multiple comparisons test (**C** and **D**), the one-way ANOVA with the Dunnett multiple comparisons test (**H**, **P** and **L**), the Brown-Forsythe and Welch ANOVA test with the Dunnett T3 multiple comparisons test (**F**, **J**; Collagen I and N; Bcl2, C-Cas-3), the one-way ANOVA with the Tukey multiple comparisons test (**J**; α-SMA and N; Bax), the Kruskal Wallis with the Dunn multiple comparisons test (**J**; Fibronectin). Survival curve rate was analyzed by the Kaplan-Meier method and compared by the log-rank test in **A**. Spl indicates splenectomy; EF, ejection fraction; FS, fractional shortening; LV, left ventricular; FN, Fibronectin; Col, Collagen I; α-TuB, α-Tubulin.

### Extracellular vesicles-related gene expression is upregulated in the spleen of AMI mice

To investigate transcriptomics alteration in the spleen post AMI, we performed RNA-Seq analysis of spleen tissues (Figure 2A). A total of 2621 differentially expressed genes (DEGs) were identified in spleens from AMI mice versus sham, with 1129 genes upregulated and 1492 genes downregulated (Figure 2B and Figure S3A). Gene Ontology (GO), KEGG, and Gene set enrichment analysis (GSEA) analysis showed that the significant enrichment of terms was related to extracellular vesicles and exosomes (Figure S3B-S3G). The biogenesis processes of sEVs are intricate and can be affected by multiple pathways^25^, such as soluble N-ethylmaleimide-sensitive fusion attachment protein receptor (SNARE), Neutral sphingomyelin phosphodiesterase 3 (Smpd3), and Rab GTPases (e.g., Rab27a/Rab27b, Rab11). Quantitative PCR validated that the key genes involved in sEVs formation and trafficking were upregulated in the spleens of AMI mice versus sham, including Rab27a, Rab31, Rab11, Pdcd6ip, Rab22a, and Smpd3 (Figure 2C). These findings indicate that myocardial ischemia might activate extracellular vesicles biogenesis in the spleen, prompting further investigation into the role and function of spleen-derived sEVs in AMI.

**Figure 2.**
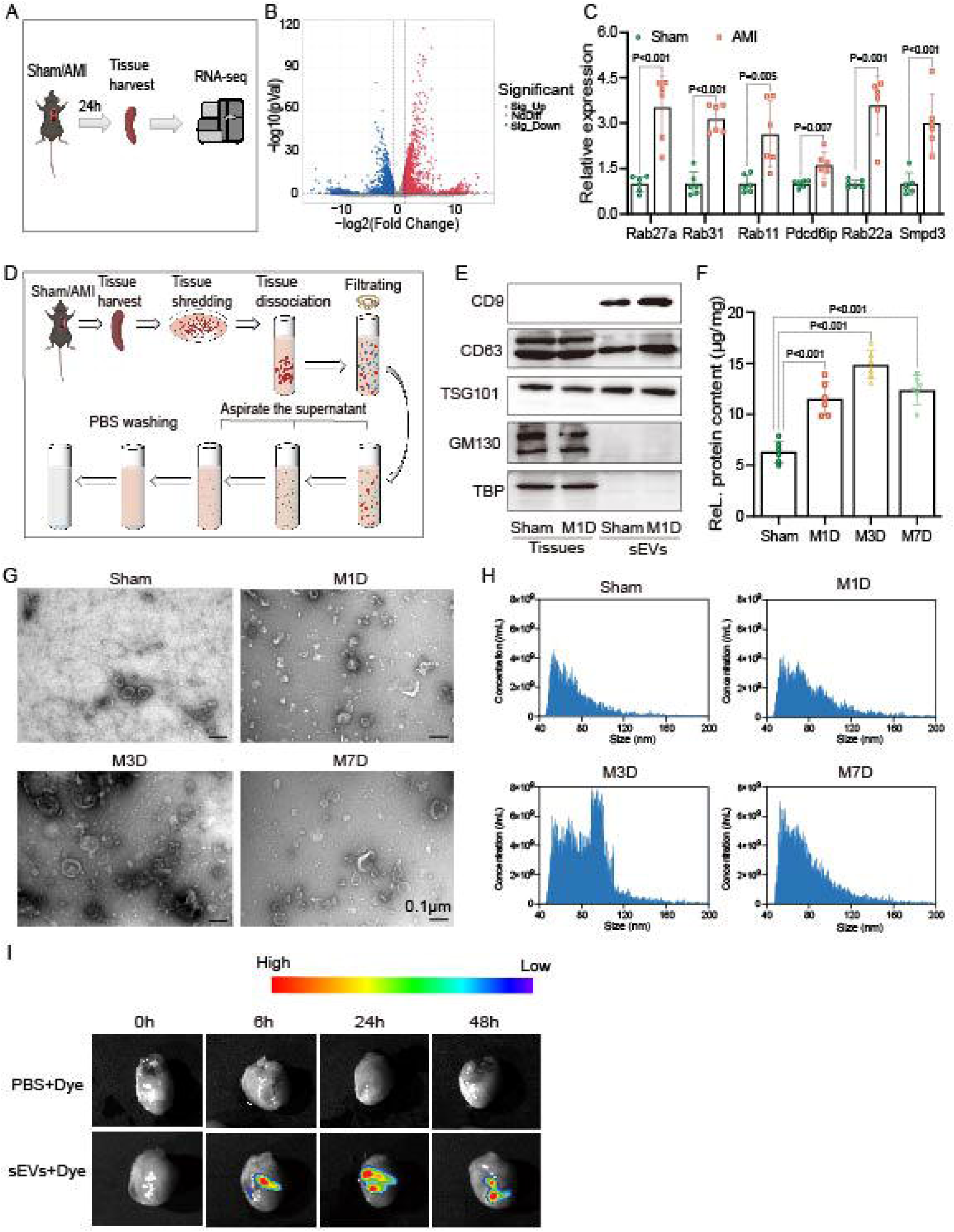
Spleen-derived sEVs biogenesis is enhanced in response to AMI and these sEVs were enriched in ischemic hearts in mice. **A,** Schematic diagram of RNA-Seq for spleens from Sham and AMI mice. **B**, Volcano plot of differentially expressed genes (DEGs) of spleens from sham and AMI mice. **C**, sEVs biogenesis-related DEGs expression in spleen tissues from sham and AMI mice were validated using qPCR 1 day post AMI. Data were normalized to GAPDH and expressed as folds of Sham group (n=6/group). **D,** The flowchart of spleen-derived sEVs extraction. **E**, Representative western blot images of sEVs markers. **F**, Relative (ReL.) protein contents of sEVs (n=6/group). **G**, Representative TEM images of spleen-derived sEVs from various mice groups as indicated. **H**, Size distributions of spleen-derived sEVs measured by nFCM. **I**, The residence of DiR-labeled spleen-derived sEVs in myocardium after intramyocardial injection. Representative heart images were shown. Statistical analyses were performed by unpaired t test (**C**; Rab27a, Rab11, Pdcd6ip and Smpd3), the Mann-Whitney U test (**C**; Rab31 and Rab22a) and the one-way ANOVA with the Dunnett multiple comparisons test (**F**). M1D indicates day 1 after AMI; M3D, day 3 after AMI; M7D, day 7 after AMI.

### Spleen-derived small extracellular vesicles biogenesis and trafficking to injured myocardium increase after AMI

sEVs were isolated from mouse spleen using our established protocol^26^ (Figure 2D). The physical characteristics of the isolated sEVs were validated by transmission electron microscopy (TEM), western blotting, and nano-flow cytometry (nFCM). The isolated particles expressed conventional sEVs markers (Tumor susceptibility gene 101, TSG101; CD63; and CD9) but not Golgi matrix protein 130 (GM130) and TATA box binding protein (TBP) (Figure 2E), displayed typical cupl⍰shaped morphology on TEM and a size range of 30-150 nm by nFCM (Figure 2G-2H), indicating high-purity sEVs preparations. Both protein contents and particle number of spleen-derived sEVs increased after AMI, peaking at day 3 (Figure 2F, Figure S4A).

For trafficking studies, spleen-derived sEVs were labelled with Dimethyl red dye (DiR) and purified by iodixanol density gradient ultracentrifugation to remove unbound dye/aggregates; PBS+dye controls were processed identically. Near-infrared fluorescence imaging after intraperitoneal injection showed preferential early accumulation of labelled spleen sEVs in ischemic hearts (Figure S4B). These sEVs could reside in the hearts of recipient AMI mice post intramyocardial injection for at least 48 h, with lower signal in liver, lung, spleen, and kidney (Figure 2I, S4C and S4D), and later redistribution to liver and kidneys (Figure S4D). PBS+dye controls showed negligible signals (Figure 2I, S4B-S4D). These findings indicate that splenic sEVs production is increased after AMI, particularly at day 3, and that spleenl⍰derived sEVs are recruited to injured myocardium.

### Spleen-derived sEVs at day 3 after AMI attenuate myocardial injury, apoptosis, inflammation, and hypertrophy

To define the timel⍰dependent effects of splenic sEVs, AMI recipient mice received intramyocardial injections of sEVs isolated from spleens of sham, dayl⍰1 (M1D), dayl⍰3 (M3D), or dayl⍰7 (M7D) AMI donors. Among all groups, M3D-sEVs exerted the greatest benefit, improving survival (Figure 3A), left ventricular ejection fraction and fractional shortening (Figure 3B-3D), reducing infarct size (Figure 3E-3F), and limiting collagen accumulation (Figure 3G and 3H) and fibrotic protein expression (α-SMA, Col, and FN) (Figure 3I and 3J) compared with vehicle, sham sEVs or M1D/M7D-sEVs. M3D-sEVs treatment improved myocardial structural morphology (Figure S5A), decreased TUNEL positivity rate (Figure S5B and S5C), lowered Bax and cleaved caspase-3 (C-Cas-3) expression, and increased Bcl2 expression (Figure 3K-3L). Besides, M3D-sEVs reduced serum CK-MB levels (Figure S5D).

**Figure 3.**
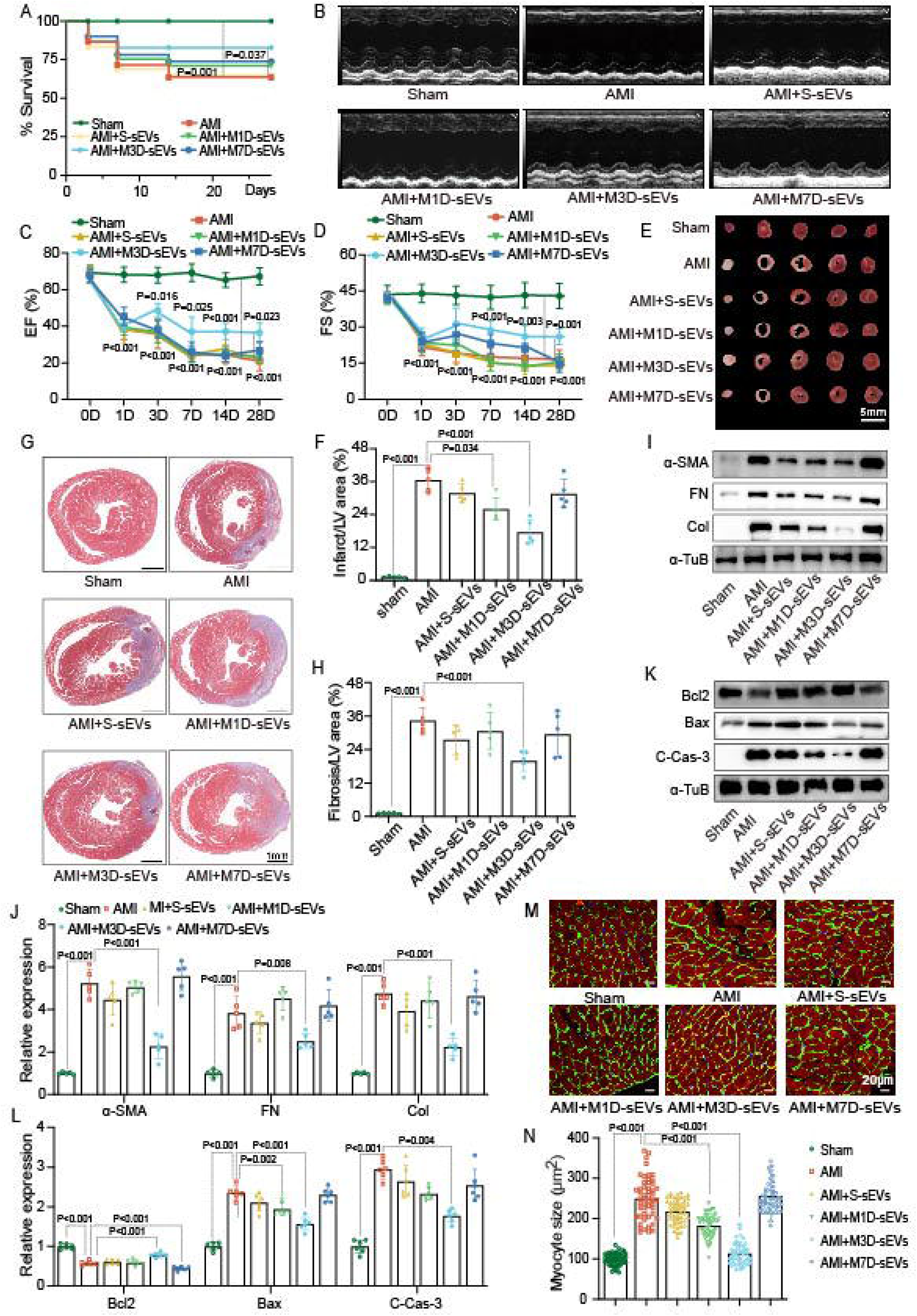
Spleen-derived sEVs from AMI mice alleviate ischemia injuries in hearts and cardiomyocytes. **A**, Twenty-eight-day Kaplan-Meier survival curves for sham or AMI mice treated with indicated sEVs or vehicle (n=30 mice per group). **B**, Representative echocardiograms of sham or AMI mice treated with indicated sEVs or vehicle 7 days after AMI (n=6/group). **C-D**, Quantitation of EF% and FS%. Deteiled echo parameters at day 7 post AMI were shown in Table S7. **E-F**, Representative images of TTC staining and quantitation of infarct area in mice hearts 1 day after AMI (n=6/group). **G-H**, Representative images of Masson’s trichrome and quantitation of the fibrotic area in mice hearts 14 days after AMI (n=6/group). **I-J**, Representative western blot images and quantitation of cardiac collagen I, Fibronectin, and α-SMA 14 days after AMI (n=6 mice per group). **K-L**, Representative western blot images and quantitation of cardiac C-Cas-3, Bax, and Bcl2 3 days after AMI (n=6/group). **M-N**, Representative WGA staining images of cardiac sections and quantitation of cardiomyocytes 28 days after AMI (n=6/group). Statistical analyses were performed using mixed-effects ANOVA followed by the Sidak multiple comparisons test (**C** and **D**), the one-way ANOVA with the Dunnett multiple comparisons test (**F** and **H**), the one-way ANOVA with the Tukey multiple comparisons test (**K**; α-SMA, Fibronectin and **L**; Bcl2, C-Cas-3), the Kruskal-Wallis with the Dunn multiple comparisons test (**K**; Collagen I and **L**; Bax), and the Brown-Forsythe and Welch-ANOVA test with the Dunnett T3 multiple comparisons test (**N**). Survival curve rate was analyzed by the Kaplan-Meier method and compared by the log-rank test in **A**. S-sEVs indicates small extracellular vesicles derived from spleens of sham mice; M1D-sEVs, M3D-sEVs and M7D-sEVs indicate sEVs derived from spleens of AMI mice 1, 3, and 7 day post AMI, respectively; EF, ejection fraction; FS, fractional shortening; LV, left ventricular; FN, Fibronectin; Col, Collagen I; and α-TuB, α-Tubulin.

Transverse sections with triple staining of WGA+cardiac troponin I+DAPI showed that M3D-sEVs attenuated cardiomyocyte hypertrophy (Figure 3M-3N), which was further supported by qPCR results showing reduced ANP and BNP expression (Figure S6A and S6B). In parallel, M3D-sEVs reduced IL-1β, IL-6, and TNF-α mRNA expression in infarcted myocardium (Figure S6C-S6E). In vitro, PKH26l⍰labelled, ultracentrifugationl⍰purified M3Dl⍰sEVs (with PBS+dye group as controls) were efficiently taken up by H9C2 cells (Figure S7A). In OGD-challenged cells, M3D-sEVs increased Bcl2 and decreased Bax and C-Cas-3 (Figure S7B-S7C). Collectively, these data identify M3D-sEVs as the most cardioprotective subset and they were used for subsequent mechanistic studies.

### Parabiosis demonstrates splenic origin and recruitment of circulating sEVs to the heart

To evaluate whether splenic sEVs contribute to the circulating pool reaching injured myocardium, we established parabiosis between CD45.1 and CD45.2 mice and induced nonl⍰reperfused AMI in the host after 4 weeks of shared circulation (Figure 4A). Stable donor chimerism of CD45^+^ leukocytes in both intact and splenectomized host mice was confirmed by FACS analysis of peripheral blood at 3 days post-AMI (Figure 4B-4C). In hosts subjected to AMI plus splenectomy, the donor spleen showed compensatory enlargement and increased sEVs release (Figure S8A-S8F), and host cardiac function, infarct size and fibrosis were comparable with hosts retaining their spleen (Figure 4D-4K), indicating functional compensation by the donor spleen.

**Figure 4.**
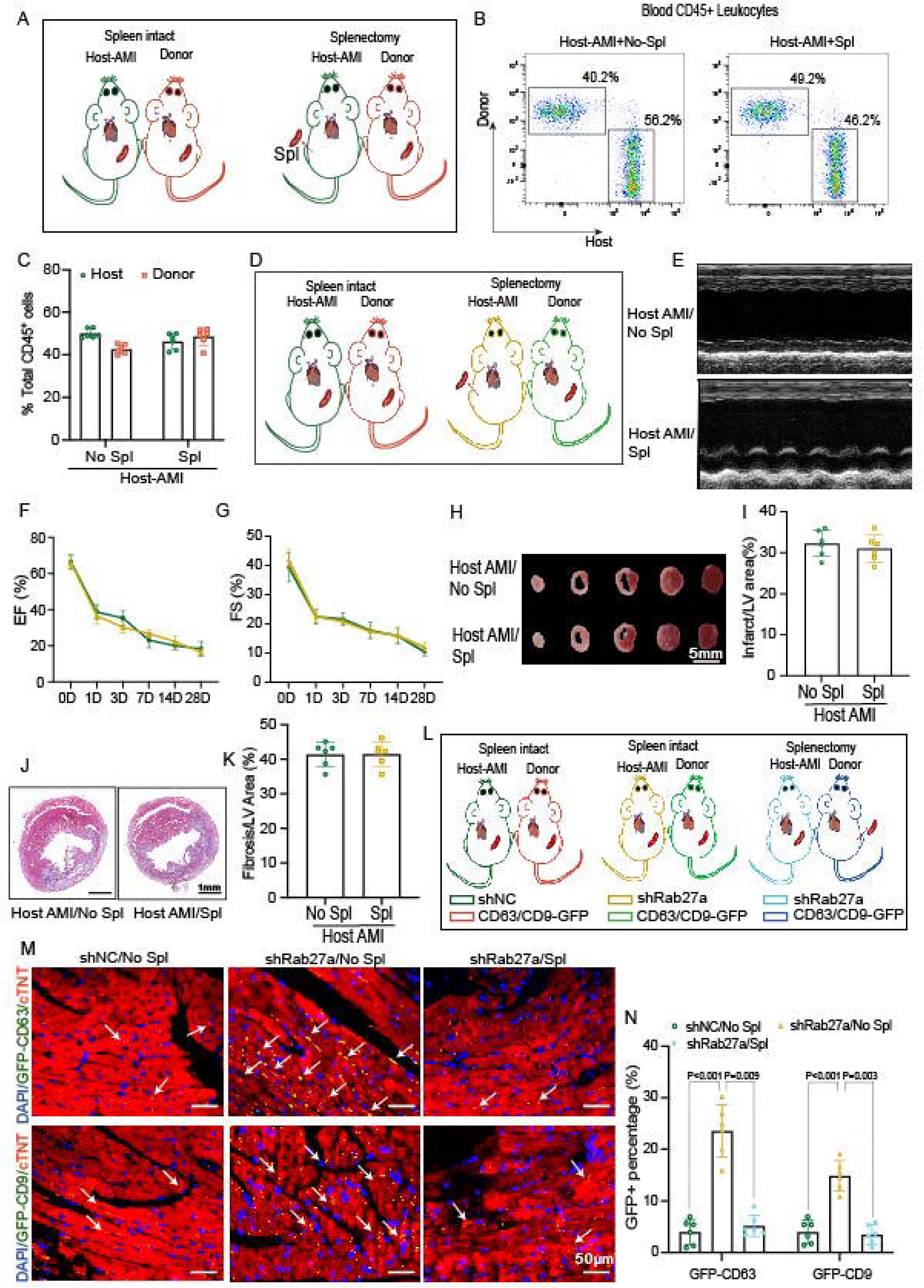
sEVs release after AMI in mice depends on the spleen. **A,** Parabiosis schema joining cluster of different (CD) 45 isotype-mismatched host (spleen intact or after splenectomy) and donor parabiont mice, with AMI induced in the host. **B,** Fluorescence-activated cell sorting plots and quantitation of donor chimerism in host mouse blood (total CD45^+^ leukocytes) in spleen-intact and splenectomized host 3 days after AMI (n=6/group). **C,** Quantitation of fluorescence-activated cell sorting plots. **D,** Parabiosis schema joining cluster of host (spleen intact or after splenectomy) and donor parabiont mice, with AMI induced in the host. **E,** Representative echocardiograms of host mice in 7 days post AMI. **F-G**, Quantitation of EF% and FS% (n=5-6/group). Detailed echo parameters at day 7 post AMI were shown in Table S8. **H-I**, Representative images of TTC staining of cardiac slices from host mice and quantitation of infarct area in mice hearts 1 day after AMI (n=6/group). **J-K,** Representative images of Masson’s trichrome and quantitation of the scar area in mice hearts 28 days after AMI (n=6/group). **L,** Parabiosis schema joining AMI host mice and either spleen-intact or splenectomized donor parabionts, with host mice given AAV2-CD45.2-shRab27a or AAV2-CD45.2-shNC to inhibit sEVs release and donor mice given AAV2-CD45.1-CD9-GFP or AAV2-CD45.1-CD63-GFP to label the sEVs of immune cells. **M-N,** Representative Immunofluorescence images and quantitation of CD9-GFP^+^ or CD63-GFP^+^ rates in myocardial tissues of host mice 3 days after AMI (n=6/group). Statistical analyses were performed using mixed-effects ANOVA followed by the Sidak multiple comparisons test (**F-G**), the Mann-Whitney U test (**I** and **K**), the two-way ANOVA with the Tukey multiple comparisons test (**C**) and the Brown-Forsythe and Welch-ANOVA test with the Dunnett T3 multiple comparisons test (**N**).

To reduce host sEVs biogenesis and secretion, we delivered AAV2l⍰CD45.2l⍰shRab27a to host immune cells, while donor immune cell sEVs were labelled with CD9/CD63l⍰GFP (AAV2l⍰CD45.1l⍰CD9/CD63l⍰GFP) (Figure 4L). With host sEVs biogenesis suppressed, donor-derived GFP^+^ sEVs were readily detectable and increased in host myocardium compared with the controls (normalized to DAPI^+^ nuclei; Figure 4M-4N). Donor splenectomy significantly reduced the number of donorl⍰derived GFP^+^ sEVs in host hearts under the same host shRab27a background (Figure 4M-4N), establishing that a substantial component of the augmented sEVs recruitment to injured myocardium originates from the spleen.

### Pharmacological suppression of neutral sphingomyelinase–dependent of sEVs biogenesis worsens post**11**AMI outcomes and is rescued by M3D**11**sEVs

To examine the functional relevance of circulating sEVs, we inhibited neutral sphingomyelinase with GW4869. A single intraperitoneal dose of GW4869 (GW4869-S) improved left ventricle systolic function (Figure S9A-S9B) and modestly increased shortl⍰term, without improving survival (Figure S9C).

A multiplel⍰dose regimen (2.5 mg/kg/day for three days; GW4869l⍰M) did not impair cardiac structure or function in nonl⍰AMI mice (Figure S10A-S10I). In AMI mice, however, GW4869l⍰M worsened survival, reduced EF and FS in the left ventricle, and increased infarct size, collagen deposition, fibrotic protein expression, CKl⍰MB, structural disarray, cardiomyocyte apoptosis, and hypertrophy (Figure 5A-5O, Figure S12A-S12B). Timel⍰course analyses showed that GW4869l⍰S suppressed splenic sEVs release for ∼24h only, with a rebound to, and later above, vehicle levels by day 3 postl⍰AMI, whereas GW4869l⍰M sustained suppression at all evaluated time points (Figure S11A-S11E), thereby effectively inhibiting the M3D-sEVs peak. Importantly, col⍰administration of M3Dl⍰sEVs, but not sham sEVs, rescued the adverse effects of GW4869l⍰M on cardiac structure and function (Figure 5A-5O, Figure S12A-S12B). These findings support an endogenously protective role of spleenl⍰derived sEVs in limiting postl⍰AMI injury and remodeling.

**Figure 5.**
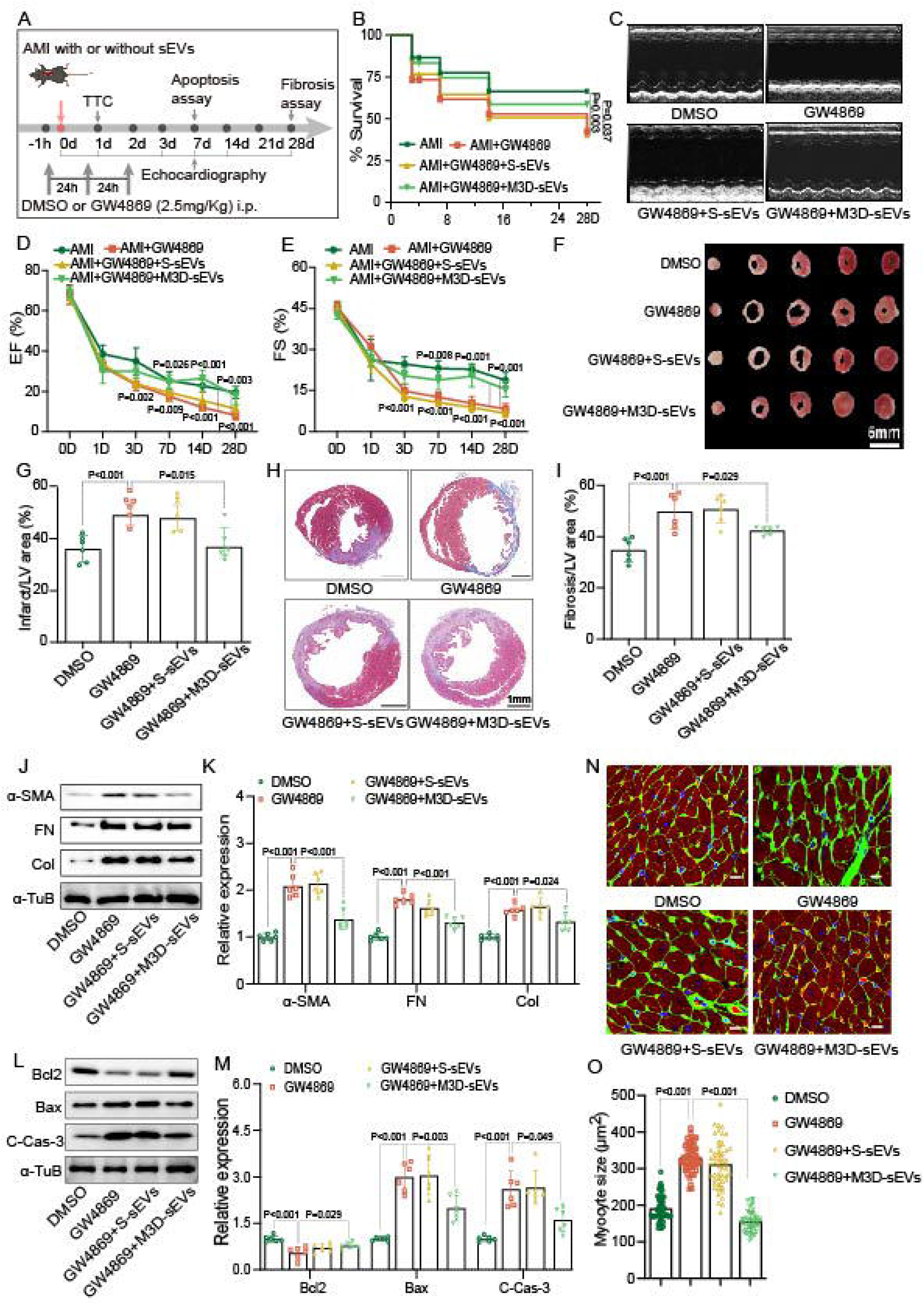
sEVs depletion aggravates cardiac ischemic injury in AMI mice. **A**, Schematic diagram of experimental design for sEVs depletion. **B**, Twenty-eight-day Kaplan-Meier survival curves for AMI mice treated with indicated conditions (n=30 mice per group). **C**, Representative echocardiograms of AMI mice treated with GW4869 plus S-sEVs or M3D-sEVs, or corresponding vehicles 7 days after AMI (n=6 mice per group). **D-E**, Quantitation of EF% and FS%. Detailed echo parameters at day 7 post AMI were shown in Table S8**. F-G**, Representative images of TTC staining of cardiac slices from AMI mice treated with GW4869 plus S-sEVs or M3D-sEVs, or corresponding vehicles, and quantitation of infarct area in mice hearts 1 day after AMI (n=6/group). **J-K**, Representative western blot images and quantitation of cardiac collagen I, fibronectin, and α-SMA 14 days after AMI in mice treated as indicated conditions (n=6/group). **L-M**, Representative western blot images and quantitation of cardiac C-Cas-3, Bax, and Bcl2 of hearts 3 days after AMI in mice (n=6/group). **N-O**, Representative WGA staining images of cardiac sections and quantitation of cardiomyocytes 28 days after AMI (n=6/group). Statistical analyses were performed using mixed-effects ANOVA followed by the Sidak multiple comparisons test (**D** and **E**), the one-way ANOVA with the Dunnett multiple comparisons test (**G**), the Brown-Forsythe and Welch-ANOVA test with the Dunnett T3 multiple comparisons test (**I** and **M**; Bax), the Kruskal-Wallis with the Dunn multiple comparisons test (**M**; C-Cas-3 and **O**), and the one-way ANOVA with the Tukey multiple comparisons test (**K** and **M**; Bcl2). Survival curve rate was analyzed by the Kaplan-Meier method and compared by the log-rank test in **B**. EF, ejection fraction; FS, fractional shortening; LV, left ventricular; FN, fibronectin; Col, Collagen I; and α-TuB, α-Tubulin.

### Spleen-derived M3D-sEVs preserve mitochondrial energetics in ischemic myocardium

To explore the mechanisms underlying the cardioprotective effects of M3D-sEVs, we performed 4D-proteomic profiling to compare the protein cargo of spleen-derived sEVs from sham and AMI mice (Figure 6A). A total of 1068 differentially expressed proteins (DEPs) were identified (M3D-sEVs vs S-sEVs), with 389 up-regulated and 679 down-regulated proteins in M3D-sEVs relative to S-sEVs (Figure 6B). GO enrichment analysis showed that these DEPs are associated with metabolic process, transport, intracellular organelle, electron transfer activity, and structural molecular activity (Figure S13A-S13C). KEGG pathway analysis demonstrated that they were mainly enriched in oxidative phosphorylation, metabolic pathways, and cardiac muscle contraction (Figure 6C). These results suggest that mitochondrial metabolism-related oxidative phosphorylation (OXPHOS) may serve as a key mechanism by which M3D-sEVs alleviate injury in AMI. In infarcted myocardium and OGD-challenged cardiomyocytes, M3D-sEVs restored mRNA expression of mitochondrial electron transport chain (ETC) subunits mRNA expression relative to vehicle or S-sEVs, including Ndufs1, Ndufs2, Sdhc, Sdhb, Uqcrfs1, Uqcrc1, Cox4il, Cox7b, Atp5po, and Atp5f1b (Figure 6D-6E). M3D-sEVs treatment improved mitochondrial ultrastructure in AMI hearts, restoring organized cristae, preserving myofibril integrity, and normali⍰ng mitochondrial size (Figure 6F-6G).

**Figure 6.**
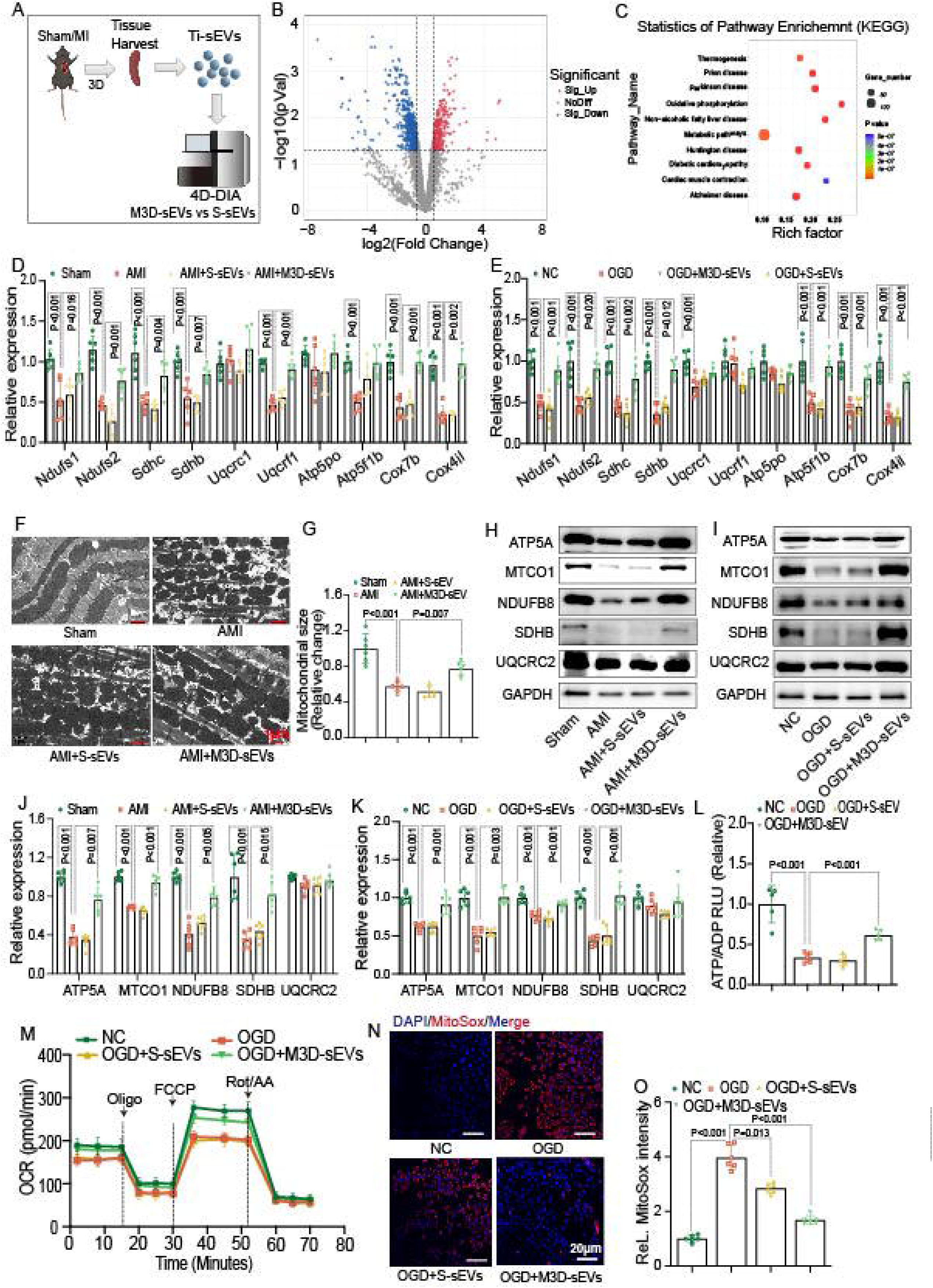
Spleen-derived sEVs from AMI mice improve myocardial mitochondrial energy metabolism. **A,** The flowchart of proteomics experiment for spleens from sham and AMI mouse. **B**, Volcano plot showing the differentially expressed proteins in spleens between sham and AMI mouse. **C,** KEGG enrichment analysis of differentially expressed proteins. **D-E,** mRNA expressions of ETC-related genes in heart tissues from sham and AMI mice treated with S-sEVs, M3D-sEVs, or vehicles for 7 days, and in normal control cells (NC) and OGD-insulted H9C2 cells treated with S-sEVs, M3D-sEVs, or vehicles for 12h (n=6/group). **F-G,** Quantitation of mitochondrial size in these TEM images as indicated groups, being presented as relative value to the sham group (n=6/group). **H-K**, Representative western blotting images and quantitation of ETC complex subunits expression in heart tissues from sham and AMI mice treated with S-sEVs, M3D-sEVs, or vehicles for 7 days, and in normal control cells (NC) and OGD-insulted H9C2 cells treated with S-sEVs, M3D-sEVs, or vehicles for 12h (n=6/group).**L**, ATP/ADP ratios of NC and OGD-insulted H9C2 cells treated S-sEVs, M3D-sEVs, or vehicles for 24h (n=6/group). **M**, Real-time monitoring OCR in H9C2 cells treated NC and OGD-insulted H9C2 cells treated S-sEVs, M3D-sEVs, or vehicles for 12h (n=6/group). **N-O**, Representative MitoSOX staining images and quantitation of the MitoSOX fluorescent intensity in NC and OGD-insulted H9C2 cells treated S-sEVs, M3D-sEVs, or vehicles for 12h (n=6/group). Statistical analyses were performed by the one-way ANOVA with the Dunnett multiple comparisons test (**G** and **L**), the one-way ANOVA with the Tukey multiple comparisons test (**J**; ATP5A, MTCO1, NDUFB8, UQCRC2, **K**;NDUFB8, **D**; Sdhb, Uqcrc1, Atp5po and **E**; Sdhb, Ndufs2, Cox4il), the Brown-Forsythe and Welch-ANOVA test with the Dunnett T3 multiple comparisons test (**D**; Sdhb, **E**; Sdhc, Cox7b, Atp5po, Atp5f1b and **O**), and the Kruskal-Wallis with the Dunn multiple comparisons test (**K**; ATP5A, MTCO1, SDHB, UQCRC2, **D**; Sdhb, Uqcrc1, Atp5f1b and **F**; Ndufs1).

To further elucidate the impact of spleen-derived sEVs in mitochondrial function, we assessed the expression of proteins associated with the ETC complexes in AMI mice and OGD-insulted cells. Immunoblot analysis demonstrated that protein levels of mitochondrial complex I (NDUFB8), complex II (SDHB), complex IV (MTCO1), and complex V (ATP5A) were significantly reduced in both animal and cellular models, while they were restored by M3D-sEVs treatment; in contrast, expression of complex III (UQCRC2) remained unchanged across experimental groups (Figure 6H-6K). In vitro, M3Dl⍰sEVs increased ATP/ADP ratio, enhanced oxygen consumption, and reduced mitochondrial superoxide generation after OGD (Figure 6L-6O). Collectively, these data suggest that M3Dl⍰sEVs mitigate mitochondrial injury and preserve bioenergetics in ischemic myocardium.

### MPC1 is enriched in M3D**11**sEVs and elevated in plasma sEVs from patients with AMI

To identify critical cardioprotective mediators within M3D-sEVs, we crossl⍰referenced spleen sEVs and plasma proteomes (Figure S14) from sham and AMI mice, identifying a small set of consistent regulated proteins (21 upregulated and 12 downregulated) (Figure 7A). Among mitochondrial energyl⍰related candidates (Table S4), mitochondrial pyruvate carrier 1 (MPC1) and phosphoenolpyruvate carboxykinase 2 (PCK2) were selected for validation. MPC1, but not PCK2, was markedly enriched in M3Dl⍰sEVs versus Sl⍰sEVs (Figure 7B-7C). Furthermore, when comparing sEVs from multiple organs post-AMI, increased MPC1 in sEVs was observed only in spleenl⍰derived sEVs after AMI, whereas cardiac and lung sEVsl⍰MPC1 levels were reduced (Figure 7D).

**Figure 7.**
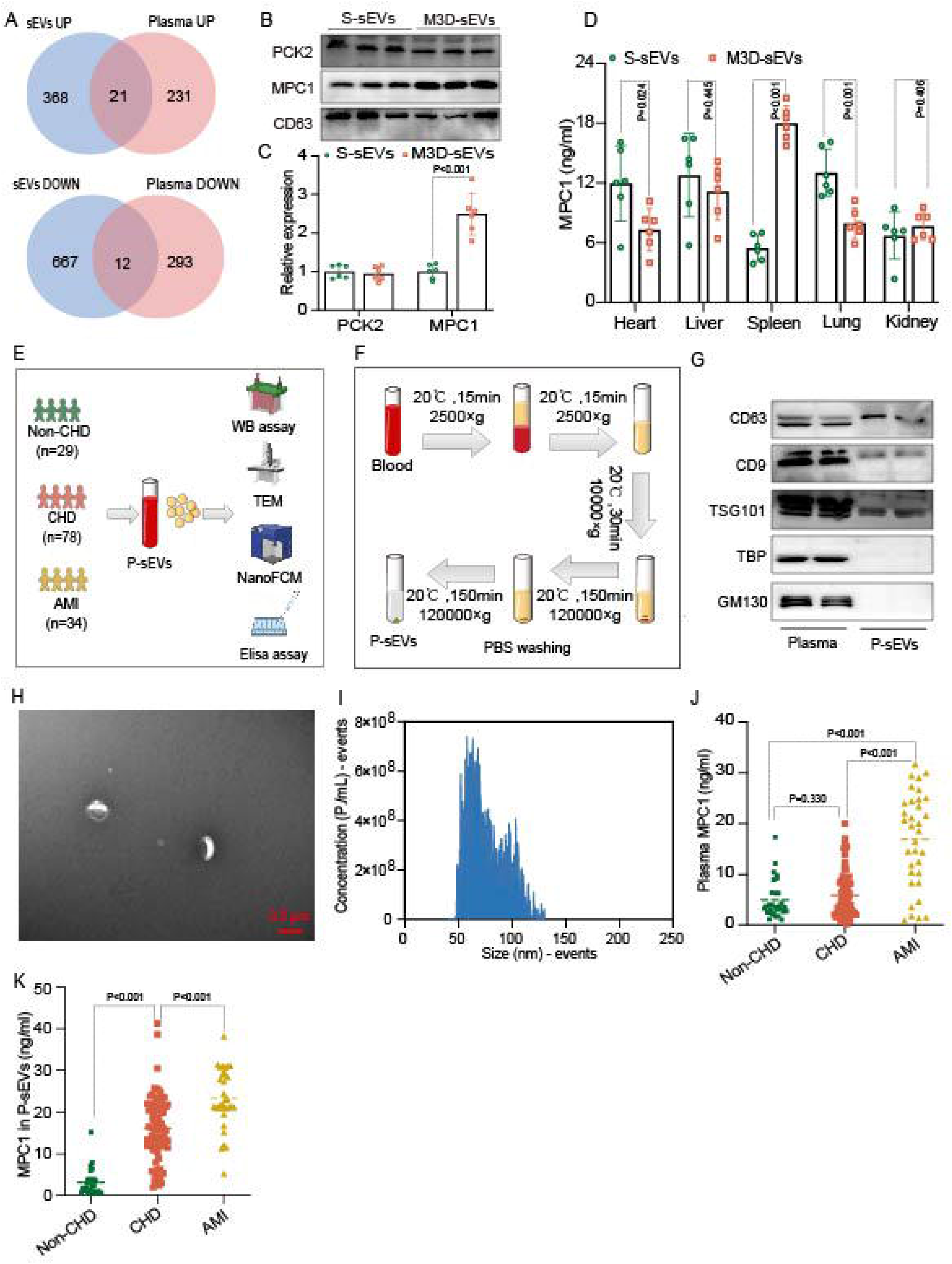
MPC1 protein expression is increased in M3D-sEVs from AMI mouse and in circulating plasma sEVs from AMI patients. **A**, Veen diagrams of differentially expressed proteins in spleen-derived sEVs and plasma between sham and AMI mouse. **B-C**, Representative western blotting images and quantitation of MPC1 and PCK2 expression inside sEVs from sham and AMI mouse. **D**, Quantitation results of ELISA analysis determining MPC1 expression in organ tissues-derived sEVs after AMI. **E**, Schematic diagram of plasma sEVs (P-sEVs) isolation from patients by commercial kits or ultracentrifugation (UC). **F**, The flowchart of plasma-derived sEVs extraction by UC. **G**, Immunoblot images of sEVs markers. **H,** Representative TEM images of P-sEVs by UC. **I**, Size distributions of P-sEVs measured by nFCM. **J**, ELISA assay results of the plasma MPC1 expression from patients. **K**, ELISA assay results of MPC1 expression in P-sEVs by UC. Statistical analyses were performed by the unpaired two-tailed Welch t-test (**C**; MPC1), the unpaired t test (**C**; PCK2 and **D**), the Kruskal-Wallis with the Dunn multiple comparisons test (**I**) and the one-way ANOVA with the Dunnett multiple comparisons test (**K**). P-sEVs indicates plasma sEVs.

Clinically, plasma sEVs (P-sEVs) from healthy controls (Non-CHD), patients with coronary heart disease (CHD) and patients with AMI were isolated by ultracentrifugation (Figure 7E-7F). The baseline parameters of recruited individuals are summarized in Table S5. Preparations met sEVs identity criteria (TEM, nFCM size 30-150 nm; CD63/CD9/TSG101 positive; GM130 and TBP negative) (Figure 7G-7I). ELISA revealed a stepwise increase in sEVs-encapsulated MPC1 across groups, showing highest in AMI, intermediate in CHD, lowest in controls, whereas plasma MPC1 showed an increase only in AMI but not CHD compared with controls (Figure 7J-7K). Moreover, a commercial isolation kit for extracting plasma sEVs yielded consistent trends (Figure S15A-S15D). These findings suggest that sEVs-associated MPC1 is increased in response to myocardial ischemia and may have biomarker potential, which requires prospective validation.

### MPC1 mediates the metabolic and cardioprotective actions of M3D-sEVs

M3D-sEVs administration increased myocardial MPC1 content after AMI (Figure S16A). Next, to clarify the role of MPC1 in M3D-sEVs, we neutralized MPC1 in M3D-sEVs using a specific antibody (sEVs+Ab) with control IgG as comparator (sEVs+IgG). Both were administered via intramyocardial injection in AMI mice. This neutralization significantly reduced the beneficial effects of M3D-sEVs, reducing survival and cardiac function (Figure 8A-8D), increasing infarct size (Figure 8E-8F), elevating cardiac fibrosis and fibrotic protein expression (Figure 8G-8J), and enhancing myocyte hypertrophy (Figure 8K-8L).

**Figure 8.**
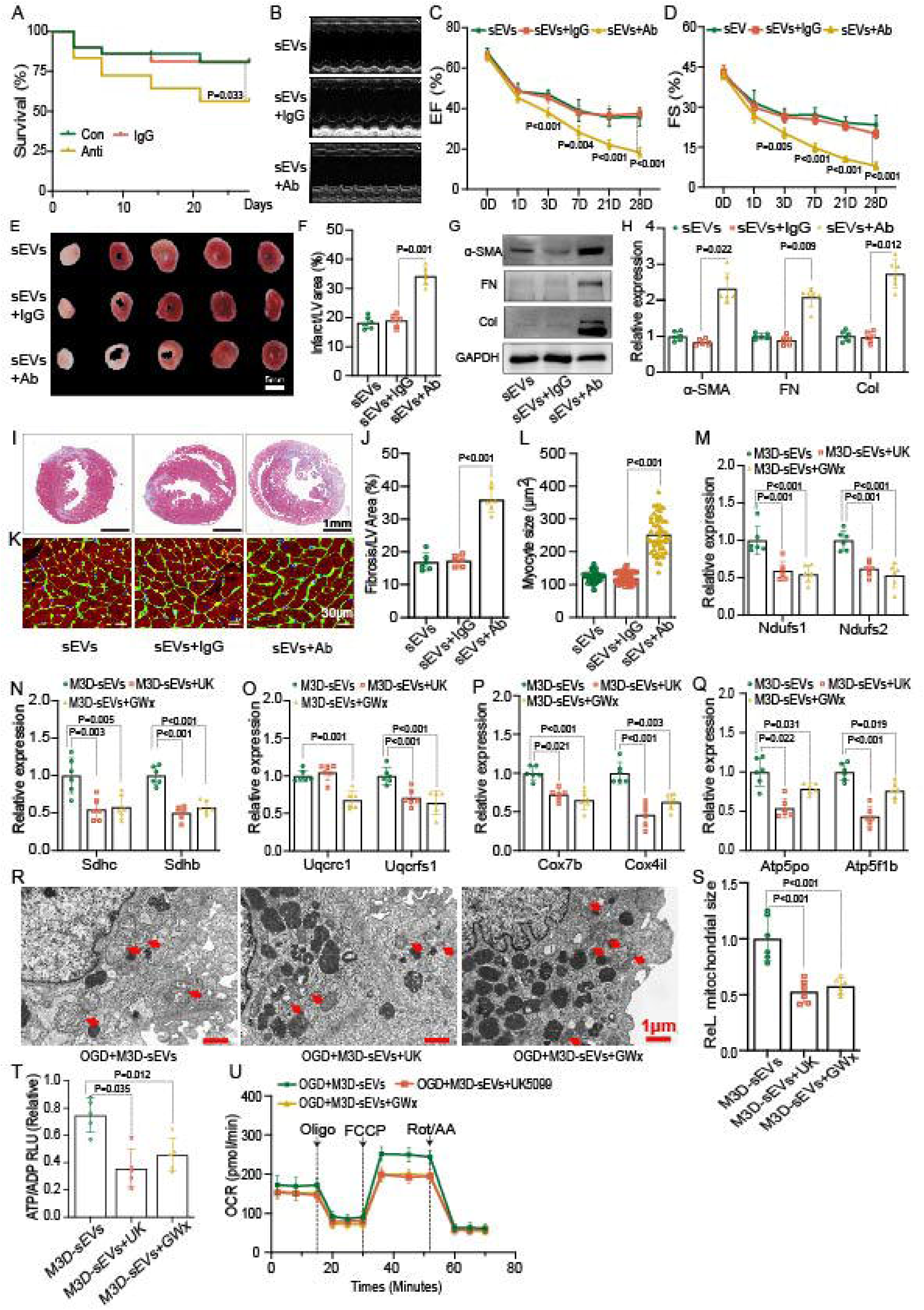
Blockage of MPC1 blunts the protective effects of M3D-sEVs against ischemic cardiac damage and disruption of myocardial mitochondrial energy metabolism. **A**, Twenty-eight-day Kaplan-Meier survival curves for AMI mouse treated with M3D-sEVs (sEVs), MPC1 antibody-neutralized M3D-sEVs (sEVs+Ab), or IgG-incubated M3D-sEVs (sEVs+IgG), n=30/group. **B**, Representative echocardiograms of AMI mice treated with indicated conditions 7 days after AMI (n=6/group). **C-D**, Quantitation of EF% and FS%. Detailed echo parameters at day 7 post AMI were shown in Table S10. **E-F**, Representative TTC staining images of cross-sectional cardiac slices from AMI mice treated as above indicated 1 day post AMI, and quantitation of infarct area percentage (n=6/group). **G-H**, Representative western blotting images and quantitation of cardiac collagen I, fibronectin, and α-SMA expression in AMI mice treated as above indicated 14 days after AMI (n=6/group). **I-L**, Representative images of Masson’s trichrome (top) and WGA staining (bottom) and quantitation of the hearts from AMI mice treated as above indicated 14- or 28-days post AMI, respectively (n=6/group). **M-Q**, mRNA expression of ETC-related genes in the OGD-insulted H9C2 cells treated with M3D-sEVs or M3D-sEVs plus MPC1 inhibitors (n=6/group). **R**, Representative TEM images showing mitochondrial ultrastructure of heart tissues from AMI mice treated as above indicated. **S**, Quantitation of mitochondrial size in TEM images as indicated groups, being presented as relative value to the M3D-sEVs group (n=6/group). **T**, Quantitation of ATP/ADP ratio of OGD-H9C2 cells treated as indicated (n=6/group). **U**, Results of OCR assay in H9C2 cells treated as indicated. Statistical analyses were performed using mixed-effects ANOVA followed by the Sidak multiple comparisons test (**C** and **D**), the one-way ANOVA with the Dunnett multiple comparisons test (**F**, **J** and **S**), the one-way ANOVA with the Tukey multiple comparisons test (**H**; Collagen I, **M**; Ndufs2, **N**, **O**, **P** and **Q**), the Brown-Forsythe and Welch-ANOVA test with the Dunnett T3 multiple comparisons test (**T**), and the Kruskal-Wallis with the Dunn multiple comparisons test (**H**; α-SMA, Fibronectin, **L**, **M**; Ndufs1). The survival curve rate was analyzed by the Kaplan-Meier method and compared by the log-rank test in **A**. UK, UK5099 (a MPC1 inhibitor). GWx, GW604714x. UK indicates UK5099; GWx, GW604714x. EF, ejection fraction; FS, fractional shortening; LV, left ventricular; FN, Fibronectin; Col, collagen I; α-TuB, α-Tubulin.

To further validate the role of MPC1, we employed MPC1 inhibitors, including UK5099 and GW604714x (GWx), to explore whether MPC1 contributes to the therapeutic outcomes of M3D-sEVs. Pharmacological MPC1 inhibition reduced mitochondrial pyruvate levels in H9C2 cells (Figure S16B) and abrogated M3D-sEVs-induced restoration of ETC transcripts in AMI hearts (Figure 8M-8Q), worsened mitochondrial ultrastructure by TEM (Figure 8R-8S), and blunted improvements in ATP/ADP ratio and oxygen consumption under OGD (Figure 8T-8U). Collectively, these findings identify MPC1 as a critical effector of M3D-sEVs-mediated preservation of mitochondrial energetics and cardioprotection after AMI.

## Discussion

In this study, we identify a spleen-heart communication pathway during AMI whereby spleen-derived sEVs exert endogenous cardioprotection. Myocardial ischemia upregulated splenic sEVs biogenesis pathways, increased splenic sEVs release, and promoted recruitment of these vesicles to injured myocardium. Functionally, day-3 post-AMI spleen sEVs (M3D-sEVs) limited infarct size, apoptosis, inflammation, and adverse remodeling, and improved cardiac function. Proteomics and functional evaluations implicate MPC1 as a key effector, showing that M3D-sEVs were enriched for MPC1, enhanced myocardial MPC1 content, restored oxidative phosphorylation and ATP generation. These benefits were abrogated by MPC1 neutralization or inhibition. Clinically, plasma sEVs-encapsulated MPC1 was highest in AMI, intermediate in CHD, and lowest in non-CHD controls, supporting potential translational relevance. To our knowledge, this is the first study linking splenic sEVs cargo to myocardial metabolic rescue via MPC1 in AMI.

The concept of a cardio-splenic axis as a critical regulator of ischemic heart disease is well supported by previous studies^3,27^. Increased metabolic activity of spleen was observed in patients with acute coronary syndrome, which was measured by (18)F-fluorodeoxyglucose-positron emission tomography imaging, and it was associated with systemic proinflammatory status and higher risks in cardiovascular disease events^28^. In rodent models, myocardial ischemia leads to splenic hypertrophy^29^, and myocardial ischemia/reperfusion (I/R) results in a rapid upregulation of splenic leukocyte mobilization-related gene expression (4h post reperfusion)^30^. Consistent with these reports, we observed acute splenic activation post AMI in mice, characterized by increased spleen size, splenic weight, and expansion of the white pulp area and marginal zone, as well as elevated expression of inflammatory cytokines (IL-6, IFN-γ, TNF-α, IL-10) in the spleen, which is consistent with Huizhen Lv and colleagues’ study^31^. These findings support the notion that the spleen responds dynamically to myocardial injury, possibly through recognition of damage-associated molecular patterns derived from the ischemic heart^27^.

The role of spleen in ischemic heart disease remains controversial. Recently, Qiang Long and colleagues reported that splenectomy inhibited the infiltration of phagocytic monocytes into the injured myocardium at the early phases and eventually improved cardiac functions in I/R mice with ischemia for 60 min^30^. Similarly, splenectomy significantly reduced cardiac infarct size in mice with ischemia for 40/50 min and subsequent reperfusion for 60 min by lowering neutrophil infiltration in ischemic myocardium. Moreover, the investigation in mice also showed that AMI enhanced platelet releasing from spleen to facilitate myocardial inflammation, while splenectomy blocked platelets and leucocytes accumulation in heart, reduced infarct size, and improved cardiac functions^32^. In contrast, the epidemiological data from World War II veterans suggest that splenectomy increases mortality rates following myocardial infarction^33^. Consistent with this reports, a recent study showed that splenectomy suppressed inflammation and exerted beneficial outcomes in long term cardiac repair in AMI mice^23^. Our data show that splenectomy immediately prior to MI surgery worsened survival, increased infarction and fibrosis, and aggravated hypertrophy, indicating a net early protective role. This aligns with reports showing that remote ischemic conditioning or neuromodulation in rats and pigs enhances the release of splenic cardioprotective factors, where splenectomy abrogates the deploying of these splenic factors and blunts the cardiac protective effects of remote ischemic condition^3,34^. However, the precise nature of these protective factors has remained elusive.

Most previous studies have focused on splenic immune cells as mediators of post-ischemic injury^30^. A set of studies demonstrated that splenic neutrophils, monocytes, and lymphocytes could be rapidly released into circulation post ischemic insults in hearts^30,32,35^. The splenic immune cells exhibit dual roles in ischemic cardiac injury. For example, Ly6C^hi^ monocytes deployed from subcapsular red pulp of the spleen infiltrate to the infarct region of ischemic hearts, while inhibiting Ly6C high monocyte infiltration promotes cardiac repair and improves cardiac function after AMI^35^. In contrast, adoptive transfer of regulatory T cells (Tregs) enhanced cardiac repair post AMI, increasing ejection fraction and reducing fibrosis^36^. Recently, Mohamed Ameen Ismahil and colleagues found splenic CD169^+^Tim4^+^ marginal metalophilic macrophages from marginal zone of spleen increased in blood 24h post AMI, and these cells alleviated inflammation and improved cardiac function after AMI in mice^23^. The dual and context-dependent roles of these cell populations have complicated the development of targeted therapies for ischemic heart disease. This complexity prompted us to investigate novel mechanisms of spleen-heart communication, focusing on sEVs as potential mediators.

sEVs are established mediators of inter-organ signaling in cardiovascular injury^37,38^, modulating the pathophysiology of ischemic heart disease by delivering diverse biomolecules, including proteins, microRNAs, and lipids^39,40^. Remote ischemic preconditioning (RIPC) protects against myocardial injury, likely by increasing circulating sEVs and underscoring the essential role of exosomes in cardiac protection^41,42^. Our transcriptomic analysis revealed that genes associated with extracellular vesicle biology are upregulated in the spleen following AMI, prompting us to characterize the temporal dynamics and functional properties of spleen-derived sEVs. We found that splenic sEVs release peaks at day 3 post-AMI, and that spleenl⍰derived sEVs enter the shared circulation and are recruited to infarcted myocardium in parabiosis experiments. Methodologically, we implemented controls to address known pitfalls in sEVs tracking, where DiR/PKH26 labelling was followed by iodixanol density gradient ultracentrifugation to remove free dye/aggregates^43^, and PBS+dye controls were processed identically and showed negligible signal. Fluorescence biodistribution demonstrated that M3D-sEVs exhibited preferential accumulation in ischemic hearts compared with non-cardiac organs and could be uptake up by cardiomyocytes. The administration of these vesicles to recipient AMI mice markedly attenuates cardiac injury, improves systolic function, and increases survival. These findings are consistent with emerging evidence that organ-specific sEVs can modulate inflammation, promote tissue repair, and limit ischemic injury^44^. Our results suggest that splenic sEVs deliver reparative cargo, including metabolic regulators, to the injured myocardium, thereby coupling immune and metabolic adaptation.

We also implemented rigorous approaches to ensure sEVs purity. Mouse splenic sEVs were isolated by differential ultracentrifugation and consistently met sEVs identity criteria (TEM morphology, 30-150 nm size, CD63/CD9/TSG101 positive, GM130/TBP negative). Patient plasma sEVs for ELISA were isolated by a harmonized ultracentrifugationl⍰based workflow with equivalent purity controls.

Splenic regulation of inflammation has a significant impact in postl⍰AMI outcomes^45^. Splenic monocytes can be activated by mitochondria-cell-free DNA from I/R-induced myocardium, causing inflammatory damage^45^. Splenectomy significantly inhibited macrophage-mediated inflammation in mouse ischemic hearts^46^. In our study, however, M3D-sEVs decreased myocardial IL-1β, IL-6, and TNF-α while improving heart function. These observations support a dual role for the spleen that produces inflammatory cells but also releases a subset of protective sEVs that preserves myocardial energetics. In the early post-AMI window, the net effect of splenic signaling may depend on the balance between inflammatory cell releasing and protective sEVs cargo.

We further examined systemic inhibition of sEVs biogenesis using the neutral sphingomyelinase inhibitor GW4869. Consistent with prior reports in I/R and inflammationl⍰dominated models^47–49^, a single dose (GW4869-S) in our study improved LV function and reduced infarct size. In contrast, a repeated-dose regimen (GW4869-M) worsened survival, systolic function, and remodeling in AMI mice. Our time-course data provide a potential explanation that GW4869-S suppressed splenic sEVs release for approximately 24 h and then allowed a rebound above vehicle levels by day 3, whereas GW4869-M produced a sustained reduction in splenic sEVs throughout the early post-AMI period, including the peak of M3D-sEVs release. Thus, GW4869-S may result in compensatory upregulation of reparative splenic sEVs, whereas GW4869-M could dramatically deplete the cardioprotective M3D-sEVs. This interpretation is supported by the observation that the adverse effects of GW4869-M were rescued by add-back of M3D-sEVs. GW4869 remains a broad, systemic agent, and work on inhibiting de novo ceramide synthesis highlights an additional, cell-intrinsic ceramide pathway that is not specific to sEVs release^50^. Overall, our findings argue that therapeutic modulation of sEVs should be temporally and source-selective, targeting specific sEVs subsets and windows of vulnerability rather than applying uniform, prolonged suppression.

Mechanistically, M3D-sEVs preserved oxidative phosphorylation, restored ETC subunits, improved oxygen consumption, and reduced mitochondrial ROS in cardiomyocytes under ischemic stress. Proteomic profiling of M3D-sEVs identified significant enrichment of proteins involved in oxidative phosphorylation, with MPC1 showing as a key candidate. Neutrali⍰ng or pharmacologically inhibiting MPC1 abolished the M3D-sEVs benefits on mitochondrial structure and function, implicating MPC1 as a critical effector. Importantly, increased MPC1 expression was unique to spleen-derived sEVs post AMI in mice and not observed in sEVs from other main organs. Functionally, M3D-sEVs enhanced MPC1 expression in recipient cardiomyocytes, improved mitochondrial ATP/ADP ratios, while reduced mitochondrial ROS generation, which are the hallmarks of optimized mitochondrial bioenergetics and reduced oxidative stress.

MPC1 is responsible for mitochondrial pyruvate import, a critical step in maintaining intracellular pyruvate and lactate balance and limiting lactate accumulation^51,52^. Previous studies have shown that impaired pyruvate metabolism and mitochondrial dysfunction exacerbate ischemic injury and that MPC1 downregulation is associated with heart failure and increased susceptibility to cell death^16,53,54^. Besides, Yue Liu and colleges demonstrated that MPC1 downregulation enhances susceptibility to ischemic injury by increasing mitochondria disruption, oxidative stress, calcium overload, and autophagy in cortical neurons^55^. Conversely, augmenting MPC1 expression or activity have been shown to restore mitochondrial metabolism and ameliorate cardiomyocyte injury in models of ischemia-reperfusion. Our findings demonstrate that M3D-sEVs-mediated delivery of MPC1 constitutes a novel, endogenous rescue pathway during AMI. The observed mitochondrial benefits, such as preserved ATP generation and reduced ROS accumulation, highlight the importance of maintaining efficient mitochondrial pyruvate flux for cardioprotection during acute ischemic stress. Thus, splenic sEVs-mediated delivery of MPC1 represents a promising therapeutic axis for preserving mitochondrial function during acute ischemia, warranting further mechanistic study and consideration for translational application in AMI.

From a translational perspective, sEVs-associated biomarkers are attractive because they are stable in circulation, reflect dynamic pathophysiological changes, and may retain organ-specific information^56^. Recent work indicates that a substantial proportion of plasma sEVs originate from the spleen in AMI patients^12^, supporting the feasibility of targeting splenic sEVs clinically. We observed that a stepwise increase in plasma sEVs-associated MPC1 across non-CHD controls, CHD and AMI patients, suggesting a close association between sEVs-associated MPC1 and disease burden. sEVs-associated markers may offer advantages over traditional biomarkers such as troponins, as MPC1 enrichment may reflect myocardial metabolic stress and mitochondrial adaptation rather than necrosis alone. sEVs-associated MPC1 might therefore complement established markers in early AMI diagnosis and risk stratification, particularly in challenging scenarios, such as early presenters, equivocal troponin changes or coexisting renal dysfunction. sEVs-based diagnostic platforms have already proven feasible in several cardiac and non-cardiac conditions^57^. Our data raise the possibility that plasma sEVsl⍰MPC1, alone or combined with troponins and natriuretic peptides, might improve diagnostic accuracy and prognostication in acute coronary syndromes. Larger, multicenter prospective studies are required to validate these findings, define optimal cutl⍰offs and establish standardized isolation and quantification protocols.

This study has several limitations. First, the murine and single-center clinical data, although internally consistent, may not fully capture the complexity of sEVs biology and MPC1 regulation in humans. The clinical sample size was modest, and residual confounding cannot be excluded. Second, although we provide convincing evidence for the role of MPC1-enriched spleen-derived sEVs in enhancing mitochondrial function and cardioprotection, the precise molecular mechanisms underlying sEVs-associated MPC1 delivery, cardiac uptake, and downstream signaling pathways were not fully described. Third, GW4869 acts systemically and is not spleen-specific; cell- or organ-targeted genetic approaches such as specific knockout of Rab27a in splenic myeloid cells would strengthen causal inference. Finally, the temporal kinetics of splenic sEVs-associated MPC1 elevation following AMI was not fully characterized, and more detailed time-course studies are warranted.

In summary, our data identify a spleen-heart axis in which AMI induces an adaptive spleen sEVs program that delivers MPC1 to the injured myocardium to preserve mitochondrial bioenergetics and limit structural and functional damage. These findings demonstrate MPC1-enriched splenic sEVs as an endogenous cardioprotective mechanism and suggest that selectively augmenting this cargo-specific pathway, either via targeted delivery of MPC1-loaded sEVs or sEVs mimetics, or by enhancing splenic release of reparative sEVs, warrants further investigation as a novel therapeutic strategy to improve outcomes in ischemic heart disease.

## Acknowledgements

None.

## Data availability statement

Data supporting the findings of this study are available from the corresponding author upon reasonable request.

## References

1. Saito Y, Oyama K, Tsujita K, Yasuda S, Kobayashi Y. Treatment strategies of acute myocardial infarction: updates on revascularization, pharmacological therapy, and beyond. J Cardiol 2023;81:168–178. doi: 10.1016/j.jjcc.2022.07.003

2. Rein H. The role of the spleen and liver in coronary or hypoxic myocardial insufficiency. Pflugers Arch Gesamte Physiol Menschen Tiere 1951;253:435–458. doi: 10.1007/BF00370032

3. Heusch G, Kleinbongard P. The spleen in ischaemic heart disease. Nat Rev Cardiol 2025. doi: 10.1038/s41569-024-01114-x

4. Tian Y, Miao B, Charles EJ, Wu D, Kron IL, French BA, et al. Stimulation of the Beta2 Adrenergic Receptor at Reperfusion Limits Myocardial Reperfusion Injury via an Interleukin-10-Dependent Anti-Inflammatory Pathway in the Spleen. Circ J 2018;82:2829–2836. doi: 10.1253/circj.CJ-18-0061

5. Lieder HR, Paket U, Skyschally A, Rink AD, Baars T, Neuhauser M, et al. Vago-splenic signal transduction of cardioprotection in humans. Eur Heart J 2024;45:3164–3177. doi: 10.1093/eurheartj/ehae250

6. Rohm TV, Cunha ERK, Olefsky JM. Metabolic Messengers: small extracellular vesicles. Nat Metab 2025;7:253–262. doi: 10.1038/s42255-024-01214-5

7. Xiao J, Sluijter JPG. Extracellular vesicles in cardiovascular homeostasis and disease: potential role in diagnosis and therapy. Nat Rev Cardiol 2025. doi: 10.1038/s41569-025-01141-2

8. Kumar MA, Baba SK, Sadida HQ, Marzooqi SA, Jerobin J, Altemani FH, et al. Extracellular vesicles as tools and targets in therapy for diseases. Signal Transduct Target Ther 2024;9:27. doi: 10.1038/s41392-024-01735-1

9. Li H, Ding J, Liu W, Wang X, Feng Y, Guan H, et al. Plasma exosomes from patients with acute myocardial infarction alleviate myocardial injury by inhibiting ferroptosis through miR-26b-5p/SLC7A11 axis. Life Sci 2023;322:121649. doi: 10.1016/j.lfs.2023.121649

10. Duan S, Wang C, Xu X, Zhang X, Su G, Li Y, et al. Peripheral Serum Exosomes Isolated from Patients with Acute Myocardial Infarction Promote Endothelial Cell Angiogenesis via the miR-126-3p/TSC1/mTORC1/HIF-1alpha Pathway. Int J Nanomedicine 2022;17:1577–1592. doi: 10.2147/IJN.S338937

11. Wang B, Cao C, Han D, Bai J, Guo J, Guo Q, et al. Dysregulation of miR-342-3p in plasma exosomes derived from convalescent AMI patients and its consequences on cardiac repair. Biomed Pharmacother 2021;142:112056. doi: 10.1016/j.biopha.2021.112056

12. Jin X, Xu W, Wu Q, Huang C, Song Y, Lian J. Detecting early-warning biomarkers associated with heart-exosome genetic-signature for acute myocardial infarction: A source-tracking study of exosome. J Cell Mol Med 2024;28:e18334. doi: 10.1111/jcmm.18334

13. Thygesen K, Alpert JS, Jaffe AS, Chaitman BR, Bax JJ, Morrow DA, et al. [Fourth universal definition of myocardial infarction (2018)]. Kardiol Pol 2018;76:1383–1415. doi: 10.5603/KP.2018.0203

14. Yang R, Ruan B, Wang R, Zhang X, Xing P, Li C, et al. Cardiomyocyte betaII spectrin plays a critical role in maintaining cardiac function by regulating mitochondrial respiratory function. Cardiovasc Res 2024;120:1312–1326. doi: 10.1093/cvr/cvae116

15. Ong SB, Samangouei P, Kalkhoran SB, Hausenloy DJ. The mitochondrial permeability transition pore and its role in myocardial ischemia reperfusion injury. J Mol Cell Cardiol 2015;78:23–34. doi: 10.1016/j.yjmcc.2014.11.005

16. Visker JR, Cluntun AA, Velasco-Silva JN, Eberhardt DR, Cedeno-Rosario L, Shankar TS, et al. Enhancing mitochondrial pyruvate metabolism ameliorates ischemic reperfusion injury in the heart. JCI Insight 2024;9. doi: 10.1172/jci.insight.180906

17. Gray LR, Tompkins SC, Taylor EB. Regulation of pyruvate metabolism and human disease. Cell Mol Life Sci 2014;71:2577–2604. doi: 10.1007/s00018-013-1539-2

18. Fernandez-Caggiano M, Prysyazhna O, Barallobre-Barreiro J, CalvinoSantos R, Aldama Lopez G, Generosa Crespo-Leiro M, et al. Analysis of Mitochondrial Proteins in the Surviving Myocardium after Ischemia Identifies Mitochondrial Pyruvate Carrier Expression as Possible Mediator of Tissue Viability. Mol Cell Proteomics 2016;15:246–255. doi: 10.1074/mcp.M115.051862

19. Zhao S, Zhang Y, Bao S, Jiang L, Li Q, Kong Y, et al. A novel HMGA2/MPC-1/mTOR signaling pathway promotes cell growth via facilitating Cr (VI)-induced glycolysis. Chem Biol Interact 2024;399:111141. doi: 10.1016/j.cbi.2024.111141

20. Lewis SM, Williams A, Eisenbarth SC. Structure and function of the immune system in the spleen. Sci Immunol 2019;4. doi: 10.1126/sciimmunol.aau6085

21. Mebius RE, Kraal G. Structure and function of the spleen. Nat Rev Immunol 2005;5:606–616. doi: 10.1038/nri1669

22. Zhang X, Fang Y, Qin X, Zhang Y, Kang B, Zhong L, et al. The Role of MCPIP1 in Macrophage Polarization and Cardiac Function Post-Myocardial Infarction. Adv Sci (Weinh) 2025:e2500747. doi: 10.1002/advs.202500747

23. Ismahil MA, Zhou G, Rajasekar S, Gao M, Bansal SS, Patel B, et al. Splenic CD169(+)Tim4(+) Marginal Metallophilic Macrophages Are Essential for Wound Healing After Myocardial Infarction. Circulation 2025. doi: 10.1161/CIRCULATIONAHA.124.071772

24. Rolland TJ, Hudson ER, Graser LA, Zahra S, Cucinotta D, Weil BR. Splenic Modulation of the Early Inflammatory Response to Regional and Global Ischemia/Reperfusion Injury in Swine. Am J Physiol Heart Circ Physiol 2025. doi: 10.1152/ajpheart.00714.2024

25. Yu J, Sane S, Kim JE, Yun S, Kim HJ, Jo KB, et al. Biogenesis and delivery of extracellular vesicles: harnessing the power of EVs for diagnostics and therapeutics. Front Mol Biosci 2023;10:1330400. doi: 10.3389/fmolb.2023.1330400

26. Luo Y, Liu K, Ling D, Gu T, Zhang L, Chen W. Isolation and Characterization of Exosomes Derived from Mouse Spleen Tissues. J Vis Exp 2024. doi: 10.3791/67234

27. Tian Y, Pan D, Chordia MD, French BA, Kron IL, Yang Z. The spleen contributes importantly to myocardial infarct exacerbation during post-ischemic reperfusion in mice via signaling between cardiac HMGB1 and splenic RAGE. Basic Res Cardiol 2016;111:62. doi: 10.1007/s00395-016-0583-0

28. Emami H, Singh P, MacNabb M, Vucic E, Lavender Z, Rudd JH, et al. Splenic metabolic activity predicts risk of future cardiovascular events: demonstration of a cardiosplenic axis in humans. JACC Cardiovasc Imaging 2015;8:121–130. doi: 10.1016/j.jcmg.2014.10.009

29. Ruggiero SA, Huber JS, Murrant CL, Brunt KR, Simpson JA. Splenic blood-flow response following myocardial infarction in rat. Can J Physiol Pharmacol 2018;96:1060–1068. doi: 10.1139/cjpp-2018-0134

30. Long Q, Rabi K, Cai Y, Li L, Huang S, Qian B, et al. Identification of splenic IRF7 as a nanotherapy target for tele-conditioning myocardial reperfusion injury. Nat Commun 2025;16:1909. doi: 10.1038/s41467-025-57048-6

31. Lv H, Wang C, Liu Z, Quan M, Li K, Gou F, et al. Suppression of the Prostaglandin I2-Type 1 Interferon Axis Induces Extramedullary Hematopoiesis to Promote Cardiac Repair After Myocardial Infarction. Circulation 2025. doi: 10.1161/CIRCULATIONAHA.124.069420

32. Gao XM, Moore XL, Liu Y, Wang XY, Han LP, Su Y, et al. Splenic release of platelets contributes to increased circulating platelet size and inflammation after myocardial infarction. Clin Sci (Lond) 2016;130:1089–1104. doi: 10.1042/CS20160234

33. Robinette CD, Fraumeni JF, Jr. Splenectomy and subsequent mortality in veterans of the 1939-45 war. Lancet 1977;2:127–129. doi: 10.1016/s0140-6736(77)90132-5

34. Lieder HR, Kleinbongard P, Skyschally A, Hagelschuer H, Chilian WM, Heusch G. Vago-Splenic Axis in Signal Transduction of Remote Ischemic Preconditioning in Pigs and Rats. Circ Res 2018;123:1152–1163. doi: 10.1161/CIRCRESAHA.118.313859

35. Zhang J, Hao W, Zhang J, Li T, Ma Y, Wang Y, et al. CXCL16 Promotes Ly6Chigh Monocyte Infiltration and Impairs Heart Function after Acute Myocardial Infarction. J Immunol 2023;210:820–831. doi: 10.4049/jimmunol.2200249

36. Alshoubaki YK, Nayer B, Lu YZ, Salimova E, Lau SN, Tan JL, et al. Tregs delivered post-myocardial infarction adopt an injury-specific phenotype promoting cardiac repair via macrophages in mice. Nat Commun 2024;15:6480. doi: 10.1038/s41467-024-50806-y

37. Huang-Doran I, Zhang CY, Vidal-Puig A. Extracellular Vesicles: Novel Mediators of Cell Communication In Metabolic Disease. Trends Endocrinol Metab 2017;28:3–18. doi: 10.1016/j.tem.2016.10.003

38. Hu S, Hu Y, Yan W. Extracellular vesicle-mediated interorgan communication in metabolic diseases. Trends Endocrinol Metab 2023;34:571–582. doi: 10.1016/j.tem.2023.06.002

39. Tian C, Gao L, Rudebush TL, Yu L, Zucker IH. Extracellular Vesicles Regulate Sympatho-Excitation by Nrf2 in Heart Failure. Circ Res 2022;131:687–700. doi: 10.1161/CIRCRESAHA.122.320916

40. Fu S, Zhang Y, Li Y, Luo L, Zhao Y, Yao Y. Extracellular vesicles in cardiovascular diseases. Cell Death Discov 2020;6:68. doi: 10.1038/s41420-020-00305-y

41. Ding S, Fan Z, Lin C, Dai Q, Zhou J, Huang H, et al. Therapeutic Effects of Ischemic-Preconditioned Exosomes in Cardiovascular Diseases. Adv Exp Med Biol 2017;998:271–281. doi: 10.1007/978-981-10-4397-0_18

42. Giricz Z, Varga ZV, Baranyai T, Sipos P, Paloczi K, Kittel A, et al. Cardioprotection by remote ischemic preconditioning of the rat heart is mediated by extracellular vesicles. J Mol Cell Cardiol 2014;68:75–78. doi: 10.1016/j.yjmcc.2014.01.004

43. Rautaniemi K, Zini J, Lofman E, Saari H, Haapalehto I, Laukka J, et al. Addressing challenges in the removal of unbound dye from passively labelled extracellular vesicles. Nanoscale Adv 2021;4:226–240. doi: 10.1039/d1na00755f

44. Zheng X, Hermann DM, Bahr M, Doeppner TR. The role of small extracellular vesicles in cerebral and myocardial ischemia-Molecular signals, treatment targets, and future clinical translation. Stem Cells 2021;39:403–413. doi: 10.1002/stem.3329

45. Xie D, Guo H, Li M, Jia L, Zhang H, Liang D, et al. Splenic monocytes mediate inflammatory response and exacerbate myocardial ischemia/reperfusion injury in a mitochondrial cell-free DNA-TLR9-NLRP3-dependent fashion. Basic Res Cardiol 2023;118:44. doi: 10.1007/s00395-023-01014-0

46. Tomczyk M, Kraszewska I, Szade K, Bukowska-Strakova K, Meloni M, Jozkowicz A, et al. Splenic Ly6C(hi) monocytes contribute to adverse late post-ischemic left ventricular remodeling in heme oxygenase-1 deficient mice. Basic Res Cardiol 2017;112:39. doi: 10.1007/s00395-017-0629-y

47. Ge X, Meng Q, Wei L, Liu J, Li M, Liang X, et al. Myocardial ischemia-reperfusion induced cardiac extracellular vesicles harbour proinflammatory features and aggravate heart injury. J Extracell Vesicles 2021;10:e12072. doi: 10.1002/jev2.12072

48. Biemmi V, Milano G, Ciullo A, Cervio E, Burrello J, Dei Cas M, et al. Inflammatory extracellular vesicles prompt heart dysfunction via TRL4-dependent NF-kappaB activation. Theranostics 2020;10:2773–2790. doi: 10.7150/thno.39072

49. Essandoh K, Yang L, Wang X, Huang W, Qin D, Hao J, et al. Blockade of exosome generation with GW4869 dampens the sepsis-induced inflammation and cardiac dysfunction. Biochim Biophys Acta 2015;1852:2362–2371. doi: 10.1016/j.bbadis.2015.08.010

50. Lima TI, Laurila PP, Wohlwend M, Morel JD, Goeminne LJE, Li H, et al. Inhibiting de novo ceramide synthesis restores mitochondrial and protein homeostasis in muscle aging. Sci Transl Med 2023;15:eade6509. doi: 10.1126/scitranslmed.ade6509

51. Sichrovsky M, Lacabanne D, Ruprecht JJ, Rana JJ, Stanik K, Dionysopoulou M, et al. Molecular basis of pyruvate transport and inhibition of the human mitochondrial pyruvate carrier. Sci Adv 2025;11:eadw1489. doi: 10.1126/sciadv.adw1489

52. He Z, Zhang J, Xu Y, Fine EJ, Suomivuori CM, Dror RO, et al. Structure of mitochondrial pyruvate carrier and its inhibition mechanism. Nature 2025;641:250–257. doi: 10.1038/s41586-025-08667-y

53. Lesnefsky EJ, Chen Q, Tandler B, Hoppel CL. Mitochondrial Dysfunction and Myocardial Ischemia-Reperfusion: Implications for Novel Therapies. Annu Rev Pharmacol Toxicol 2017;57:535–565. doi: 10.1146/annurev-pharmtox-010715-103335

54. Lopez-Vazquez P, Fernandez-Caggiano M, Barge-Caballero E, Barge-Caballero G, Couto-Mallon D, Grille-Cancela Z, et al. Reduced mitochondrial pyruvate carrier expression in hearts with heart failure and reduced ejection fraction patients: ischemic vs. non-ischemic origin. Front Cardiovasc Med 2024;11:1349417. doi: 10.3389/fcvm.2024.1349417

55. Liu Y, Yuan Y, Yan Y, Wang R, Wang Z, Liu X, et al. Mitochondrial pyruvate carrier 1 alleviates hypoxic-ischemic brain injury in rats. Life Sci 2023;325:121686. doi: 10.1016/j.lfs.2023.121686

56. Garcia-Martin R, Brandao BB, Thomou T, Altindis E, Kahn CR. Tissue differences in the exosomal/small extracellular vesicle proteome and their potential as indicators of altered tissue metabolism. Cell Rep 2022;38:110277. doi: 10.1016/j.celrep.2021.110277

57. Das D, Jothimani G, Banerjee A, Dey A, Duttaroy AK, Pathak S. A brief review on recent advances in diagnostic and therapeutic applications of extracellular vesicles in cardiovascular disease. Int J Biochem Cell Biol 2024;173:106616. doi: 10.1016/j.biocel.2024.106616

