## Supplementary material for "Spleen-derived Small Extracellular Vesicles Protect against Myocardial Infarction via Mediating Spleen-heart Crosstalk": sup

**Supplementary Methods** Pages 3-24

**Supplementary Tables** Pages 25-37

**Supplementary Figures** Pages 38-54

**Supplementary references** Pages 55-56

### Methods and materials

#### Experimental animals

Male C57BL/6 mice (6-8 weeks old, 20-25g) were purchased from Guangzhou Tengke Biomedical Technology Co., Ltd (Guangzhou, China). Animals were housed in standard plastic cages with husk bedding at 25 ± 2°C under a 12-hour light/dark cycle, with free access to food and water, and maintained under specific pathogen-free conditions. All animal procedures were approved by the Guangdong Medical University Animal Care and Use Committee (GDMU-2024-000043) and conformed to the ARRIVE 2.0 guidelines.

Mice were randomly assigned to experimental groups using a computerized randomization sequence. Investigators responsible for outcome assessment and data analysis were blinded to group allocation throughout the study. Animals were monitored daily for general health, activity, body weight, grooming behavior, and signs of distress. Humane endpoints were strictly observed, and any animal displaying persistent distress, severe weight loss, or impaired mobility was promptly euthanized according to institutional guidelines.

#### AMI mouse model

AMI mice model was established as previously described[1]. Briefly, male mice (8 weeks old) were anesthetized with 3% sevoflurane and maintained with 2% sevoflurane (tidal volume 200 μL, 150 breaths/min) on a ventilator. A left thoracotomy was performed between the third and fourth intercostal spaces to expose the heart. The left anterior descending (LAD) coronary artery was ligated using an 8-0 suture approximately 2 mm below the left atrial appendage. The chest wall and skin were closed in layers, and the tracheal tube was removed after surgery. Hearts were harvested at predetermined time points post-AMI. At the end of experiments or upon any sign of significant distress, animals were euthanized by $CO_2$ asphyxiation.

**Splenectomy**

Splenectomy was performed in randomly selected mice undergoing LAD ligation, as previously described[2]. Briefly, following induction of anesthesia and appropriate preparation, a midline abdominal incision of approximately 1 cm was made. The hepatic hilum was gently retracted, and the splenic vessels were carefully ligated before excising the spleen. The peritoneal cavity was inspected for hemostasis, and the muscle and skin layers were closed with 5-0 sterile sutures. Sham-operated mice underwent an identical procedure without removal of the spleen. Animals were monitored closely during recovery, and postoperative analgesia was provided as per institutional guidelines.

**Parabiosis model**

Murine parabiosis experiments were performed following Kamran et al.'s protocol[3]. In summary, two male mice (one CD45.1, one CD45.2) of similar size, co-housed for a minimum of two weeks, were anesthetized using 1.0-1.5% isoflurane. The lateral flanks of each mouse were shaved and disinfected with iodine followed by alcohol. An incision was made in the skin of each mouse from the elbow to the knee, and the animals were joined through continuous suturing. Buprenorphine (0.1 mg/kg, subcutaneously) was administered for postoperative analgesia. Four weeks post-surgery, either coronary ligation or a sham procedure was conducted on the host mouse following verification of chimerism via flow cytometric analysis of donor blood or splenic tissue. In selected instances, either the host or donor mice underwent splenectomy concurrent with AMI as indicated.

**Isolation of spleen sEVs**

Spleen-derived sEVs were isolated following the established protocols by our lab [4]. Briefly, spleen tissues were harvested, minced into small fragments, and processed according to established protocols with collagenase digestion followed by ultracentrifugation to obtain purified sEVs for downstream analyses.

**Transmission electron microscopy (TEM) for sEVs**

Isolated sEVs (2-6 μg/μL) were fixed with 2% electron microscopy (EM)-grade paraformaldehyde at room temperature for 2 h. Ten microliters of sEVs suspension were placed on a copper grid and incubated for 10 min. Grids were rinsed with sterile distilled water, and stained with 2% uranyl acetate for 1 min, and air-dried. Samples were imaged at 80 kV using a transmission electron microscope (JEM-1400plus, JEOL, Tokyo, Japan)[4].

**Transmission electron microscopy (TEM) for mitochondria**

To assess mitochondrial ultrastructure, hearts were perfused with 0.1 M PBS (pH 7.4) immediately after euthanasia. Cardiac tissue was cut into 1 × 1 × 1 $mm^3$ cubes and fixed in cold 2.5% glutaraldehyde, followed by post-fixation in 1% osmium tetroxide. Tissues were then dehydrated, embedded in resin, and ultrathin sections were prepared using an ultramicrotome. Sections were imaged using a JEOL JEM-F200 transmission electron microscope (JEOL, Japan).

**Nano-flow cytometer (n-FCM)**

A nano-flow cytometer was calibrated using standard 250 nm polystyrene nanoparticles. sEVs samples were diluted in PBS to a final concentration of 1-10 ng/μL, as determined by BCA assay. The side scatter intensity (SSI) of sEVs was measured, and particle concentration was calculated by comparing SSI to that of the nanoparticle standard [4].

**Animal treatments with sEVs**

For sEVs administration experiments, AMI was induced as described above. Immediately following LAD ligation, a total of 2 × $10^{12}$ sEVs were resuspended in 15 μL phosphate buffered saline (PBS), with or without anti-MPC1, was injected into three separate injection sites evenly distributed around the ischemia area. All treatments and sample collections were performed by investigators blinded to group allocation.

**In Vivo tracing of sEVs**

DiR (1,1'-dioctadecyltetramethyl indotricarbocyanine iodide, Cat# 100068. MedChemExpress, USA) was dissolved in DMSO at 1 mg/mL. The DiR solution was then mixed with sEVs at a ratio of 2 μg DiR per 100 μg sEVs in PBS and incubated for 30 min, followed by density gradient ultracentrifugation (DGU) to remove unbound DiR and residual DMSO. In brief, DGU centrifuge tubes were prepared with iodixanol solutions of varying concentrations, layered from the bottom up as follows: 1.5 mL of 40%, 2 mL of 20%, 2 mL of 15%, and 1.5 mL of 10%. The sEVs-DiR mixture was then added to the top and centrifuged at 100,000 × g for 16 hours at 4°C. Finally, the corresponding fractions were collected and subjected to ultracentrifugation using a 70 Ti fixed-angle rotor at 100,000 × g for

70 minutes at 4°C to obtain purified DiR-labeled sEVs. The resulting pellet was resuspended in PBS and adjusted to a final concentration of $10^{11}$ particles/mL. Subsequently, PBS+dye (control) or DiR-labeled sEVs were administered to mice via either intraperitoneal (i.p.) injection or intramyocardial injection. In vivo fluorescence imaging was performed during 96 h using an In Vivo Imaging System (IVIS)-like imaging system (Gel View 6000Plus, Biolight, Guangzhou, China).

**sEVs uptake detection in vitro**

sEVs were labeled with PKH26 Red Fluorescent Cell Linker Mini Kit (Sigma) as previously described[5] and the purified procedure was performed similarly as the DiR labeling. H9C2 cells were incubated with PKH26-labeled sEVs ($10^{13}$ particles/mL) for 12 h at 37 °C, washed with PBS buffer, fixed with 4% paraformaldehyde, and stained with phalloidin and DAPI. Uptake was visualized by confocal microscopy (FV3000, OLYMPUS, Tokyo, Japan).

**GW4869 treatments and M3D-sEVs rescue experiments**

To determine the inhibition of splenic exosome release by GW4869 in mice after AMI, we isolated splenic sEVs following a single dose of GW4869 injection (2.5 mg/kg, i.p.) 1 h prior to LAD, and assessed sEVs abundance by quantifying particle numbers using a Nano Coulter G nanoparticle analyzer (RESUN TECH CO., LTD, China).

GW4869 (2.5 mg/kg, i.p.) or vehicle control was further administered once daily for three consecutive days with the initial injection at 1h prior to LAD and following MI induction. M3D-sEVs was co-administrated immediately following LAD ligation for rescue experiments, with Sham-sEVs or DMSO as controls. At the specified time points, echocardiographic assessments were conducted, followed by collecting hearts for further analysis.

**Echocardiography**

Cardiac function was assessed by transthoracic B- and M-mode echocardiography (Vevo 3100 LT, VisualSonics, Toronto, Canada) at specified time points post-AMI. Mice were anesthetized with 2% isoflurane during imaging. Left ventricular ejection fraction (EF%), fractional shortening (FS%) were calculated from long-axis and short-axis M-mode images, averaged over four cardiac cycles. EF and FS were used as primary indicators of cardiac function[6]. Other cardiac parameters were also analyzed, including heart rate (HR), left ventricle internal diameter in systole (LVIDs), left ventricle internal diameter in diastole (LVIDd), left ventricular end-diastolic volume (LVEDV), and left ventricular end-systolic volume (LVESV).

**TUNEL staining**

Apoptosis was detected using a TUNEL Apoptosis Detection Kit (Yeasen, Shanghai, China) according to the manufacturer's instructions. For in vivo experiments, paraffin-embedded heart sections (5 μm) were fixed using 4% paraformaldehyde in PBS for 1 h at room temperature and permeabilized with 0.5% Triton X-100 for 20 min. Sections were then incubated with TUNEL reagent and counterstained with DAPI. Apoptotic nuclei were visualized by confocal microscopy (FV3000, OLYMPUS, Tokyo, Japan). Positive percentages were calculated as the number of positive nuclei divided by the total number of DAPI-stained nuclei.

**Hematoxylin and Eosin (H&E) staining**

Heart tissues were fixed in 4% paraformaldehyde at 4°C for 6 h, dehydrated, and embedded in paraffin. Sections (5 μm) were stained with hematoxylin (3 min) and eosin (15 sec) at room temperature. Images were acquired using a digital pathology scanner (Leica, Wetzlar, Germany).

**Masson's trichrome staining**

Paraffin-embedded heart sections (5 μm) were stained using a Trichrome Stain Kit (ab150686, Abcam, Cambridge, UK) according to the manufacturer's protocol. Collagen fibers stained blue and muscle fibers red were visualized with a digital pathology scanner (Leica, Wetzlar, Germany).

**TTC staining**

Myocardial infarct size was assessed by 2,3,5-triphenyltetrazolium chloride (TTC) staining. Excised hearts were frozen, sliced into 5 sections, and incubated in 2% TTC at 37°C for 30 min. Infarcted (unstained) and viable (red) areas were quantified from five sections per mouse.

**Wheat germ agglutinin (WGA) staining**

Frozen heart sections were cut at mid-ventricle level and fixed with 4% paraformaldehyde for 15 min, washed, and incubated with Alexa Fluor 555-conjugated WGA reagent (1:200) for 20 min at room temperature in the dark. The cells were further counterstained with anti-troponin I (1:200) and DAPI. Secondary antibodies were obtained from Abcam. Sections were imaged and cardiomyocyte cross-sectional size was randomly measured in the confocal microscope (FV3000, OLYMPUS, Tokyo, Japan). A total of 50-60 cardiomyocytes from each group were quantified.

**Construction of AAV vectors for the GFP-Labeled CD63 or CD9 and**

**treatments**

The recombinant adeno-associated virus 2 (AAV2) expressing CD9 or CD63 tag-fused proteins was developed by Hanbio Biotechnology, Shanghai, China. The coding sequence (CDS) of mouse CD9 was amplified via polymerase chain reaction (PCR) using the forward primer 5'-ATGCCGGTCAAAGGAGGTAGCAAGTAGCAAGTGCAT-CAAATAC-3' and the reverse primer 5'-GACTCAACGGGAGATTAGAAGTATTCGTT-ATA-3'. Similarly, the CDS of mouse CD63 was amplified using the forward primer 5'-GGGGAATTCACCACCATGGCGG-TGGAAGGAGGAAT-3' and the reverse primer 5'-ATGGGGCCCCCCTACATTACTT-TCAT-AGCCAC-3'. This DNA fragment was subsequently cloned into the pAAV-ITR-CD45.1promoter-GFP plasmid to create the pAAV-ITR-CD45.1-mCD63(CD9)-GFP vector. An empty plasmid was utilized as the control vector. Following the establishment of parabiosis models for a duration of two weeks, recipient mice were administered an intravenous injection of $1\times 10^{11}$ viral genomes/mouse of AAV2-CD45.1-CD9 (CD63)-GFP or AAV2-CD45.1-Scramble virus. Two weeks after the injection, splenectomy models were induced in these mice.

**Construction of AAV vectors for Rab27a knockdown in immune cells and treatments**

AAV2 vectors carrying the short hairpin Rab27a (shRab27a) or scramble control were designed and constructed by the Hanbio Biotechnology, Shanghai, China. The sequences of shRab27a were as follows: #1: GCTTCTGTTCGACCTGACAAA, #2: CCAGTACACTGATGGCAAGTT, and #3:

GCCAGTTTAAGAGAAGTGTTT. The titer of the AAV2 vectors were 1 × $10^{12}$ vector genome (v.g.)/mL. Following the establishment of parabiosis models for two weeks, recipient mice were injected intravenously with a density of 1× $10^{11}$ viral genomes/mouse of AAV2-CD45.2-shRab27a or AAV2-CD45.2-Scramble virus. Two weeks post-injection, we induced AMI models in these mice, performing the procedure with or without splenectomy as previously outlined.

**Flow cytometry**

Blood cells were first incubated with anti-mouse CD16/32 at 4°C for 10 min to block Fc receptors. The cells were then stained for 60 min at 4°C with fluorochrome-conjugated antibodies in staining buffer. The panel included CD45.2-PE and CD45.1-FITC. Information on antibodies and reagents is in Table S1 or Table S3. Flow cytometry was performed on a BD LSR II instrument. Data were analyzed using FlowJo software (v10.6.1). Fluorescence compensation was set using controls, with unstained cells used for accurate gating. For high-dimensional analysis, t-SNE algorithm was executed via FlowJo. Samples were down-sampled to 10,000 events before t-SNE analysis. The t-SNE heat maps display fluorescence intensity of cell surface markers, with color scale showing expression levels.

**Immunofluorescent staining**

Tissue was embedded in optimal cutting temperature compound (Tissue Tek) and cryopreserved at -80ºC. Tissue sections with a thickness of 5 μm were thawed at room temperature, and fixed with 1% paraformaldehyde for 20 min. The sections were then incubated overnight at 4°C with a combination of anti-cTNT (Abcam,

USA) and either anti-CD63 or anti-CD9 (Proteintech, China). Then, the sections underwent three washes with PBS. The slides were incubated with respective secondary antibodies (Abcam, USA) in the dark for 1 h. Six random fields of view from the border zone of each infarcted heart section were imaged using a fluorescence microscope (FV3000, OLYMPUS, Tokyo, Japan). The percentage of positive cells was calculated as the number of positive nuclei divided by the total number of DAPI-stained nuclei. Information regarding all antibodies and reagents can be found in Table S1 or Table S3.

**Isolation of cardiac sEVs**

Cardiac sEVs were isolated from heart tissue following protocols described previously[7,8]. Briefly, hearts were thoroughly washed with phosphate-buffered saline (PBS), then dissected and digested in 0.1% type II collagenase (Sigma-Aldrich, USA) at 37°C for 30 min. The digested tissue was centrifuged at 300 × g for 10 min to remove the tissues and cells. The supernatant was sequentially centrifuged at 3,000 × g for 20 min and then at 10,000 × g for 30 min at 4°C to further eliminate residual cellular components. The resulting supernatant was sequentially centrifuged at 120,000 g for 2 h at 4°C to pellet extracellular vesicles. The pellet was washed in PBS and re-centrifuged, then resuspended in 200 μL PBS for downstream applications.

**Isolation of liver sEVs**

Liver sEVs were isolated with minor modifications to previously described methods[9]. Briefly, liver tissue was finely minced and digested in 0.1% collagenase type II (Sigma-Aldrich, St. Louis, MO, USA) at 37°C with gentle shaking. The digested tissue was filtered through a 70 μm cell strainer to remove debris. The filtrate was then sequentially centrifuged at 500 × g for 10 min to remove intact cells, 3,000 × g for 20 min to remove apoptotic bodies and microsomes, and 10,000 × g for 30 min to eliminate large extracellular vesicles. The supernatant was subjected to ultracentrifugation at 120,000 × g for 70 min at 4°C, repeated twice to ensure purity. The final sEVs pellet was resuspended in PBS and stored at -80°C until further use.

**Isolation of lung and kidney sEVs**

Lung and kidney-derived sEVs were isolated using minor modifications to established protocols[10-12]. Briefly, lung and kidney tissues were minced in PBS and digested in 0.1% collagenase type I (Sigma-Aldrich, St. Louis, MO, USA) at 37°C with gentle shaking until complete dissociation. The digested suspension was filtered through a 70 μm cell strainer to remove debris. Filtrates were sequentially centrifuged at 500 × g for 10 min at 4°C, 3,000 × g for 20 min at 4°C, and 10,000 × g for 30 min at 4°C to remove cells, membrane debris, and large extracellular vesicles, respectively. Supernatants were then ultracentrifuged at 120,000 × g for 70 min at 4°C to pellet sEVs. The pellets were washed once with ice-cold PBS and re-pelleted under the same conditions. The final sEVs were resuspended in PBS for immediate analysis or stored at -80°C.

**Cell culture**

H9C2 rat cardiomyocytes (SNL-029, SUNNCELL, Wuhan, China) were cultured in DMEM (Cat# 11965092, Thermo Fisher Scientific, Pittsburgh, PA, USA) supplemented with 10% fetal bovine serum (FBS; Cat# 26140079, Thermo Fisher

Scientific, Pittsburgh, PA, USA) and 2% penicillin-streptomycin (C0222, Beyotime, Beijing, China) at 37°C in a humidified atmosphere with 5% $CO_2$. Culture medium was replaced every 3 days. Only cells in the exponential growth phase were used for experiments.

**OGD model and treatments**

In vitro myocardial ischemia was induced using an oxygen and glucose deprivation (OGD) model. Briefly, H9C2 cells were transferred to serum-free, glucose-free DMEM and incubated at 37°C in a hypoxic atmosphere (95% $N_2$/5% $CO_2$) for 12 h. For MPC1 inhibition studies, GW604714X (60 nM) or UK-5099 (50 μM) was added to the culture medium during OGD. For sEVs treatment, sEVs (30 μg total protein) or an equivalent volume of PBS as a control was added to the medium during OGD.

**Protein content measurement**

The protein content of isolated sEVs was quantified using the Pierce BCA Protein Assay Kit (Thermo Fisher Scientific, Waltham, MA, USA) according to the manufacturer's instructions.

**Western blotting**

Protein lysates were separated by 7.5-12% SDS-PAGE and transferred to polyvinylidene fluoride (PVDF) membranes (Millipore, Billerica, MA, USA). Membranes were blocked in 5% non-fat milk in Tris-buffered saline with 0.1% Tween-20 (TBST), then incubated with primary antibodies (Table S1) overnight at 4°C. After washing, membranes were incubated with appropriate HRP-conjugated secondary antibodies (Table S1) and developed using an enhanced chemiluminescence substrate (BL520A, Biosharp, Beijing, China). Protein bands were imaged (Gel View 6000Plus, Biolight, Guangzhou, China) and quantified using ImageJ software (NIH, Bethesda, MD, USA).

**RNA isolation and real-time quantitative PCR (RT-qPCR)**

Total RNA was extracted from tissues or H9C2 cells using TRIzol reagent (Invitrogen) according to the manufacturer's instructions. cDNA synthesis was performed using the PrimeScript RT reagent kit (Takara Bio, Otsu, Japan). RT-qPCR was conducted with SYBR Green Master Mix (Applied Biosystems, Thermo Fisher Scientific) in 20 μL reactions, with GAPDH as internal control. Relative mRNA expression levels were calculated using the $2^{-\Delta\Delta CT}$ method. Primer sequences are provided in Table S2.

**Mitochondrial isolation**

Mitochondria were isolated from tissues or cells using a mitochondrial separation kit (C3601, Beyotime, China) with 1 mM PMSF. Cell suspensions were kept on ice for 15 min and then homogenized in a glass homogenizer with 30 strokes. Homogenates were centrifuged at 600 × g for 10 min at 4°C, and supernatants were further centrifuged at 11,000 × g for 10 min. The pellet was collected to extract mitochondrial protein; the supernatant was centrifuged at 12,000 × g for 10 min to obtain cytoplasmic proteins.

**Enzyme-linked immunosorbent assay (ELISA)**

Cytokine levels (IL-6, IL-10, TNF-α, IFN-γ) in plasma, heart, and spleen were measured using ELISA kits (Mlbio, Shanghai, China), according to the manufacturer's instructions. Human MPC1 in plasma and plasma-derived sEVs was quantified with a Human MPC1 ELISA kit (SMK0490HA, MEIKE, Jiangsu, China).

**Oxygen consumption rate (OCR)**

H9C2 cells (5 × $10^5$ cells/mL) were plated in black 96-well plates overnight, treated with drugs or sEVs for 48 h, and OCR was measured using the OCR Plate Assay Kit (E297, Dojindo, Kumamoto, Japan) per manufacturer's instructions. In brief, cardiomyocyte single-cell suspensions were seeded in assay microplates at a density of 1 × $10^4$ cells/100 μL/well in 10% DMEM. Following hypoxic stimulation and specific sEVs incubation, the plates were incubated at 37°C in a $CO_2$-free incubator for 1 hour. Subsequently, 1.5 μM oligomycin (an ATP synthase inhibitor), 2 μM FCCP (a mitochondrial uncoupler), and 0.5 μM Rot/AA (Complex I and Complex III inhibitors) were sequentially added. Fluorescence readings were recorded every 10 minutes to generate OCR kinetic profiles.

**ATP assessment**

H9C2 cells were seeded into a 6-well plate and allowed to adhere and grow until reaching 50% confluency. After treatment, the culture medium was removed, and 200 μL of lysis buffer was added to each well to lyse the cells. The supernatant was obtained after centrifuging the lysate at 12,000g for 5 min at 4 °C. The ATP content in supernatants was measured using an ATP assay kit, following the manufacturer's instructions. Fluorescence spectrophotometry was employed to measure the ATP level in various groups, assessing mitochondrial activity.

**Mitochondrial reactive oxygen species (MitoSOX assay)**

Mitochondrial ROS levels were assessed using the MitoSOX assay as previously described[13]. H9C2 cells (2 × $10^5$) were seeded on coverslips and cultured for 2-3 days, stained with MitoSOX (100 nM) for 10 min, treated with sEVs for indicated time points, washed with 1×PBS, fixed with 4% polyformaldehyde for 10 minutes, permeabilized with 0.1% Triton X-100 in 1×PBS for 10 minutes, and counterstained with DAPI. Images were captured by confocal microscopy (FV3000, Olympus, Japan).

**CK-MB detection**

Blood was collected from the cardiac aorta, centrifuged at 3,000 rpm for 10 min at 4°C within 2 hours, and serum stored at -80°C. Serum creatine kinase-MB (CK-MB) levels were measured using an ELISA kit (Sangong Biotech, D721065, China) according to the manufacturer's instructions.

**Bulk RNA sequencing**

Bulk RNA sequencing was performed on splenic tissue from AMI and sham mice. RNA quality and quantity were initially assessed prior to library preparation and sequencing at LC-BIO Co., Ltd. (Hangzhou, China). Strand-specific libraries (fragment size 300 ± 50 bp) were constructed from 1 ng total RNA. Libraries were quantified and quality-checked using NanoDrop ND-1000 (NanoDrop, Wilmington, DE, USA), Bioanalyzer 2100 (Agilent, CA, USA), and qPCR (Invitrogen SuperScriptTM II Reverse Transcriptase, cat.1896649, CA, USA). Equimolar pooled libraries were sequenced on an Illumina NovaSeq 6000 platform (LC Biotechnology CO., Ltd.， Hangzhou, China) and MGITech MGISEQ2000RS platform, generating 150 bp paired end reads at a minimum depth of 30 million reads per sample.

**Bulk RNA sequencing data analysis**

Raw FASTQ files were processed using Cutadapt (version: cutadapt-1.9) and aligned to the Mus musculus GRCm38 reference genome using HISAT2 (version: hisat2-2.2.1). Gene quantification produced raw count matrices and quality control metrics. Reads were further analyzed with gffcompare (version: gffcompare-0.9.8), normalized using the trimmed mean of M values (TMM), and differential expression was assessed using StringTie/Ballgown (Version: StringTie4-3.0) and DESeq2, with edgeR used for additional comparisons. Genes with a false discovery rate (FDR) $< 0.05$ and absolute fold change $\geq 2$ were considered differentially expressed.

**Gene set enrichment analysis (GSEA)**

Differential gene expression results were ranked using signed p-value for analysis. Enrichment was performed against KEGG and GO gene sets from MsigDB database using R package fgsea (version 1.12.0). Enrichment significance for each gene set was determined through 10,000 permutations, with p-values adjusted using Benjamini-Hochberg procedure. Bar plots showing NES values were generated with ggplot2 (version 3.3.3), where color indicated enrichment direction (negative/positive) and length showed NES magnitude. Barcode plots were created using plotEnrichment function from fgsea package.

**Plasma proteomics analysis**

Plasma samples were collected from sham and AMI mice (3 days post-AMI). Each 100 μg plasma protein sample was diluted to 300 μL using PBS. The diluted samples were denatured, cleaned, reduced, and then incubated with 15 μL of 400 mM iodoacetamide at room temperature for 30 min. Dithiothreitol (200 mM) was used to quench alkylation. Equal hydrophobic and hydrophilic carboxyl magnetic beads were added to each plasma protein sample at a ratio of 15 μg beads/μg protein. All the samples were acidified with 1% formic acid and 100% MS (LC-MS)-grade acetonitrile. Protein-laden beads were isolated and collected using a magnetic rack and washed twice with 70% ethanol and once with ACN. Samples were digested by incubating with 0.1 μg/μL trypsin/LysC solution overnight at 37 °C and then sonicated. For final LC-MS/MS analysis, 2 μg of total peptides from each sample were separated and analyzed on an Orbitrap Fusion Lumos Mass Spectrometer (Thermo Fisher Scientific). A reversed-phase column was used for separation, with 0.1% FA in water as mobile phase A and 0.1% FA in ACN as mobile phase B. The Orbitrap Mass Analyzer (Thermo Fisher Scientific, USA) was operated in the data-dependent acquisition mode for sample analysis. MS1 was performed with a resolution of 120 k, MS2 was performed at a resolution of 45 k, and MS3 was performed with a resolution of 60 k. Data for the TMT-labelled samples were processed using Proteome Discoverer (v.2.4.0.305, Thermo Fisher Scientific) and a built-in Sequest HT search engine. The MS spectra from each run were searched against the UniProt FASTA databases (uniprot-Musmusculus-10090-2021-8. fasta). The search criteria included trypsin as the protease, allowing 2 missed cleavages. Carbamidomethyl was set as a fixed modification, whereas oxidation and acetyl (protein N-terminal) were set as variable modifications. The FDR was set to 0.01 for both peptide spectral matches (PSMs) and peptide levels, the false discovery rate was set at 0.01. Peptide identification was performed with an initial precursor mass deviation of up to 10 ppm and a fragment mass deviation of 0.02 Da, while other parameters were set as default. All the above technologies are provided by LC-BIO Co., Ltd (Hangzhou, China).

**Four-dimensional (4D) label-free proteomic of sEVs**

4D label-free proteomics protocol was as previously described[14]. The buffer (4% SDS, 25 mM ammonium bicarbonate, 100 mM Tris-HCl, 1 mM DTT, pH 7.6) was used for sample lysis and protein extraction from 20μg aliquot of extracted proteins from each sample. The amount of protein was quantified with a Bradford method Protein Assay kit (Bio-Rad, Hercules, CA, USA). Protein digestion by trypsin was performed according to a previously described filter-aided sample preparation (FASP) procedure [15]. For digestion, the protein solution was reduced with 200 mM dithiothreitol for 1h at 37℃ and alkylated with 11 mM iodoacetamide for 30 min at room temperature in darkness. The sample was diluted 4 times by adding 25 mM ammonium bicarbonate (ABC) buffer. Then adding trypsin (trypsin: protein =1:50) and incubating at 37℃ overnight, and the resulting peptides were collected as a filtrate. Nanoflow liquid chromatography-mass spectrometry (LC-MS/MS) analysis of tryptic peptides was performed on timsTOF Pro2 (Bruker, Germany) coupled to an L-3000 ultra-high-pressure system (RIGOL, China) via a nano-electrospray ion source. The mass spectrometer was operated in “top 40” data-dependent mode. Mass spectrometric measurements were performed using the parallel accumulation serial fragmentation (PASEF) acquisition method. The MS raw data for each sample were combined and searched using Proteome Discoverer 2.4 for identification and quantitation analysis. All the above technologies are provided by

ECHO Biotech Co., Ltd. (Beijing, China).

**Proteomics data analysis**

Proteomics RAW files were analyzed with Proteome Discoverer (Thermo Fisher Scientific, USA). Differentially expressed proteins (DEPs) were identified using t-tests (fold change >1.5 or <0.67, $P < 0.05$). Gene Ontology (GO) and Kyoto Encyclopedia of Genes and Genomes (KEGG) pathway enrichment analyses were performed using clusterProfiler R packages.

**Human plasma sample collection**

A total of 213 subjects undergoing evaluation for coronary artery disease were enrolled in this study. Among these, 141 patients scheduled for coronary angiography were included, being composed of 34 patients with acute myocardial infarction (AMI), 78 patients with CHD, and 29 patients without CHD (Non-CHD). Plasma samples were obtained from individuals scheduled for coronary angiography at the Second Affiliated Hospital of Guangdong Medical University, China. sEVs were extracted as described above, and MPC1 levels in plasma and sEVs were measured by ELISA. Inclusion criteria were age >18 years and first-time suspicion of coronary heart disease. Exclusion criteria encompass the following: severe valvular heart disease; hemodynamic instability necessitating mechanical support; a history of heart failure or myocardial infarction; acute cerebrovascular accident, or severe hepatic or renal dysfunction; concurrent severe infection, systemic inflammatory disease, or autoimmune disease; significant bleeding tendency or recent history of hemorrhagic disease; pregnancy or lactation; refusal to provide informed consent, or participation in another clinical trial within the preceding 3 months; and a diagnosis of chronic obstructive pulmonary disease. All participants provided written informed consent. The study received approval from the Ethics Committee of the Second Affiliated Hospital of Guangdong Medical University (PJKT2024-163), in accordance with the ethical guidelines of the 1975 Declaration of Helsinki. Patient demographics and clinical characteristics are summarized in Table S10.

**Plasma sEVs extraction**

Blood samples were collected in EDTA tubes and centrifuged at 2,000 × g for 10 min at 4°C to obtain plasma. Plasma sEVs were isolated using both ultracentrifugation approach and a commercial extraction kit (Geneseed Biotech, Guangzhou, China). Based on previous reports[16,17], the ultracentrifugation procedure commenced with a centrifugation step at 2,500 × g for 15 minutes at 20°C to collect the supernatant, then centrifuged at 10,000 × g for 30 minutes at 20°C. The final step involved two rounds of ultracentrifugation of the supernatant at 120,000 × g for 2 h and 30 min each at 20°C to collect sEVs. The kit applied sequential centrifugation steps (2,000 × g, 10,000 × g, and 13,000 × g) and proprietary solutions per the manufacturer’s protocol. Plasma sEVs isolated from both protocols were used for subsequent analysis.

**Data analysis and statistics**

Experimental groups were randomly assigned. For both in vitro and in vivo experiments, at least five biologically independent replicates were performed. Data are presented as mean ± SEM for normally distributed data or as median (IQR) as appropriate. Each dataset was first tested for normality typically with Shapiro-Wilk test. Parametric or nonparametric analyses were then selected accordingly. For normally distributed data, differences between two groups were analyzed with an unpaired two-tailed Student's T-test. Comparisons among more than two groups were performed using one-way ANOVA with the Dunnett multiple-comparisons test. If the data were not normally distributed, the Mann-Whitney U test was used for comparisons between two groups, and Kruskal-Wallis ANOVA followed by Dunn's multiple-comparisons test was used for multiple-group comparisons. The homogeneity of variance was assessed using the Brown-Forsythe test. When the data exhibited heteroscedasticity, the Welch test was subsequently employed. When two independent factors were involved, two-way ANOVA combined with Tukey multiple comparisons test was used. Mixed-effects ANOVA was used when analyzing serial echocardiographic measurements taken at different time points. The details of the statistical analyses and biological replicate numbers for each figure can be found in the accompanying figure legends. For the survival curve rate, the Kaplan-Meier method and compared by the log-rank test was used. All data were analyzed with GraphPad Prism software (version 9.0) using the appropriate statistical analysis methods as indicated in the figure legends. Significance was accepted at $P < 0.05$.

For clinical analysis, categorical variables are presented as counts and percentages and were compared using the Chi-square or Fisher's exact test as appropriate. Continuous variables are presented as mean ± standard deviation or median (interquartile range). The normality of the data was assessed using the Shapiro-Wilk test. Differences between groups were analyzed using one-way

ANOVA with the Tukey multiple comparisons test or the Kruskal-Wallis test with the Dunnett multiple comparisons test for continuous variables. The Chi-Square test ($\chi 2$) or Fisher's exact test was employed for categorical variables. All statistical tests were two-sided, and *P* value < 0.05 was considered statistically significant. Statistical analyses were performed using SPSS 27.0 (IBM).

**Data Availability Statement**

The data, analytic methods, and study materials supporting the findings of this study are available to other researchers from the corresponding author upon reasonable request. RNA sequencing (RNA-seq) raw data have been deposited at the National Center for Biotechnology Information (BioProject: PRJNA1262715). Mass spectrometry proteomics data have been deposited to the Proteome Xchange Consortium (https://proteomecentral.proteomexchange.org) with the dataset identifier PXD064000 and PXD063946.

### Supplementary Tables

**Table S1. Antibody information.**

| Antibodies | Dilution | Clone/Catalog | Supplier |
|---|---|---|---|
| Bax | 1:1000 | 2772S | Cell Signaling Technology, USA |
| Bcl2 | 1:1000 | 26593-1-AP | Proteintech, USA |
| C-Cas3 | 1:1000 | 9661S | Cell Signaling Technology, USA |
| MPC1 | 1:1000 | DF3852 | Affinity Biosciences, China |
| MPC1 | 1:1000 | 30868-1-AP | Proteintech, USA |
| PCK2 | 1:1000 | DF12686 | Affinity Biosciences, China |
| ATP5A | 1:1500 | DF3806 | Affinity Biosciences, China |
| UQCRC2 | 1:1500 | DF12339 | Affinity Biosciences, China |
| SDHB | 1:1500 | DF12732 | Affinity Biosciences, China |
| NDUFB8 | 1:1500 | DF9666 | Affinity Biosciences, China |
| MTCO1 | 1:1500 | DF8920 | Affinity Biosciences, China |
| Collagen I | 1:1000 | ab26043 | Abcam, UK |
| Fibronectin | 1:1000 | ab45688 | Abcam, UK |
| α-SMA | 1:1000 | ab124964 | Abcam, UK |
| TSG101 | 1:1000 | ab125011 | Abcam, UK |
| CD9 | 1:1000 | 98327 | Cell Signaling Technology, USA |
| GM130 | 1:1000 | 66662-1-Ig | Proteintech, USA |
| CD63 | 1:1000 | ab315108 | Abcam, UK |
| TBP | 1:1000 | 22006-1-AP | Proteintech, USA |
| Tubulin | 1:5000 | 11224-1-AP | Proteintech, USA |

| Antibodies | Dilution | Clone/Catalog | Supplier |
|---|---|---|---|
| GAPDH | 1:5000 | AP0063 | Bioworld, China |
| Troponin T | 1:200 | ab209813 | Abcam, UK |
| CD45.1-FITC | 1:100 | 11-0453-81 | eBioscience, China |
| CD45.2-PE | 1:100 | 11-0454-85 | eBioscience, China |
| Anti-Mouse-HRP | 1:5000 | SA00001-1 | Proteintech, USA |
| Anti-Rabbit-HRP | 1:5000 | SA00001-4 | Proteintech, USA |
| Troponin I | 1:200 | ab47003 | Abcam, UK |
| Anti-Rabbit -Alexa488 | 1:500 | ab150077 | Abcam, UK |
| Anti-Mouse-Alexa488 | 1:500 | ab150113 | Abcam, UK |
| Anti-Mouse-Alexa 647 | 1:500 | ab150115 | Abcam UK |

**Table S2. Real-Time PCR sequences of primers.**

| Genes | Species | Sequences (5′-3′) |
|---|---|---|
| Gapdh | Mouse | F: TGACAACTCCCTCAAGATTGTCA |
| | | R: GGCATGGACTGTGGTCATGA |
| IL-6 | Mouse | F: AGTCCGGAGAGGAGACTTCA |
| | | R: ATTTCCACGATTTCCCAGAG |
| IFN-γ | Mouse | F: CACGCCGCGTCTTGGT |
| | | R: TCTAGGCTTTCAATGAGTGTGCC |
| TNF-α | Mouse | F: CACCACCATCAAGGACTCAA |
| | | R: AGGCAACCTGACCACTCTCC |
| IL-10 | Mouse | F: GACCAGCTGGACAACATACTGCTAA |
| | | R: GATAAGGCTTGGCAACCCAAGTAA |
| Ndufs2 | Mouse | F: CGGTGCCATGACTCCTTTCT |
| | | R: TCCGCTGTTACAACCCCAAT |
| Ndufs1 | Mouse | F: CCAGGAGGTTCTTGCTGACC |
| | | R: TGTCTCAGCATATGGACGGC |
| Sdhc | Mouse | F: TCCAGGCCGGAACTCAAGAT |
| | | R: CAGCCAGACCTGGGGTATTG |
| Sdhb | Mouse | F: TGGTCAGACCCGCTTATGTG |
| | | R: GGTCCAGTGGAGAGATGCAG |
| Uqcrc1 | Mouse | F: AAGTGCCTGCCTACGAGTTC |

| | | |
|---|---|---|
| | | R: TGTCCCCACCTTTTCTCAAGT |
| Uqcrfs1 | Mouse | F: GAGGTGACTCGGGGCAAAAA |
| | | R: GGAAGGACAACACAGTTCTCAAT |
| Cox7b | Mouse | F: CAGCCTTTCCAGGGATGAGAA |
| | | R: AGAAGCCCTGAATGGGGTTG |
| Cox4il | Mouse | F: TGGAACATGTCCCCTGTTGG |
| | | R: CACCATGCTTGGGCTTCATT |
| Atp5po | Mouse | F: GGTTCATCCTGCCAGAGACTA |
| | | R: AATCCCTCATCGAACTGGACG |
| Atp5f1b | Mouse | F: ATCTGTGGTCAGGCCCTTTG |
| | | R: CCAGAGACACTTTGGGGTCC |
| ANP | Mouse | F: ACCTGCTAGACCACCTGGAG |
| | | R: CCTTGGCTGTTATCTTCGGTACCG |
| BNP | Mouse | F: GAGGTCACTCCTATCCTCTGG |
| | | R: GCCATTTCCTCCGACTTTTCTC |

**Table S3. Reagent materials information.**

| Reagent materials | Clone/Catalog | Supplier |
|---|---|---|
| Enhanced BCA Protein Assay Kit | P0010S | Beyotime, China |
| Collagenase Type I | C1-28 | Sigma-Aldrich, USA |
| Collagenase Type II | C2-22 | Sigma-Aldrich, USA |
| Centrifuge tube | 788211 | Biotechnology, China |

| | | |
|---|---|---|
| Ultracentrifugation tube | 355618 | Beckman Coulter, USA |
| TUNEL Apoptosis Detection Kit | 40306ES60 | Yeasen, China |
| Trichrome Stain Kit | ab150686 | Abcam, UK |
| Human MPC1 ELISA kit | SMK0490HA | MEIKE, China |
| Mouse MPC1 ELISA kit | LBK-M02891 | LIANBOKEBio, China |
| Oxygen Consumption Rate (OCR) Plate Assay Kit | E297 | Dojindo, Japan |
| WGA Conjugate | I3310 | Solarbio, China |
| Mitochondrial extraction kit | C3601/C3606 | Beyotime, China |
| Cell mitochondrial separation reagent | C3601 | Beyotime, China |
| Tissue mitochondrial separation reagent | C3606 | Beyotime, China |
| GW4869 | HY-19363 | MCE, USA |
| DiR (Synonyms: Cy7 DiC18) | HY-D1048 | MCE, USA |

| Reagent materials | Clone/Catalog | Supplier |
|---|---|---|
| PKH26 Red Fluorescent Cell Linker Mini Kit | MINI26-1KT | Sigma-Aldrich, USA |
| 5% (v/v) bovine serum albumin | Cat#10837091001 | Roche, Switzerland |
| DAPI Staining Solution | Ab228549 | Abcam, UK |

**Table S4. Statistical analysis of mitochondrial energy metabolism-related differentially expressed proteins listed in Veen diagram (Figure 7A).**

| Name (Unipro ID) | Test set | LogFC. (AMI vs Sham) | *P*-value* |
|---|---|---|---|
| Mpc1 (P63030) | sEVs | 3.173149631 | 0.00516739 |
| | Plasma | 2.39 | 0.000413 |
| Ndufa5 (Q9CPP6) | sEVs | 1.122866964 | 0.02386223 |
| | Plasma | 2.52 | 0.000315 |
| Ndufa1 (O35683) | sEVs | 0.686165379 | 0.03488619 |
| | Plasma | 1.49 | 0.02252 |
| Ndufs2 (Q91WD5) | sEVs | 0.727611981 | 0.0008817 |
| | Plasma | 1.35 | 0.000117 |
| Ndufs1 (Q91VD9) | sEVs | 0.825125752 | 0.03921296 |
| | Plasma | 0.89 | 0.002094 |
| Pck2 (Q8BH04) | sEVs | -2.765588519 | 0.00195796 |
| | Plasma | -1.45 | 0.002437 |

**Table S5. The physical and biochemical parameters of patients.**

| Variables | Non-CHD (n= 29) | CHD (n= 78) | AMI (n= 34) |
|---|---|---|---|
| Age, years | 59.66 ± 11.54 | 65.36 ± 11.10* | 65.91 ± 13.28 * |
| Male, % | 51.7 (15) | 57.7 (45) | 58.8 (20) |
| Smoking, % | 37.9 (11) | 50.0 (39) | 67.6 (23) |

| | | | |
|---|---|---|---|
| Drinking, % | 27.6 (8) | 42.3 (33) | 52.9 (18) |
| HTN, % | 44.8 (13) | 60.3 (47) * | 38.2 (13) |
| HLP, % | 48.3 (14) | 64.1 (50) * | 58.8 (20) |
| DM, % | 6.9 (2) | 14.1 (11) | 32.4 (11) |
| BW, kg | 65.00 ± 14.12 | 63.89 ± 10.54 | 65.27 ± 13.13 |
| HR, Times/min | 77.66 ± 13.49 | 78.69 ± 12.34 | 83.65 ± 21.70 |
| DBP, mmHg | 87.07 ± 14.40 | 88.21 ± 17.80 | 84.85 ± 15.25 |
| SBP, mmHg | 136.34 ± 20.72 | 160.92 ± 20.49 | 139.62 ± 27.03 |
| EF, % | 62.17 ± 7.40 | 61.55 ± 5.82 | 50.68 ± 11.53 *,# |
| FS, % | 34.38 ± 5.52 | 32.67 ± 3.76 | 27.94 ± 6.98* |
| LVIDd, mm | 46.52 ± 5.15 | 44.78 ± 5.38 | 48.06 ± 7.34# |
| LVIDs, mm | 31.31 ± 4.58 | 30.59 ± 4.62 | 33.71 ± 6.48# |
| hs-CRP, mg/L | 3.87(0.25-30.50) | 4.14(0.01-18.74) | 6.33(0.32-15.81) # |
| NT-proBNP, ng/L | 53.29 (10-1892) | 84.71(18-1891) | 332.1(28-13700) *,# |
| hs-TnT, μg/ L | 4.15(0.03-17.68) | 7.34(3.19-26.30) | 17.85(3.60-5434.00) *,# |
| LDH, U/L | 209.76(133-338) | 197.85(4.38-291) | 323(122-1630) *,# |
| CKMB, U/L | 17.43(8.90-76.50) | 15.75(0.46-43.40) | 39.72(1.60-237.80) *,# |
| CK, U/L | 154.14(31-589) | 117.54(27-397) * | 148(31-2517) # |
| α-HBDH, U/L | 150 (102-247) | 146(1.37-244) | 156(100-1526) *,# |
| HbA1c.% | 5.56(3.70-7.60) | 6.85(3.70-16.40) | 6.94(4.00-12.20) * |
| GLU, mmol/L | 6.65(4.80-10.39) | 8.08(3.99-27.56) | 8.54(4.66-16.70) |
| TC, mmol/L | 4.90(2.99-7.36) | 5.24(1.62-9.37) | 5.38(2.79-8.78) * |

| | | | |
|---|---|---|---|
| LDL-C, mmol/L | 3.28(1.54-5.41) | 3.51(0.32-7.34) | 3.74(1.56-6.94) |
| HDL-C, mmol/L | 1.27(0.83-1.88) | 1.38(0.81-2.63) | 1.29(0.79-2.04) |
| UREA, mmol/L | 5.90(3.20-29.70) | 5.46(2.80-10.00) | 6.36(3.20-13.90) |
| ALT, U/L | 27.80(10.00-148.40) | 22.70(7.80-72.00)$^{*}$ | 25.42(7.80-81.60) |
| EGFR, mL /min | 91.57(63.27-119.32) | 83.89(41.55-119.32) $^{*}$ | 76.97(26.37-106.44) $^{*}$ |
| WBC, $10^9$/L | 6.97(4.37-10.82) | 7.10(3.48-13.87) | 8.29(4.36-19.84) $^{*}$ |
| RBC, $10^9$/L | 4.77(3.46-6.38) | 4.50(3.03-5.17) $^{*}$ | 4.53(2.85-6.68) |
| NEU, $10^9$/L | 4.66(2.14-8.51) | 4.64(1.82-11.94) | 4.95(2.14-11.99) $^{\#}$ |
| PLT, $10^9$/L | 265 (155-399) | 257(148-364) | 235(115-416) |
| HGb, g/L | 136 (92-160) | 132(92-160) | 131(85-158) |

Statistical analyses were performed using was performed using the one-way ANOVA with the Tukey multiple comparisons test or the Kruskal Wallis with the Dunn multiple comparisons test for continuous variables and χ2 or Fisher exact test for categorical variables. $^{*}$, CHD/AMI vs Non-CHD, $P$<0.05; $^{\#}$ AMI vs CHD, $P$<0.05.

Abbreviations: CHD, coronary atherosclerotic heart disease; Non-CHD , AMI, acute myocardial infarction; HTN, hypertension; HLP, hyperlipidemia; DM, diabetes mellitus; BW, body weight; SBP, systolic blood pressure; DBP, diastolic blood pressure; EF, left ventricular ejection fraction; FS, left ventricular fraction shortening; LVIDs, left ventricular internal diameter in systole; LVIDd, left ventricular internal diameter in diastole; hs-CRP, high sensitivity; C-reactive protein; NT-pro-BNP, N-terminal pro-brain natriuretic peptide; hs-TnT, high sensitivity troponin T; LDH, lactate dehydrogenase; CK-MB, creatine kinase MB isoenzyme; HbA1c, hemoglobin A1c; TC, total cholesterol; HDL-c, high-density lipoprotein cholesterol; LDL-c, low-density lipoprotein cholesterol; CREA, creatinine; ALT, Alanine Aminotransferase; EGFR, Estimated glomerular filtration rate; WBC, white blood cell; RBC, red blood cell; NEU, Neutrophil count; PLT, platelet; HGb, Hemoglobin.

**Table S6. Echocardiographic parameters at day 7 post AMI (For Figure 1B).**

| | Sham | Sham+Spl | AMI | AMI+Spl |
|---|---|---|---|---|
| HR (bpm) | 451.0±9.4 | 455.6±9.5 | 460.6±9.3 | 463.7±10.2 |
| EF (%) | 69.3±3.7 | 75.4±1.6 | 32.8±3.4*** | 20.4±3.6# |
| FS (%) | 43.3±1.9 | 42.7±1.4 | 18.5±1.8*** | 12.1±1.5# |
| LVIDs (mm) | 1.4±0.2 | 1.5±0.3 | 5.9±0.1*** | 6.4±0.5 |
| LVIDd (mm) | 3.0±0.3 | 3.3±0.5 | 6.9±0.4* | 7.2±0.2 |
| LVESV (μL) | 24.5±3.5 | 28.9±2.9 | 58.4±4.3** | 64.5±3.6 |
| LVEDV (μL) | 68.3±4.3 | 70.8±3.8 | 84.5±5.1 | 99.3±4.9# |

Statistical analyses were performed using the one-way ANOVA with the Tukey multiple comparisons test (HR, LVIDs and LVESV), the Brown-Forsythe and Welch ANOVA test with the Dunnett T3 multiple comparisons test (EF, FS, LVIDd, LVIDs, LVESV, and LVEDV). *, AMI vs Sham, $P<0.05$; **, AMI vs Sham, $P<0.01$; ***, AMI vs Sham, $P<0.001$; #, AMI+Spl vs AMI, $P<0.05$; n=6/group.

Abbreviations: HR, heart rate; EF, ejection fraction; FS, fraction shortening; LVIDs, left ventricular internal diameter in systole; LVIDd, left ventricular internal diameter in diastole; LVESV, Left ventricular end-diastolic volume; LVEDV, Left ventricular end-systolic volume.

**Table S7. Echocardiographic parameters at day 7 after AMI (For Figure 3B).**

| | AMI | AMI+S-sEVs | AMI+M1D-sEVs | AMI+M3D-sEVs | AMI+M7D-sEVs |
|---|---|---|---|---|---|
| HR (bpm) | 459.0±8.9 | 446.6±9.3 | 442.4±6.5 | 450.4±8.5 | 472.4±7.5 |
| EF (%) | 24.17±2.5 | 24.25±3.9 | 24.89±2.3 | 37.04±3.6* | 26.16±2.6 |
| FS (%) | 17.5±2.6 | 15.1±2.1 | 14.9±2.0 | 28.6±2.2** | 17.4±2.3 |
| LVIDs (mm) | 5.7±0.3 | 6.2±0.5 | 5.3±0.2 | 4.3±0.6** | 5.7±0.4 |
| LVIDd (mm) | 3.3±0.2 | 3.5±0.1 | 2.9±0.4 | 1.8±0.4* | 2.9±0.6 |
| LVESV (μL) | 57.9±5.4 | 59.8±6.3 | 56.2±4.7 | 34.5±5.1** | 58.3±4.2 |
| LVEDV (μL) | 85.2±2.2 | 86.7±3.7 | 88.5±2.3 | 82.9±4.7 | 88.6±2.1 |

Statistical analyses were performed using the one-way ANOVA with the Tukey multiple comparisons test (HR, LVIDd and LVESV), the Brown-Forsythe and Welch ANOVA test with the Dunnett T3 multiple comparisons test (EF, LVIDs, LVESV, and LVEDV) and the Kruskal Wallis with the Dunn multiple comparisons test (FS). *, vs AMI, $P<0.05$; **, vs AMI, $P<0.01$; n=6/group.

**Table S8. Echocardiographic parameters of host mice at day 7 post AMI (For Figure 4E).**

| | Host AMI/No Spl | Host AMI/Spl |
|---|---|---|
| HR (bpm) | 463.6±10.3 | 467.0±9.6 |
| EF (%) | 23.2±1.9 | 26.8±2.0 |
| FS (%) | 17.7±1.5 | 17.3±1.1 |
| LVIDs (mm) | 2.7±0.4 | 2.9±0.1 |
| LVIDd (mm) | 4.9±0.3 | 5.2±0.3 |
| LVESV (µL) | 87.1±7.5 | 89.3±8.4 |
| LVEDV (µL) | 115±9.9 | 109±13.2 |

Statistical analyses were performed using the student's t-test among groups. n=5-6/group.

**Table S9. Echocardiographic parameters at day 7 post AMI (For Figure 5C).**

| | DMSO | GW4869 | GW4869+S-sEVs | GW4869+M3D-sEVs |
|---|---|---|---|---|
| HR (bpm) | 456.2±9.8 | 450.6±9.7 | 465.3±9.2 | 463.2±10.2 |
| EF (%) | 25.3±2.3 | 17.5±3.4** | 19.4±2.8 | 24.86±2.5$^{\$}$ |
| FS (%) | 23.1±1.5 | 12.6±1.4*** | 10.61±1.6 | 18.81±1.9$^{\$\$}$ |
| LVIDs (mm) | 6.3±0.4 | 7.8±1.2* | 7.2±0.6 | 6.7±0.5$^{\$}$ |
| LVIDd (mm) | 3.6±0.6 | 5.2±0.4** | 6.1±0.7$^{\#}$ | 4.2±0.2$^{\$\$}$ |
| LVESV (µL) | 58.8±5.1 | 69.8±4.3* | 71.2±2.9 | 63.5±4.4 |
| LVEDV (µL) | 85.7±3.2 | 95.5±7.3* | 103.6±5.2$^{\#\#}$ | 90.6±4.9 |

Statistical analyses were performed using the one-way ANOVA with the Tukey multiple comparisons test (EF, FS, LVIDd and LVESV), the Brown-Forsythe and

Welch ANOVA test with the Dunnett T3 multiple comparisons test (HR, LVIDs and LVEDV). *, GW4869 vs DMSO, $P$<0.05; **, GW4869 vs DMSO, $P$<0.01; ***, GW4869 vs DMSO, $P$<0.001; #, GW4869+S-sEVs vs GW4869, $P$<0.05; ##, GW4869+S-sEVs vs GW4869, $P$<0.01; ###, GW4869+S-sEVs vs GW4869, $P$<0.001; \$, GW4869+M3D-sEVs vs GW4689, $P$<0.05; \$\$, GW4869+M3D-sEVs vs GW4689, $P$<0.01; \$\$\$, GW4869+M3D-sEVs vs GW4689, $P$<0.001; n=6/group.

**Table S10. Echocardiographic parameters at day 7 post AMI (For Figure 8B).**

| | sEVs | sEVs+IgG | sEVs+Ab |
|---|---|---|---|
| HR (bpm) | 456.7±8.9 | 462.4±11.0 | 458.3±9.1 |
| EF (%) | 39.0±1.2 | 37.9±2.6 | 28.2±1.9** |
| FS (%) | 27.1±1.7 | 25.3±1.3 | 14.7±1.1*** |
| LVIDs (mm) | 4.2±0.3 | 4.0±0.1 | 5.4±0.3* |
| LVIDd (mm) | 1.8±0.3 | 2.1±0.2 | 3.1±0.1** |
| LVESV (μL) | 33.2±5.3 | 32.1±4.9 | 56.6±5.3** |
| LVEDV (μL) | 85.3±2.9 | 85.9±3.1 | 87.8±4.1 |

Statistical analyses were performed using the one-way ANOVA with the Tukey multiple comparisons test (HR, EF, FS and LVEDV), the Brown-Forsythe and Welch ANOVA test with the Dunnett T3 multiple comparisons test (LVIDs and LVESV) and the Kruskal Wallis with the Dunn multiple comparisons test (LVIDd). *, sEVs+Ab vs sEVs+IgG, $P$<0.05; **, sEVs+Ab vs sEVs+IgG, $P$<0.01; ***, sEVs+Ab vs sEVs+IgG, $P$<0.001; n=6/group.

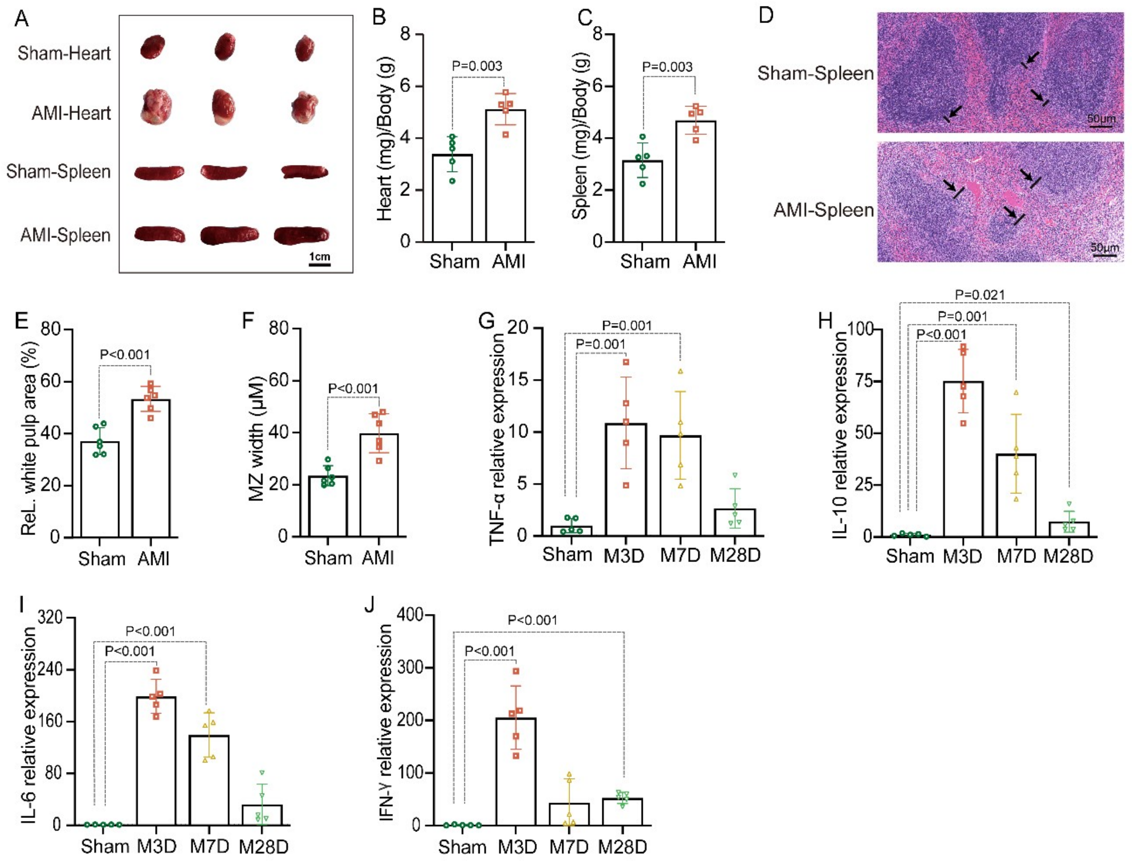


**Figure S1. Spleen activity is increased in AMI mice.**

**A**, Representative images of hearts and spleens in mice 3 days after AMI (n=6/group). **B**, Heart weights of sham and AMI mice (n=6/group). C. Spleen weights of sham and AMI mice (n=6/group). **D-F**, Representative H&E images of spleen slices from sham and AMI mice and quantitation of relative (ReL.) white pulp area and marginal zone 3 days after AMI (n=6/group). **G-J**, Cytokines mRNA levels by qPCR in spleen tissues from sham and AMI mice on day 3, 7, 28 post AMI. Data were normalized to GAPDH and expressed as folds of sham group (n=6/group). Statistical analyses were performed by unpaired t test (**B**, **C**, **E** and **F**), the one-way ANOVA with the Dunnett multiple comparisons test (**H** and **I**), and the Kruskal-Wallis with the Dunnett multiple comparisons test (**G** and **J**).

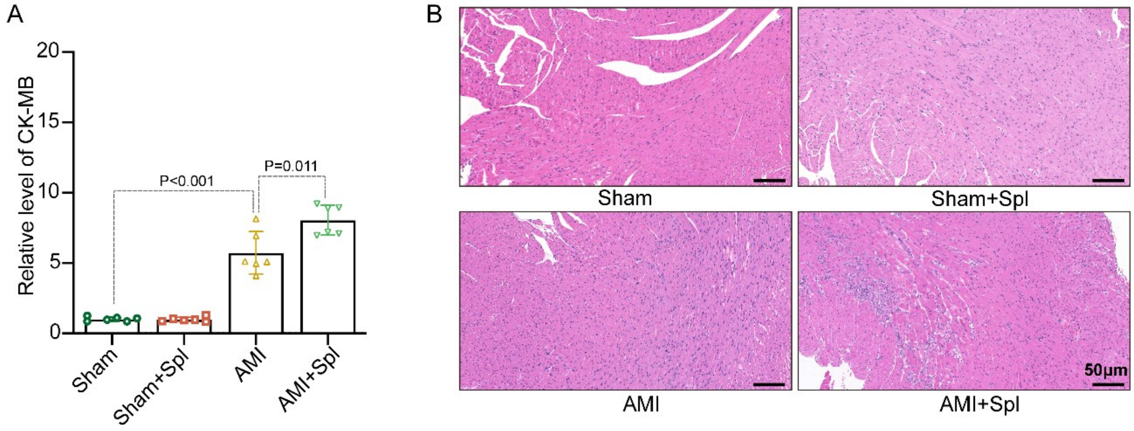


**Figure S2. Splenectomy aggravates cardiac injury in AMI mice.**

**A**, CK-MB levels in cardiac tissues of sham and AMI mice with or without splenectomy 28 days after AMI (n=6/group). **B**, Representative H&E images in sham and AMI mice with or without splenectomy 14 days after AMI (n=6/group).. Statistical analyses were performed using the Kruskal-Wallis with the Dunnett multiple comparisons test in **A**.

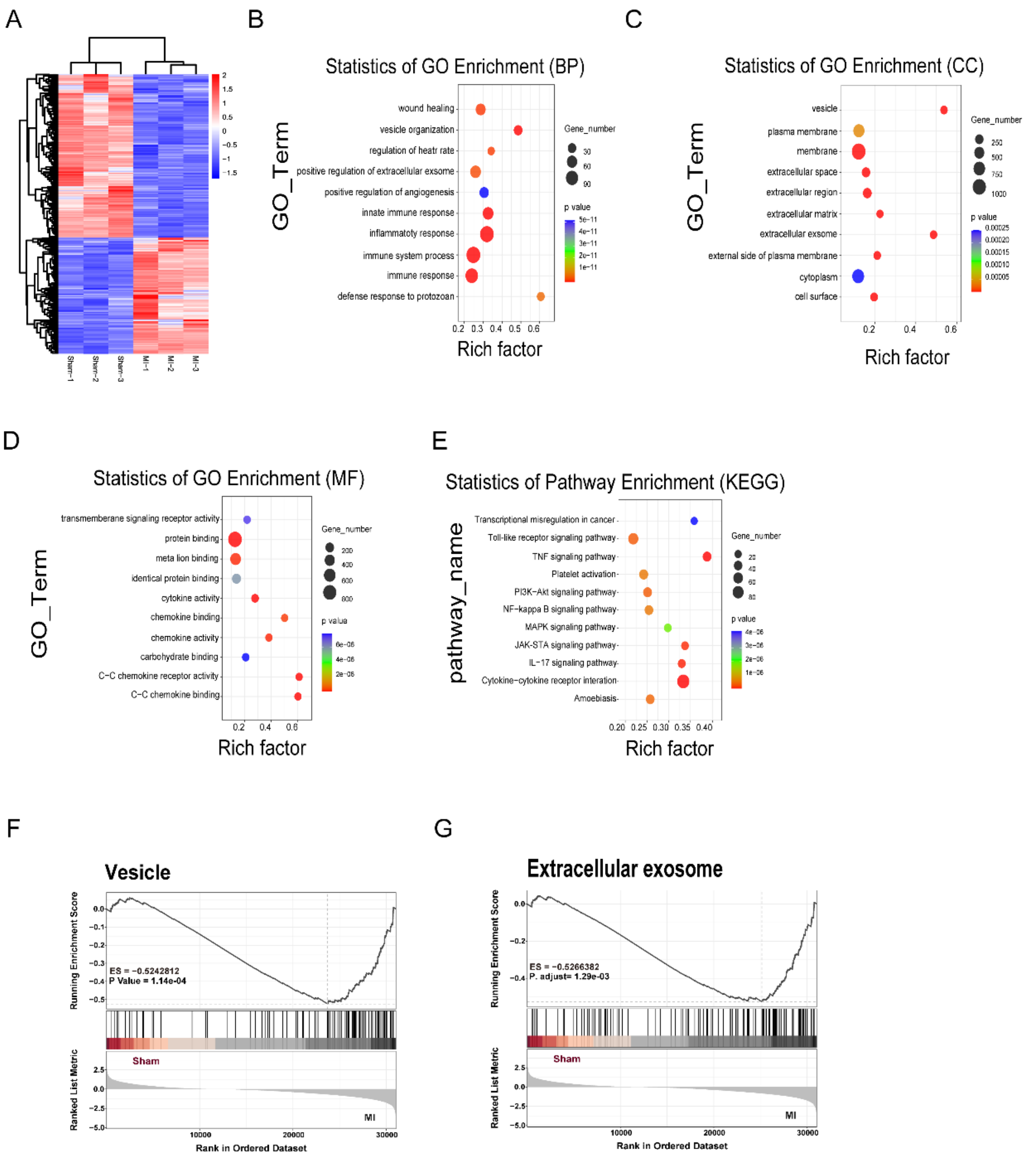


**Figure S3. Spleen-derived sEVs biogenesis is enhanced in spleens of AMI mice.**

**A**, Heatmap of the differentially expressed genes (DEGs) of spleen tissues between sham and AMI mice 1 day post AMI. **B-D**, GO enrichment analysis of DEGs. **E**, KEGG enrichment analysis of DEGs. **F-G**, GSEA analysis of vesicle or extracellular exosomes-related DEGs in spleens between sham and AMI mice.

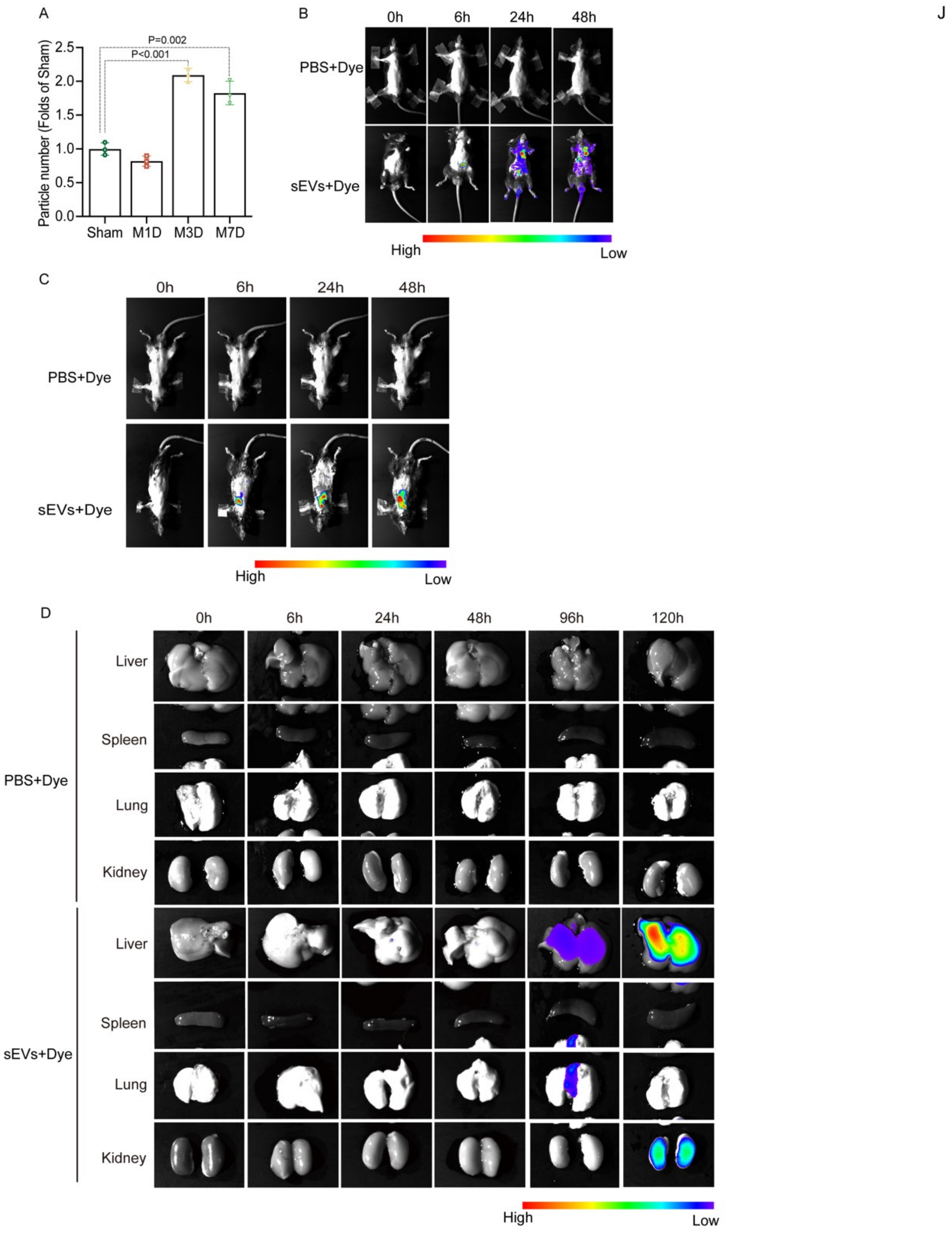


**Figure S4. Spleen-derived sEVs number is increased in response to AMI and these sEVs are predominantly captured by mouse hearts.**

**A**, Quantitative nFCM analysis for Figure 2H showing the abundance of Spleen-derived sEVs at different time points after myocardial infarction. P. concentration represents particle concentration. **B-C**, Biodistribution of DiR-labeled spleen-derived sEVs administrated with intraperitoneal injection (**B**) or intramyocardial injection (**C**) in mice. **D**, Images showed the biodistribution of DiR-labeled spleen-derived sEVs from AMI mice in recipient mice's liver, spleen, lung and kidney at indicated time-points after intramyocardial injection. Statistical analyses were performed by the one-way ANOVA with the Dunnett multiple comparisons test in **A**.

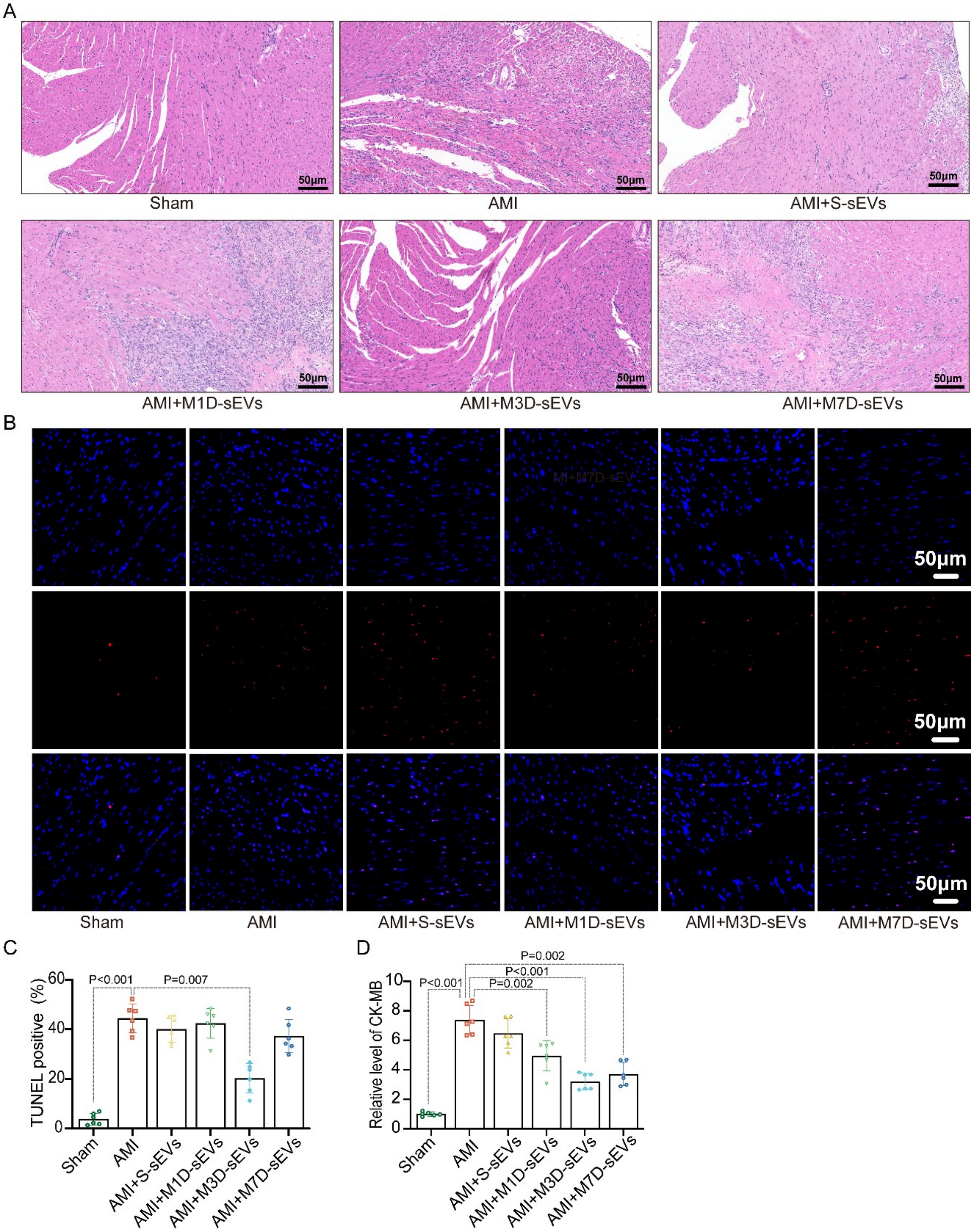


**Figure S5. Spleen-derived sEVs from AMI mice alleviate cardiac ischemic injuries in hearts from recipient AMI mice.**

**A**, Representative H&E images from sham and AMI mice treated as indicated 14 days after AMI (n=6/group). **B-C**, Representative TUNEL staining images and quantitation of TUNEL positive rates in hearts from sham and AMI mice treated as indicated 3 days after AMI (n=6/group). **D**, CK-MB levels in cardiac tissues from sham and AMI mice treated as indicated 28 days after AMI (n=6/group). Statistical analyses were performed by the one-way ANOVA with the Dunnett multiple comparisons test in **C** and **D**.

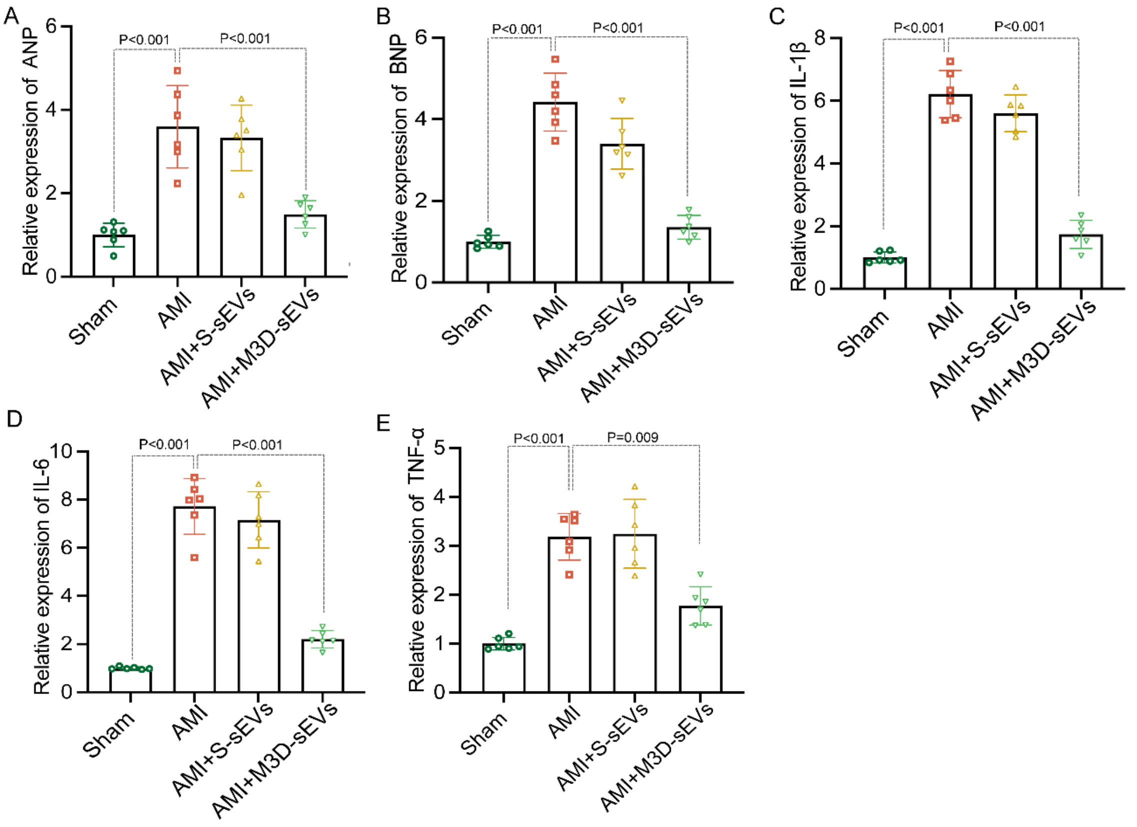


**Figure S6. M3D-sEVs reduce fibrotic gene expression and cytokines production in hearts from recipient AMI mice.**

**A-B,** ANP and BNP mRNA levels by qPCR in heart tissues 28 days post AMI (n=6/group). **C-E,** Cytokines levels by ELISA in cardiac tissues from sham and AMI mice treated as indicated 7 days after AMI (n=6/group). Statistical analyses were performed using the one-way ANOVA with the Dunnett multiple comparisons test (A, B and E), the Brown-Forsythe and Welch ANOVA test with the Dunnett T3 multiple comparisons test (C) and the Kruskal Wallis with the Dunn multiple comparisons test (D).

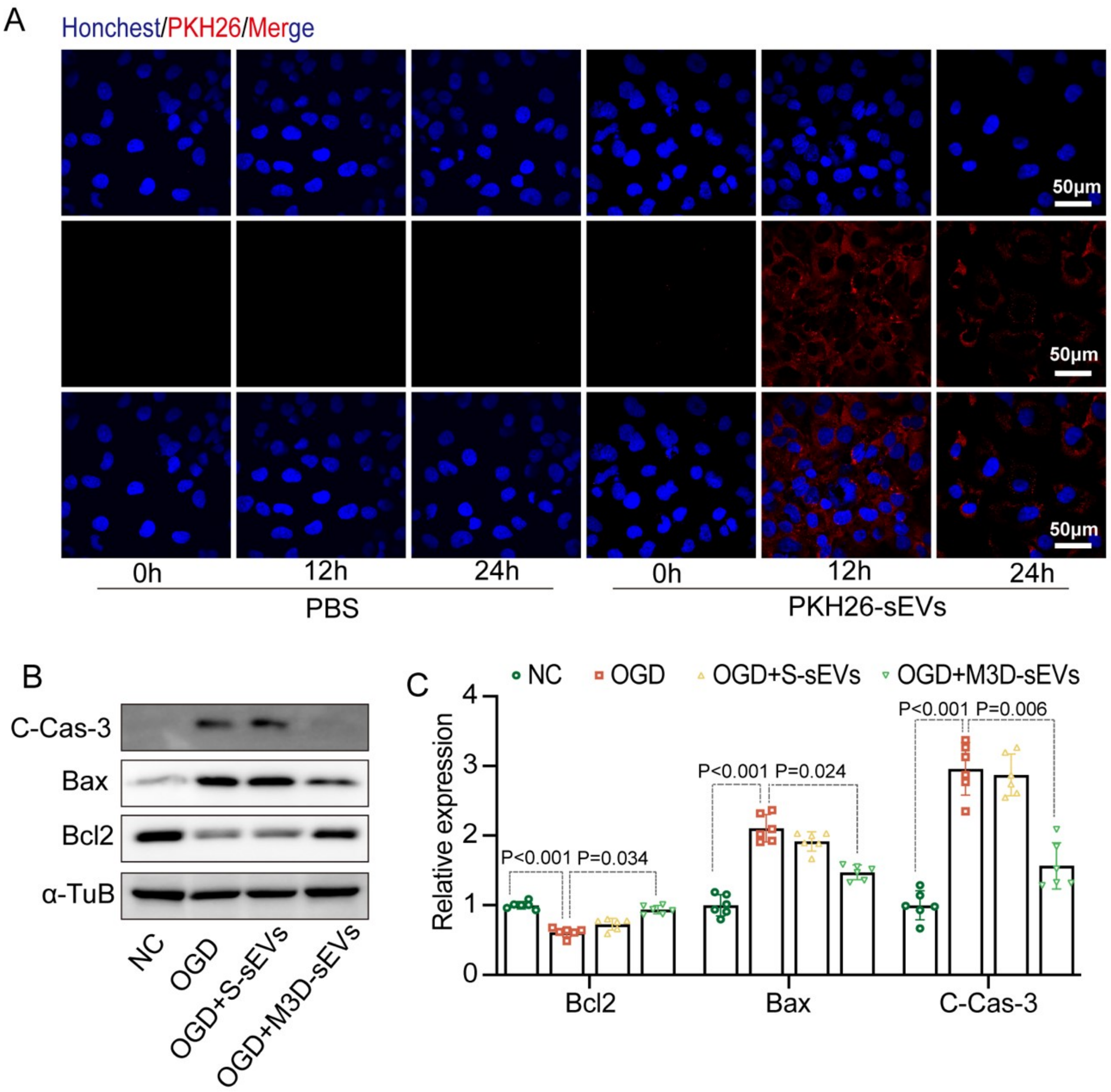


**Figure S7. Uptake of spleen-derived M3D-sEVs alleviates OGD-induced cardiomyocyte damage.**

**A**, Representative immunofluorescent images of PKH26-labeled M3D-sEVs inside H9C2 cells. PBS+PKH26 was used as control. **B** and **C**, Representative western blotting images and quantitation of C-Cas-3, Bax, and Bcl2 expression in NC or OGD-insulted H9C2 cells treated with S-sEVs, M3D-sEVs, or vehicles (n=6/group). Statistical analyses were performed by the one-way ANOVA with the Tukey multiple comparisons test in **C**.

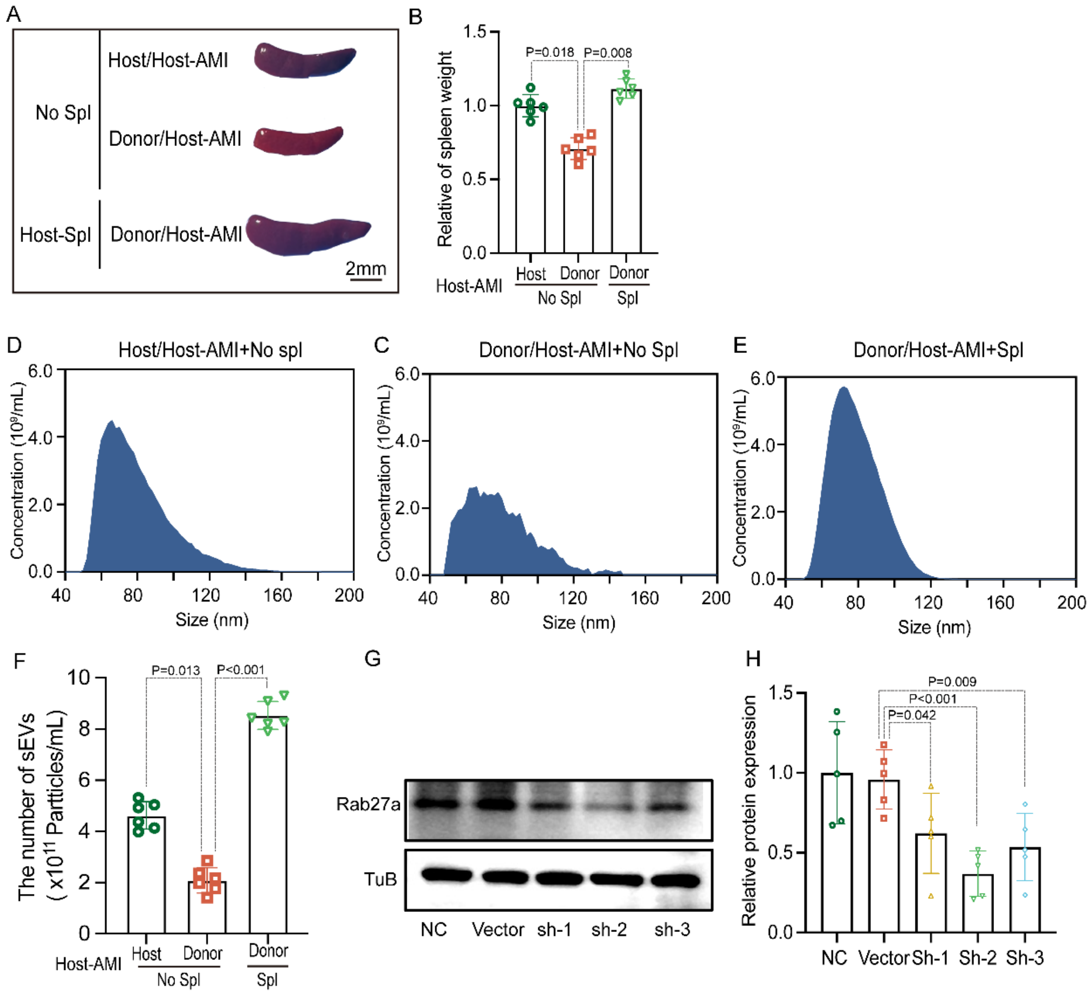


**Figure S8. Alterations of splenic M3D-sEVs release after AMI in parabiosis mice and shRab27a knockdown efficiency.**

**A**, Representative images of spleens from a parabiosis model, in which host and donor parabiont mice are joined, with AMI induced in the host. Spl indicates splenectomy. **B**, Relative weight of spleen (n=6/group, normalized to humerus length, folds of control group). **C-F**, Representative images of size distribution and concentration of M3D-sEVs (n=6/group). **G-H**, Western blotting results for testing the knockdown efficiency of AAV2-shRab27a in mouse hearts (n=6/group). Statistical analyses were performed using mixed-effects ANOVA followed by the Sidak multiple comparisons test (**H;** sh-1 and sh-3), the one-way ANOVA with the Dunnett multiple comparisons test (**B**, **F** and **H;** sh-2).

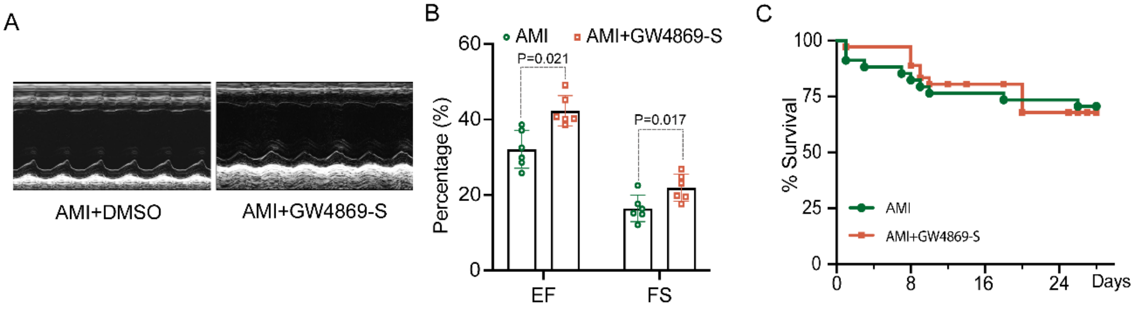


**Figure S9. GW4869-S alleviates cardiac dysfunction in AMI mice**

**A-B,** Representative echocardiographic images of AMI mice treated with GW4869-S 3 days post AMI and quantitation of EF% and FS% (n=6/group). **C，** Survival curves for AMI mice treated with GW4869-S (n=30/group). Statistical analyses were performed using the two-way ANOVA with the Tukey multiple comparisons test (**B**). Survival curve rate was analyzed by the Kaplan-Meier method and compared by the log-rank test in **C**.

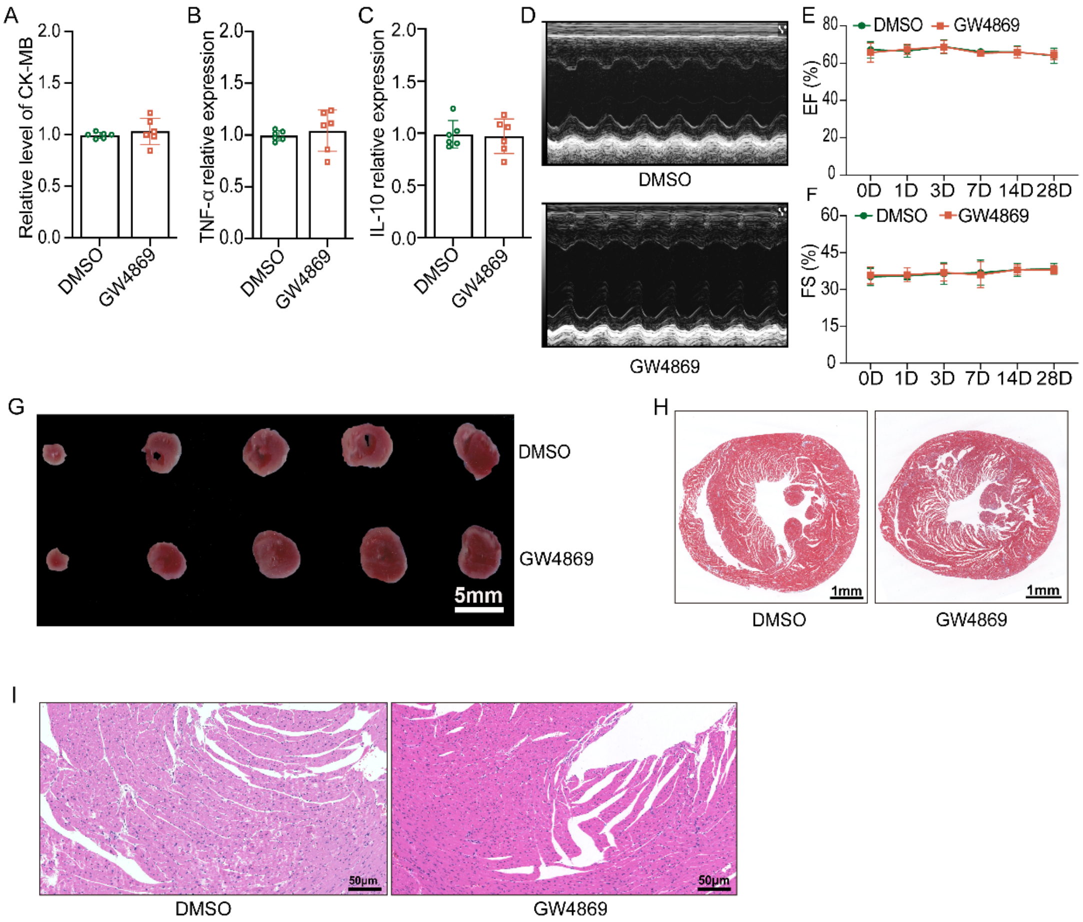


**Figure S10. GW4869 does not induce damage to hearts from normal mice.**

**A**, CK-MB levels in hearts from mice treated with GW4869 (2.5mg/kg) or equivalent vehicle (n=6/group). **B-C**, TNF-α and IL-10 mRNA levels in cardiac tissues from mice treated GW4869 or equivalent vehicle (n=6/group). **D-F**, Representative echocardiograms and quantitation of EF% and FS% in mice treated with GW4869 or equivalent vehicle (n=6/group). **G-I**, Representative images of TTC, Masson's trichrome, and H&E staining, respectively. Statistical analyses were performed with the Mann-Whitney U test (**A** and **B**), unpaired t test (**C**), and mixed-effects ANOVA followed by the Sidak multiple comparisons test (**E** and **F**).

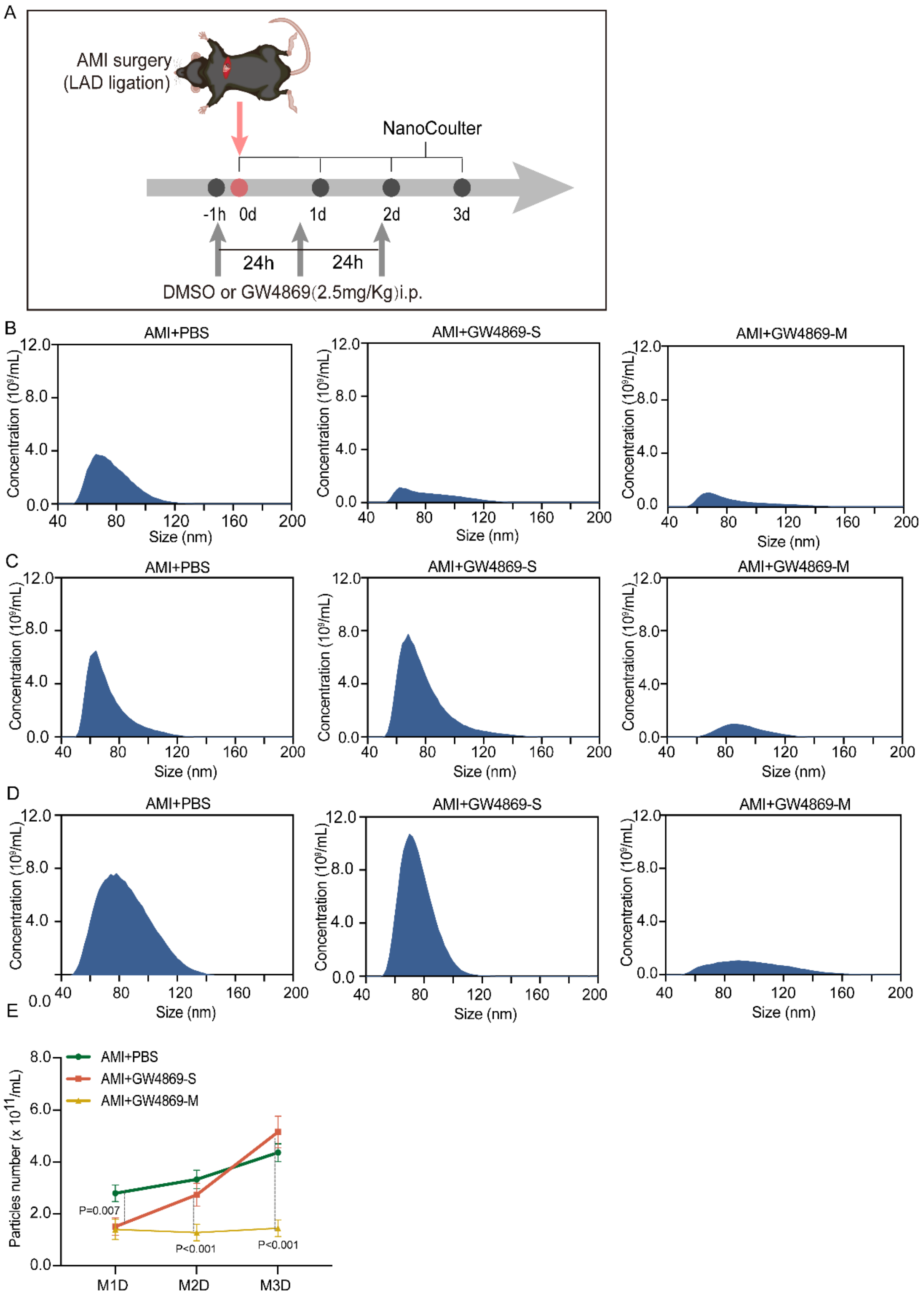


**Figure S11. Splenic sEVs release dynamics in acute myocardial infarction mice treated with GW4869-S/M.**

**A**, Flowchart for detecting the dynamics of splenic sEVs release in AMI mice treated with GW4869-S/M. **B**, Splenic sEVs size and concentration in mice at day 1 post AMI. **C**, Splenic sEVs size and concentration in mice at day 2 post AMI. **D**, Splenic sEVs size and concentration in mice at day 3 post AMI. **E**, Quantification of splenic sEVs concentration in mice for day 1 to 3 post AMI. Statistical analyses were performed using mixed-effects ANOVA followed by the Sidak multiple comparisons test in **E**.

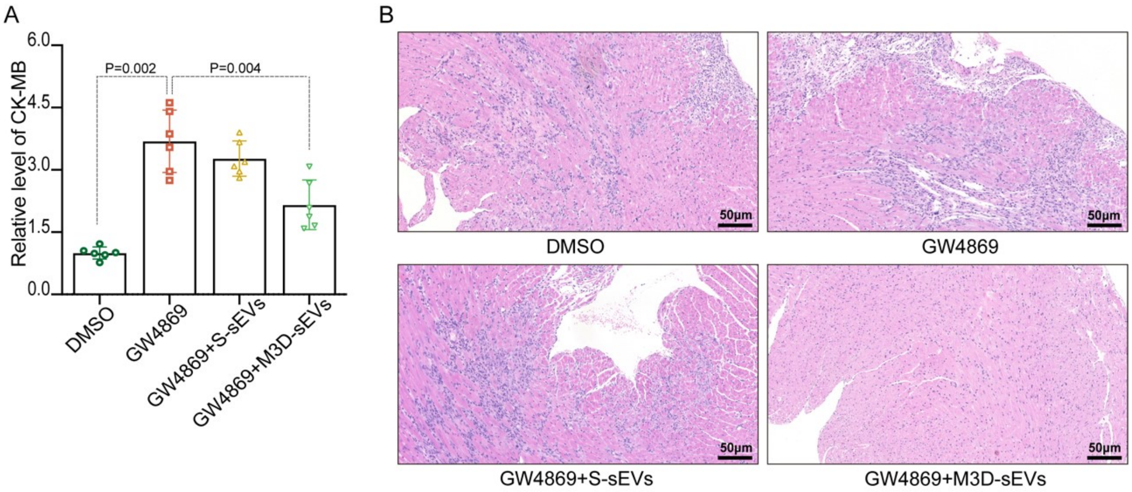


**Figure S12. GW4869-M aggravates myocardial injury in AMI mice.**

**A**, CK-MB levels in cardiac tissue 28 days after AMI. Contents of sEVs protein determined by BCA (n=6/group). **B**, Representative H&E images of cardiac sections from AMI mice treated GW4869-M plus S-sEVs, M3D-sEVs, or equivalent vehicles 7 days post AMI (n=6/group). Statistical analyses were performed with the Brown-Forsythe and Welch-ANOVA test with the Dunnett T3 multiple comparisons test (**A**).

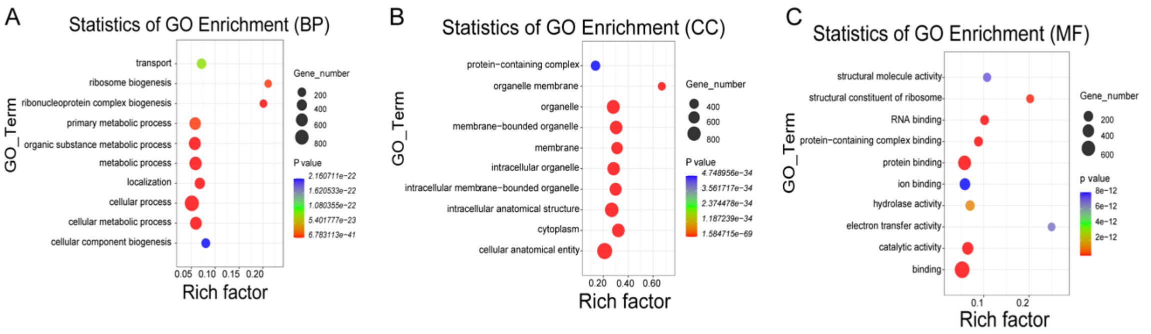


**Figure S13. GO enrichment analysis of spleen-derived sEVs between sham and AMI mice 3 days post AMI.**

**A-C**, BP, Biological Process. CC, Cellular Component. MF, Molecular Function.

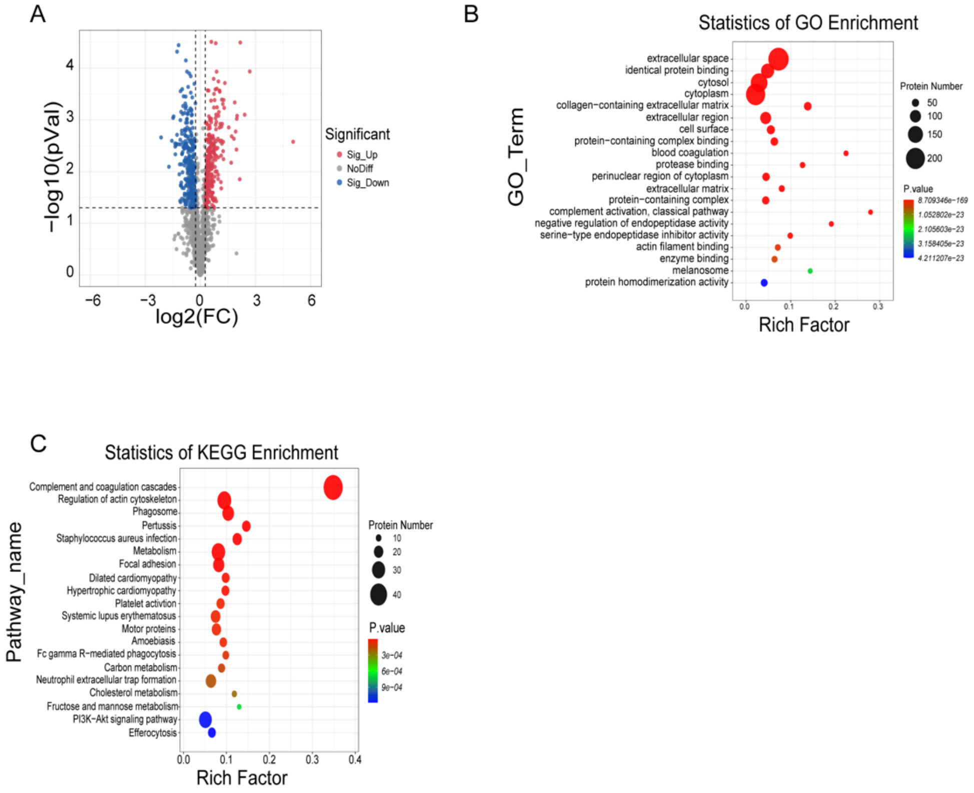


**Figure S14. Plasma proteomic profile of between sham and AMI mice.**

A, Volcano plot of differentially expressed proteins in plasma between sham and AMI mice 3 days after AMI. B, GO enrichment analysis of plasma differentially expressed proteins. C, KEGG enrichment analysis of plasma differentially expressed proteins.

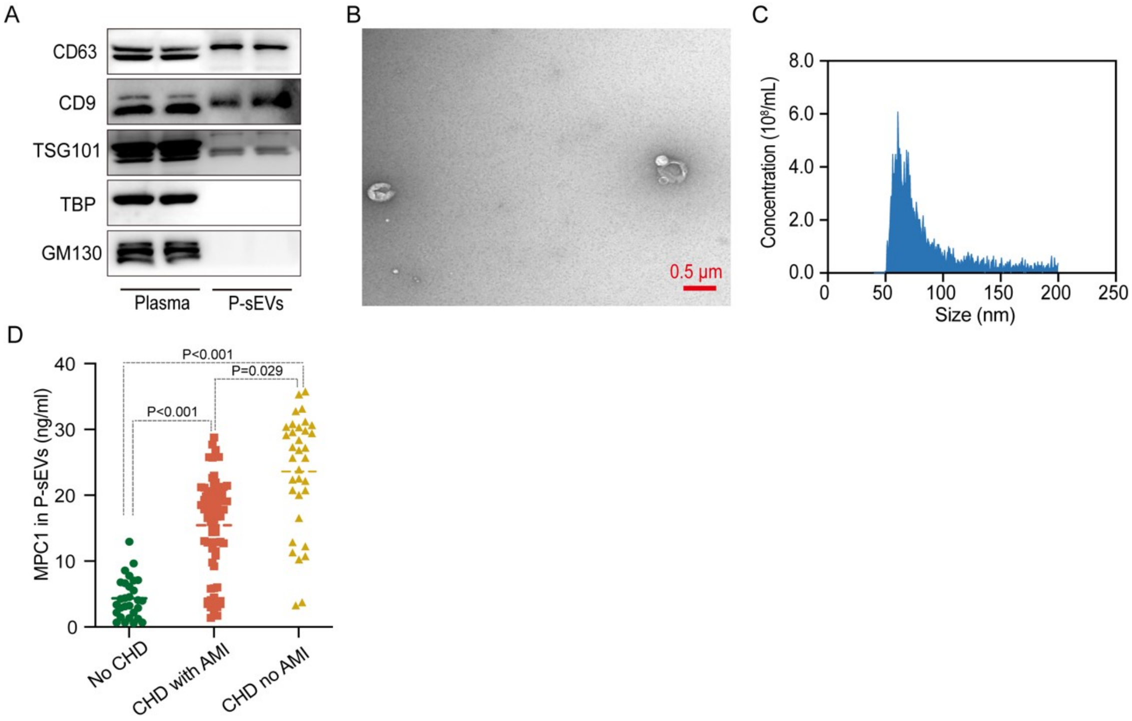


**Figure S15. MPC1 protein expression is increased in circulating plasma sEVs isolated using kits from patients.**

**A,** Western blotting results of P-sEVs markers. **B,** Representative TEM images of P-sEVs. **C**, Size distributions of P-sEVs measured by nFCM. **D**, ELISA assay results of MPC1 expression in P-sEVs among patient groups. Statistical analyses were performed by the one-way ANOVA with the Dunnett multiple comparisons test in **D**.

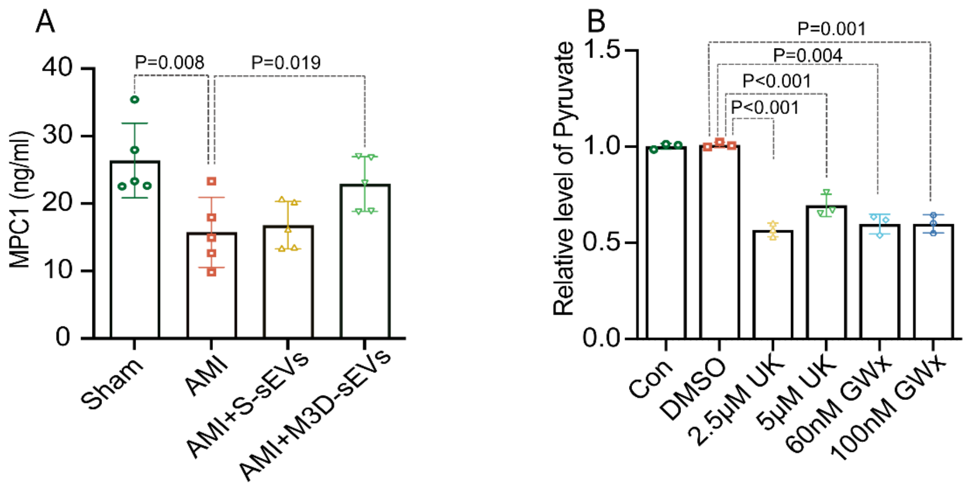


**Figure S16. MPC1 inhibitors suppressed mitochondrial pyruvate influx.**

**A,** MPC1 levels in myocardial tissues of sham, AMI, and AMI mice treated with sEVs or M3D-sEVs. **B**, Quantification of pyruvate levels in H9C2 cells treated with vehicles or MPC1 inhibitors with various concentrations. GWx indicates GW604714x; UK, UK5099. Statistical analyses were performed the one-way ANOVA with the Dunnett multiple comparisons test (**A-B**).

### References

1. Li S, Shen S, Xu H, Cai S, Yuan X, Wang C, et al. IGF2BP3 promotes adult myocardial regeneration by stabilizing MMP3 mRNA through interaction with m6A modification. *Cell Death Discov* 2023;9:164. doi: https://doi.org/10.1038/s41420-023-01457-3

2. Tian Y, French BA, Kron IL, Yang Z. Splenic leukocytes mediate the hyperglycemic exacerbation of myocardial infarct size in mice. *Basic Res Cardiol* 2015;110:39. doi: https://doi.org/10.1007/s00395-015-0496-3

3. Kamran P, Sereti KI, Zhao P, Ali SR, Weissman IL, Ardehali R. Parabiosis in mice: a detailed protocol. *J Vis Exp* 2013. doi: https://doi.org/10.3791/50556

4. Luo Y, Liu K, Ling D, Gu T, Zhang L, Chen W. Isolation and Characterization of Exosomes Derived from Mouse Spleen Tissues. *J Vis Exp* 2024. doi: https://doi.org/10.3791/67234

5. Zhang X, Wang C, Xu H, Cai S, Liu K, Li S, et al. Propofol inhibits myocardial injury induced by microvesicles derived from hypoxia-reoxygenated endothelial cells via lncCCT4-2/CCT4 signaling. *Biol Res* 2023;56:20. doi: https://doi.org/10.1186/s40659-023-00428-3

6. Chen L, Li S, Zhu J, You A, Huang X, Yi X, et al. Mangiferin prevents myocardial infarction-induced apoptosis and heart failure in mice by activating the Sirt1/FoxO3a pathway. *J Cell Mol Med* 2021;25:2944-2955. doi: https://doi.org/10.1111/jcmm.16329

7. Ge X, Meng Q, Wei L, Liu J, Li M, Liang X, et al. Myocardial ischemia-reperfusion induced cardiac extracellular vesicles harbour proinflammatory features and aggravate heart injury. *J Extracell Vesicles* 2021;10:e12072. doi: https://doi.org/10.1002/jev2.12072

8. Ge X, Meng Q, Zhuang R, Yuan D, Liu J, Lin F, et al. Circular RNA expression alterations in extracellular vesicles isolated from murine heart post ischemia/reperfusion injury. *Int J Cardiol* 2019;296:136-140. doi: https://doi.org/10.1016/j.ijcard.2019.08.024

9. Zuo R, Ye LF, Huang Y, Song ZQ, Wang L, Zhi H, et al. Hepatic small extracellular vesicles promote

microvascular endothelial hyperpermeability during NAFLD via novel-miRNA-7. *J Nanobiotechnology* 2021;19:396. doi: https://doi.org/10.1186/s12951-021-01137-3

10. Lasser C, Kishino Y, Park KS, Shelke GV, Karimi N, Suzuki S, et al. Immune-Associated Proteins Are Enriched in Lung Tissue-Derived Extracellular Vesicles during Allergen-Induced Eosinophilic Airway Inflammation. *Int J Mol Sci* 2021;22. doi: https://doi.org/10.3390/ijms22094718

11. Crescitelli R, Lasser C, Lotvall J. Isolation and characterization of extracellular vesicle subpopulations from tissues. *Nat Protoc* 2021;16:1548-1580. doi: https://doi.org/10.1038/s41596-020-00466-1

12. Jingushi K, Uemura M, Ohnishi N, Nakata W, Fujita K, Naito T, et al. Extracellular vesicles isolated from human renal cell carcinoma tissues disrupt vascular endothelial cell morphology via azurocidin. *Int J Cancer* 2018;142:607-617. doi: https://doi.org/10.1002/ijc.31080

13. Zielonka J, Kalyanaraman B. Hydroethidine- and MitoSOX-derived red fluorescence is not a reliable indicator of intracellular superoxide formation: another inconvenient truth. *Free Radic Biol Med* 2010;48:983-1001. doi: https://doi.org/10.1016/j.freeradbiomed.2010.01.028

14. Mol EA, Lei Z, Roefs MT, Bakker MH, Goumans MJ, Doevendans PA, et al. Injectable Supramolecular Ureidopyrimidinone Hydrogels Provide Sustained Release of Extracellular Vesicle Therapeutics. *Adv Healthc Mater* 2019;8:e1900847. doi: https://doi.org/10.1002/adhm.201900847

15. Wisniewski JR, Zougman A, Nagaraj N, Mann M. Universal sample preparation method for proteome analysis. *Nat Methods* 2009;6:359-362. doi: https://doi.org/10.1038/nmeth.1322

16. Hoshino A, Kim HS, Bojmar L, Gyan KE, Cioffi M, Hernandez J, et al. Extracellular Vesicle and Particle Biomarkers Define Multiple Human Cancers. *Cell* 2020;182:1044-1061 e1018. doi: https://doi.org/10.1016/j.cell.2020.07.009

17. Lacroix R, Judicone C, Poncelet P, Robert S, Arnaud L, Sampol J, et al. Impact of pre-analytical parameters on the measurement of circulating microparticles: towards standardization of protocol. *J*

*Thromb Haemost* 2012;10:437-446. doi: https://doi.org/10.1111/j.1538-7836.2011.04610.x
